# Natural compound screening predicts novel GSK-3 isoform-specific inhibitors

**DOI:** 10.1101/2024.04.22.590490

**Authors:** Firdos Ahmad, Anamika Gupta, Hezlin Marzook, James R. Woodgett, Mohamed A. Saleh, Rizwan Qaisar

## Abstract

Glycogen synthase kinase-3 (GSK-3) plays important roles in the pathogenesis of cardiovascular, metabolic, neurological disorders and cancer. Isoform-specific loss of either GSK-3α or GSK-3β often provides cytoprotective effects under such clinical conditions. However, available synthetic small molecule inhibitors are relatively non-specific, and their chronic use may lead to adverse effects. Therefore, screening for natural compound inhibitors to identify the isoform-specific inhibitors may provide improved clinical utility. Here, we screened 70 natural compounds to identify novel natural GSK-3 inhibitors employing comprehensive *in silico* and biochemical approaches. Molecular docking and pharmacokinetics analysis identified two natural compounds Psoralidin and Rosmarinic acid as potential GSK-3 inhibitors. Specifically, Psoralidin and Rosmarinic acid exhibited the highest binding affinities for GSK-3α and GSK-3β, respectively. Consistent with *in silico* findings, the kinase assay-driven IC50 revealed superior inhibitory effects of Psoralidin against GSK-3α (IC50=2.26 µM) vs. GSK-3β (IC50=4.23 µM) while Rosmarinic acid was found to be more potent against GSK-3β (IC50=2.24 µM) than GSK-3α (IC50=5.14 µM). Taken together, these studies show that the identified natural compounds may serve as GSK-3 inhibitors with Psoralidin serving as a better inhibitor for GSK-3α and Rosmarinic for GSK-3β isoform, respectively. Further characterization employing *in vitro* and preclinical models will be required to test the utility of these compounds as GSK-3 inhibitors for cardiometabolic and neurological disorders and cancers.

**Highlights:**

- Current GSK-3 inhibitors lack specificity and cause side effects.
- This study identifies potential GSK-3 isoform-specific natural compounds.
- Psoralidin is likely a better inhibitor for GSK-3α while Rosmarinic for GSK-3β.
- These natural compounds may be promising future treatments.

## 1. Introduction

Glycogen synthase kinase-3 (GSK-3) serves as a critical connection in complex cellular signaling pathways [1]. It is implicated in a diverse range of cellular processes ranging from cell growth and proliferation to cell death and metabolic control [1-4]. Aberrant activity of GSK-3 leads to dysregulation of cellular processes that underscore a variety of clinical conditions such as diabetes, inflammation, cancer, cardiovascular diseases and neurodegeneration [5]. Hence, targeting GSK-3 isoforms has emerged as a promising therapeutic strategy with potential applications across diverse medical conditions.

GSK-3, which comprises two highly related kinases, GSK-3α and GSK-3β encoded by distinct genes, modulates numerous downstream substrates to regulate a variety of cellular functions [6]. Unlike most protein kinases, GSK-3 is active in resting cells and is inhibited by a variety of selected cellular stimuli [4]. In healthy states, GSK-3 plays crucial roles in maintaining cellular homeostasis, regulating glycogen metabolism, cell cycle progression, and cell death [7, 8]. GSK-3α and GSK-3β exhibit both overlapping and non-redundant distinctive functions [9, 10] but their high similarity (∼97% identical residues in the catalytic domain) renders it difficult to target each selectively [11, 12]. The activities of GSK-3α and β isoforms can be regulated through phosphorylation at Ser21 and Ser9 and one isoform cannot completely compensate for the loss of the other. For example, germline global deletion of GSK-3β causes embryonic lethality [13] while germline global loss of GSK-3α results in viable animals [14] of which the males are sterile and both sexes later develop early aging and age-related complications [15]. Moreover, cardiomyocyte-specific conditional loss of both GSK-3α and GSK-3β isoforms in adult cardiomyocytes leads to mitotic catastrophe and fatal dilated cardiomyopathy [16, 17]. Interestingly, isoform-specific targeting of GSK-3 is found to be protective in a variety of clinical conditions including cancer, neurodegenerative and cardiometabolic disorders [1, 18, 19].

In conjunction with recent discoveries on therapeutic importance and new functions of GSK-3, many additional small molecular inhibitors of GSK-3 have been synthesized and characterized [20]. While synthetic GSK-3 small molecule inhibitors have shown therapeutic efficacy in preclinical models, their side effects and limited selectivity have impeded clinical translation. Most inhibitors are less specific due to their mechanism of action as reversible ATP competitors [21]. This provides an avenue for screening nature-derived bioactive molecules. These nature-derived compounds represent a promising approach to treat various conditions as they are often present in our diet or can be added as supplements. Furthermore, natural compounds often exhibit better safety profiles and favorable pharmacokinetic properties, compared to synthetic molecules. For example, natural phytoconstituents are attracting interest in use as protein kinase inhibitors and show promise against a variety of kinases [22]. Natural products like curcumin, berberine, Rosmarinic acid, resveratrol, apocynin, caffeine, indirubin, tetrandrine, discorea, wogonin, and gambogenic acid demonstrate modest inhibition of GSK-3 isoforms [23]. Other natural compounds such as Psoralidin, a natural phenolic coumarin, show antioxidative and anti-inflammatory activities and protection against cancer and osteoporosis [24, 25]. Psoralidin induces cellular apoptosis and inhibits proliferation and promotes autophagy-mediated cell death to show the protective effects in cancer [26]. Moreover, Psoralidin induces oxidative stress and activates the estrogen receptors (ER)-signaling pathway in cancerous cell. Further studies have shown that Psoralidin has activity against colon cancer by inhibiting NF-κB and Bcl-2/Bax signaling pathways [27]. Similarly, another antioxidant compound, Rosmarinic acid, a potential GSK-3 inhibitor, displays protective roles in inflammatory and neurodegenerative diseases [28]. These natural compounds appear to have effects in several pathological conditions however, it is largely undefined whether they target GSK-3 and if so, which isoform of GSK-3.

The most challenging aspect of natural compound inhibitors is their high molecular weight, poor stability and weak solubility. Therefore, computational investigation can assist in focusing on inhibitors with more selective characteristics, and potency assessed in their derivatives. Employing such *in silico* approaches, natural cellular pathways including the PI3K-Akt pathway have emerged as a key target of natural products [29]. Downstream of this important signaling pathway, GSK-3 could be a prominent target of natural compounds to show the protective effects in many critical diseases including cancer, neurodegenerative and cardiovascular complications.

Here, we have sought to initially identify natural GSK-3 inhibitors by screening 70 natural compounds employing computational approaches. Among several potential GSK-3 inhibitors, Rosmarinic acid showed highest binding for GSK-3β vs GSK-3α. In contrast, Psoralidin displayed highest binding toward GSK-3α though this compound also bound GSK-3β. Kinase activity assessment revealed the potency of these compounds against GSK-3 isoforms where Psoralidin was found to be more potent against GSK-3α and Rosmarinic acid more specific against GSK-3β.

## 2. Methods

### 2.1 Ligand Preparation

We retrieved the 3D structures of 70 natural phytoconstituents and 5 synthetic reference drugs from freely available chemical structure database, PubChem (https://pubchem.ncbi.nlm.nih.gov/) in SDF format. Before subjecting to docking, energy minimization was carried out of all ligands using Merck Molecular Force Field (MMFF94) as described previously [30, 31]. Structural optimization of all ligands was done using AutoDock Tools (ADT) version 4.2.6. The PubChem ID of all phytoconstituents and reference synthetic drugs are provided as supplementary materials **(Suppl. table 1A-D)**.

### 2.2 Target Protein Source

The 3D crystal structure of the target protein GSK-3β (based on X-ray diffraction structure available in the RCSB database) was downloaded from the Protein Data Bank (PDB) available at http://www.rcsb.org/pdb. Refinements and energy minimization of GSK-3β (PDB ID. 1H8F) were carried out before subjecting it to molecular docking analysis and structures were visualized by Accelrys Biovia Discovery Studio 2017 R2 (Biovia, San Diego, CA, USA) as described previously [30]. Protein refinement was carried out by inserting polar hydrogen atoms and Kollman charges, removing crystallographic water molecules as well as removing normal and unnecessary protein ligands and ions.

### 2.3 Homology modeling of GSK-3α

Since the crystallographic structure of GSK-3α is not available in the RCSB database, homology modeling was employed using the online server ProMod3 Version3.2.0 SWISS-MODEL (http://swissmodel.expasy.org) as described previously [32]. The amino acid sequences of human GSK-3α were obtained from the universal protein knowledge database (UniProtKB) with ID P49840 (GSK3A_HUMAN).

To run sequence alignment, the sequences of GSK-3α were retrieved from different organisms in FASTA format using Expert Protein Analysis System, or ExPASy Molecular Biology Server and additional sequences were selected from distally related organisms, to compare the conserved amino acids sequences as described previously [31]. The sequences of GSK-3α were subjected to BLAST (Basic Local Alignment Search Tool) against the sequences of selected known protein structures from the ExPDB, the Swiss Model template library. Generally, a percentage sequence identity above 50% was selected as a relatively candid modeling project, while anything below required more careful planning. Further, 3D atomic models of the query proteins were generated using ProMod3 *version* 1.0.2.

### 2.4 Homology model validation

The quality of the GSK-3α modeled protein structure was validated by generating a Ramachandran plot using the Molprobity web server which is a structure-validation web service that evaluates the quality of homology-modeled protein structures and nucleic acid (http://molprobity.biochem.duke.edu) [33]. For visualization purposes, Accelrys Biovia Discovery Studio 2017 R2 (Biovia, San Diego, CA, USA) was used [30]. AutoDock Tools version 4.2.6 was used for the preparation of the protein model of GSK-3α where all existing water or solvent molecules were removed to terminate the impact the solvent interactions protein-ligand docking.

### 2.5 Molecular Docking

The optimized proteins and ligand structures were subjected to molecular docking analysis using AutoDock *version* 4.2.6, (Molecular Graphics Lab, Scripps Research Institute, La Jolla, CA 92037, USA http://autodock.scripps.edu/). pdbqt files were prepared against GSK-3α and GSK-3β. The binding sites were determined by a grid box with XYZ dimensions of 60 Å × 60 Å × 60 Å around the active site of the proteins and a spacing value of 0.375 Å. The best docking orientations were analyzed using Accelrys Biovia Discovery Studio 2017 R2 (Biovia, San Diego, CA, USA). The docking software calculates the binding energies (kcal/mol) and dissociation constant (K_d_) values of potential ligands to GSK-3α and β isoforms. Lower binding energies and *K*_*d*_ values indicate greater affinity of the phytoconstituent towards a particular target protein

### 2.6 Prediction of Activity Spectra for Substances analysis

Prediction of Activity Spectra for Substances (PASS), is an automated platform that detects biological activity of chemical compounds. PASS analysis was performed using OSIRIS Property Explorer *version* 4.5.1 (http://www.openmolecules.org/propertyexplorer/index.html) to predict biological activities of chemical compounds considering molecular weight, number of hydrogen bond acceptors and donors, rotatable bond [30].

### 2.7 Drug likeness

Drug-like behavior of selected phytoconstituents and synthetic reference drugs were evaluated using the Molinspiration web server (https://www.molinspiration.com/) which follows Lipinski’s rule of five [34]. The oral activity of potential drug-like quality in the selected compounds/drugs must obey Lipinski’s rule of five (RO5) parameters such as ≤5 H-bond donors, ≤10 H-bond acceptors, ≤500 molecular weight, ≤140 Å^2^ topological polar surface area (TPSA), and ≤ calculated Log P (CLogP), that would make it potential drug for humans. In general, an orally active potential drug must outline the molecular property of drug-like pharmacokinetics in the human body which includes their absorption, distribution, metabolism, excretion and toxicity (ADMET). The violation of one or more of the mentioned parameters predicts a molecule as a non-orally available drug [35]. Further, the mentioned phytoconstituents and drugs were evaluated for obeying some set of rules of Veber (GSK) rule [36], Ghose (Amgen) filter [37], Egan filter [38] and Muegge (Bayer) filters [39] for druglikeness by user-friendly interface SwissADME (http://www.swissadme.ch/).

### 2.8 Bioactivity score prediction

The bioactivity scores including drug likeness scores towards G-protein coupled receptor (GPCR) ligands, ion channel modulators, kinase inhibitors and nuclear receptor ligands were evaluated against selected phytoconstituents and reference drugs using Molinspiration. Drug likeness score values help determine a compound’s overall medical potential as a drug candidate. Compounds having a bioactivity score >0.0 are considered as biologically active; score values between −5.0 and 0.0 are considered as moderately biologically active and score values <⍰−5.0 are perceived as having poor biological activity [35].

### 2.9 Toxicity potential assessment

The toxicity prediction of a potential drug is crucial and a preliminary step in drug discovery and development that gives an idea about the potential side effects of a drug-like compound. The toxic parameters such as mutagenic, tumorigenic, irritant, and reproductive toxicities of included phytoconstituents and reference drugs were assessed using OSIRIS Data Warrior Software (http://www.openmolecules.org/datawarrior/) [35].

### 2.10 Pharmacokinetics Properties

The properties such as Absorption, Distribution, Metabolism, Excretion and Toxicity (ADMET) of prospective therapeutic drugs must be predicted early in the drug discovery process to avoid drug failure in the clinical phase. ADMET properties of selected phytoconstituents and reference drugs were evaluated using SwissADME software that assessed lipophilicity and polarity of target compounds/drugs through parameters such as blood-brain barrier, gastrointestinal absorption, Cytochromes P450 isoenzymes (CYPs) and skin permeability [40].

### 2.11 Swiss target prediction analysis

Bioactive small molecules bind to proteins or other macro-molecular targets and modulate their bioactivity. Swiss Target Prediction software (http://www.swisstargetprediction.ch) is an interface that provides molecular insight into bioactive molecules similar to biochemical targets such as kinase, enzyme, hydrolase, phosphatase, transferase, nuclear receptor, oxidoreductase, G-protein coupled receptors, transcription factors ion channels, cytochrome P450, cytosolic protein, protease and enzyme inhibitor. In the present study, Swiss Target Prediction was used to predict protein targets of potential drug-like candidates as described previously [40].

### 2.12 Bioavailability Radar plot

SwissADME tool was used to predict bioavailability radar following six physicochemical properties including lipophilicity, size, polarity, solubility, flexibility and saturation to reside within the pink zone to be considered as a potential drug candidate as described previously [41]. Briefly, the pink zone represents an optimum range of each parameter such as lipophilicity: XLOGP3 between −0.7 to +5.0 range, size: molecular weight (MW) between 150 to 500 g/mol range, polarity: TPSA between 20 to 130 Å^2^ range, insolubility: log S(ESOL) between 0 to −6 range, unsaturation: fraction of Csp3 between 1 to 0.25 range, and flexibility: number of rotatable bonds between 9 to 0 range [42].

### 2.13 Kinase Assay

Kinase assay for GSK-3α and GSK-3β to assess the inhibitory potency of Psoralidin and Rosmarinic acid was performed employing GSK-3α and GSK-3β enzyme system (Promega Cat# V9361 and V9371, respectively) following the manufacturer’s instructions. Briefly, GSK-3α and GSK-3β kinases were diluted with 5X reaction buffer containing 200µM DTT to reach final varying concentrations of 2, 4, 6, 8 and 10nM. An ATP/substrate mixture was prepared using of 5X reaction buffer. The inhibitors were used in a range of concentrations as indicated. The reactions were performed in 384-well white flat bottom plates. To each well, inhibitor, enzyme solution and ATP/substrate mix were added making up the volume of 5μL/well. The reaction standards were also kept as per the manufacturer’s instructions. A total of 5μL/well ADP-Glo reagent was added and incubated for 40 minutes followed by the kinase detection reagent (10 μL/well) at room temperature and allowed to equilibrate for 1 hour for ADP-ATP conversion. The luminescence intensity was recorded by GLOMax plate reader (Promega #GM3000) at 1 sec compensation. The relative light units (RLU) were normalized against 100% enzyme activity and, a dose-response curve was generated. The percent inhibition was calculated using non-linear curve fitting of log (inhibitor) against normalized response using GraphPad Prism software.

## 3. Results

### 3.1 Generation of GSK-3α homology protein structure model

Since 3D-crystallographic structure of GSK-3α protein is not available, first we generated the GSK-3α homology protein structure model. A BLAST search via SWISS-MODEL displayed multiple templates to match the target sequence of GSK-3α. Template 3say.1.A was predicted to have the highest sequence identity to the query GSK-3α protein sequence which was 82.97% as shown in **Suppl. Fig. 1**. The query GSK-3α protein contained 442 amino acids. The reasonably best template was selected based on the sequence identity, sequence similarity, query coverage and the availability of a 3D structure **(Fig. 1A)**. The MolProbity score of the generated GSK-3α model was determined as 1.56 which was well defined within the allowed region **(Suppl. Fig. 2)**. The modeled structure was validated using Ramachandran plots that highlight the most favored, allowed and disallowed regions of the modeled protein structure. Ramachandran plots of the GSK-3α model revealed that 95.7% of residues fall inside the favored 98% region **(Fig. 1B, Suppl. Fig. 3)** and there were two out of 348 Ramachandran outliers (Pro 321 and Asn 416), indicating the predicted GSK-3α model was of reasonably high quality. The 3D structure for GSK-3β was acquired from the protein database **(Fig. 1C)**.

**Figure 1:**
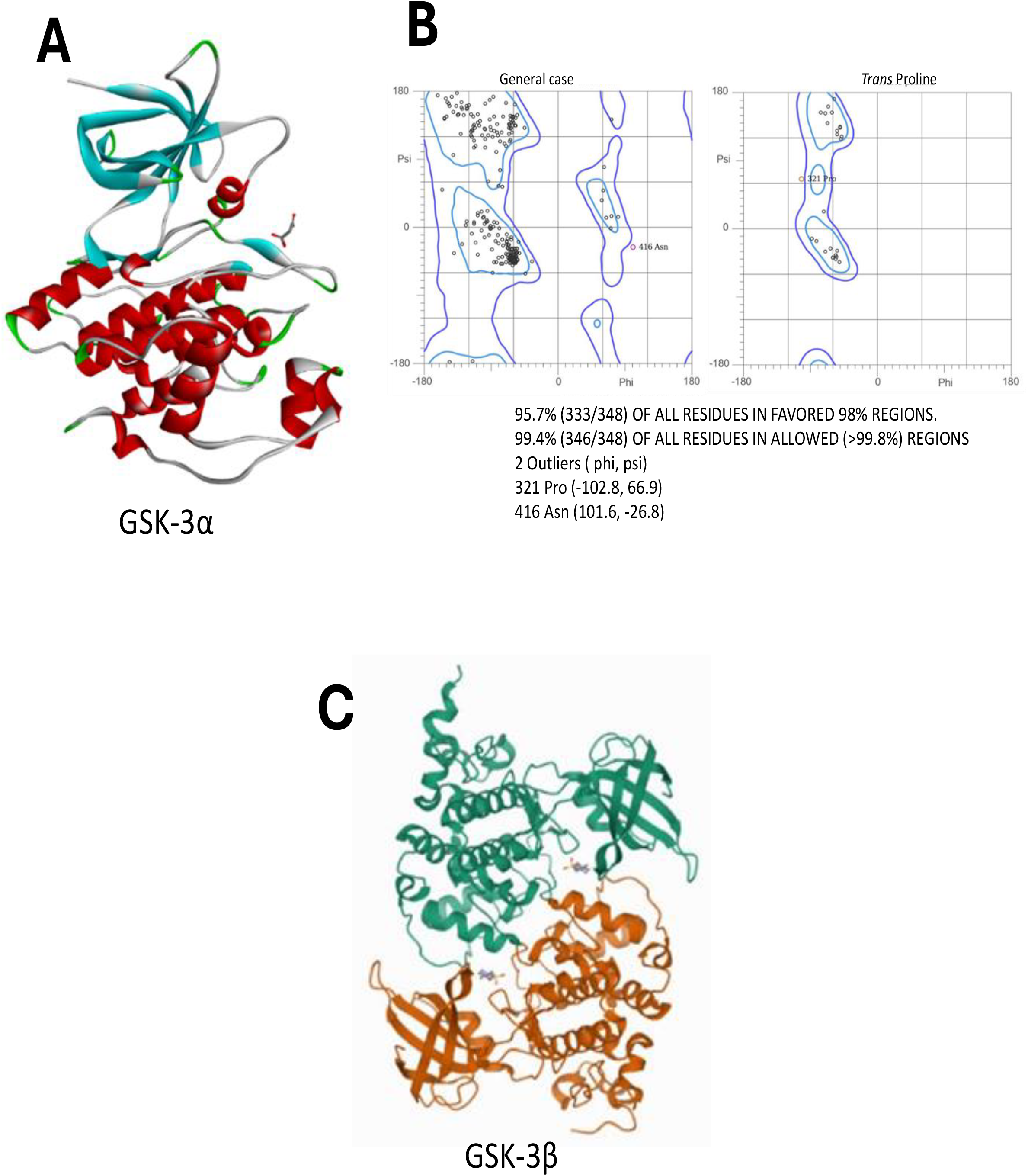
Molecular Modeling of GSK-3α. **(A)** A 3D structure of GSK-3α protein model and **(B)** The Ramachandran plot of GSK-3α model structure. (C) The crystal structure of GSK-3β was acquired from the protein databank.

### 3.2 Screening of natural compounds to identify the novel GSK-3 inhibitors

Small compounds (ligands) that fit well into the predicted binding pocket of a selected target protein can be identified using molecular docking. To associate GSK-3 with natural compounds, 70 naturally derived ligands were short-listed from publicly available datasets and publications. Six small molecular inhibitors with previously known inhibitory effects on GSK-3α or GSK-3β were included as positive controls in the analysis. Docking was carried out for the 70 phytoconstituents **(Suppl. Table 1A)** using AutoDock *version* 4.2.6 and the six-reference synthetic small molecular GSK-3 inhibitors **(Suppl. Table 1B)**. Of the analyzed natural compounds, 40 were calculated to have significant binding energies and dissociation constants against either GSK-3α and/or GSK-3β **(Table 1A, Suppl. Tables 1C-D)** respectively. The comparative display of binding energies and dissociation constants against GSK-3α and GSK-3β revealed that some of the selected phytoconstituents had significantly higher predicted binding affinity for GSK-3β compared to GSK-3α **(Tables 1A-B)**. In contrast, a few phytoconstituents including Berberine, Rosmarinic acid, Luteolin, Apigenin, Daidzein, Galangin, Psoralidin, Catechin, Crocetin, Piperine, Sesamin, Quercetin, Resveratrol, Kaempferol, Costunolide, Withaferin A, and Serpentine were predicted to display higher affinity towards GSK-3α compared to GSK-3β **(Table 1A)**.

**Table 1A;.**
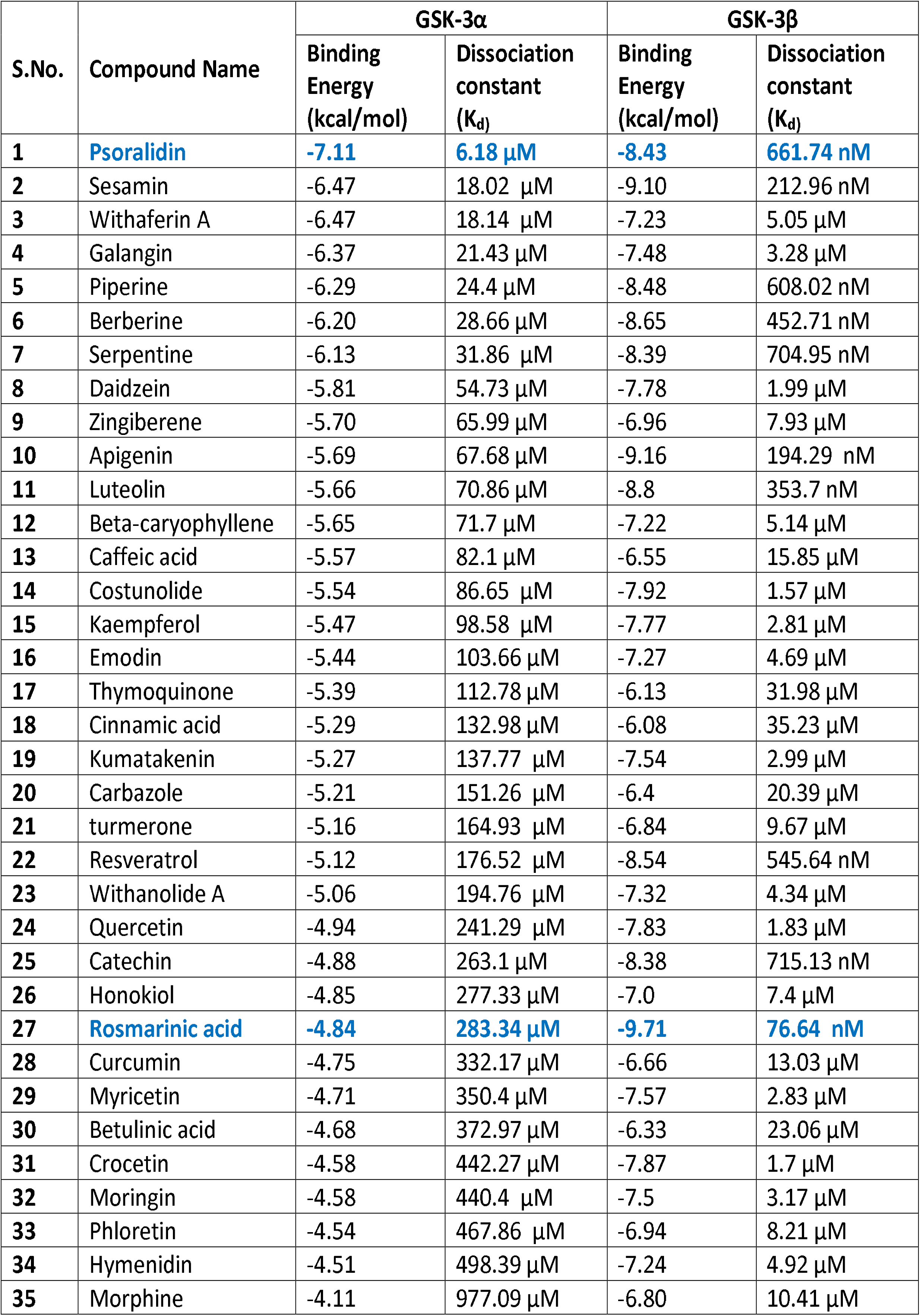
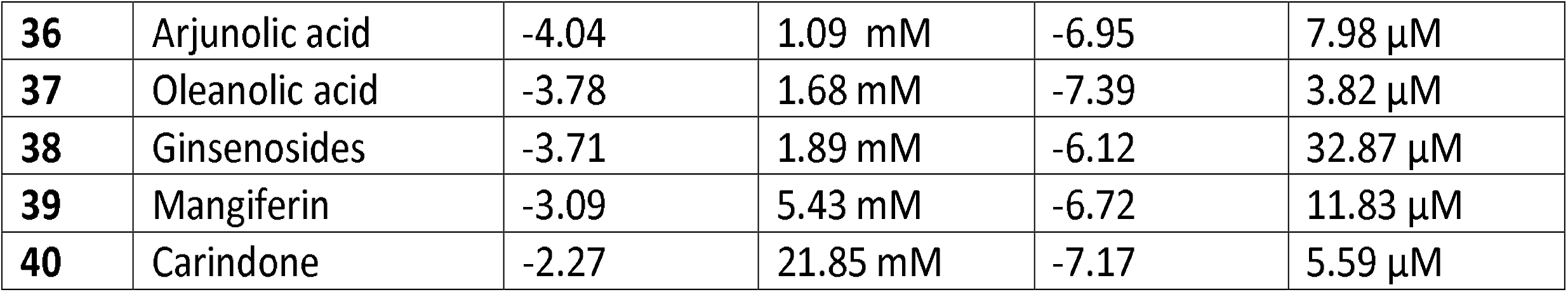
Computational comparative binding energies (kcal/mol) and dissociation constants (Kd) towards GSK-3α and GSK-3β against selected natural compounds using AutoDock version 4.2.6.

| S.No. | Compound Name | GSK-3α | | GSK-3β | |
| --- | --- | --- | --- | --- | --- |
|  |  | Binding Energy (kcal/mol) | Dissociation constant (K<sub>d</sub>) | Binding Energy (kcal/mol) | Dissociation constant (K<sub>d</sub>) |
| 1 | Psoralidin | -7.11 | 6.18 μM | -8.43 | 661.74 nM |
| 2 | Sesamin | -6.47 | 18.02 μM | -9.10 | 212.96 nM |
| 3 | Withaferin A | -6.47 | 18.14 μM | -7.23 | 5.05 μM |
| 4 | Galangin | -6.37 | 21.43 μM | -7.48 | 3.28 μM |
| 5 | Piperine | -6.29 | 24.4 μM | -8.48 | 608.02 nM |
| 6 | Berberine | -6.20 | 28.66 μM | -8.65 | 452.71 nM |
| 7 | Serpentine | -6.13 | 31.86 μM | -8.39 | 704.95 nM |
| 8 | Daidzein | -5.81 | 54.73 μM | -7.78 | 1.99 μM |
| 9 | Zingiberene | -5.70 | 65.99 μM | -6.96 | 7.93 μM |
| 10 | Apigenin | -5.69 | 67.68 μM | -9.16 | 194.29 nM |
| 11 | Luteolin | -5.66 | 70.86 μM | -8.8 | 353.7 nM |
| 12 | Beta-caryophyllene | -5.65 | 71.7 μM | -7.22 | 5.14 μM |
| 13 | Caffeic acid | -5.57 | 82.1 μM | -6.55 | 15.85 μM |
| 14 | Costunolide | -5.54 | 86.65 μM | -7.92 | 1.57 μM |
| 15 | Kaempferol | -5.47 | 98.58 μM | -7.77 | 2.81 μM |
| 16 | Emodin | -5.44 | 103.66 μM | -7.27 | 4.69 μM |
| 17 | Thymoquinone | -5.39 | 112.78 μM | -6.13 | 31.98 μM |
| 18 | Cinnamic acid | -5.29 | 132.98 μM | -6.08 | 35.23 μM |
| 19 | Kumatakenin | -5.27 | 137.77 μM | -7.54 | 2.99 μM |
| 20 | Carbazole | -5.21 | 151.26 μM | -6.4 | 20.39 μM |
| 21 | turmerone | -5.16 | 164.93 μM | -6.84 | 9.67 μM |
| 22 | Resveratrol | -5.12 | 176.52 μM | -8.54 | 545.64 nM |
| 23 | Withanolide A | -5.06 | 194.76 μM | -7.32 | 4.34 μM |
| 24 | Quercetin | -4.94 | 241.29 μM | -7.83 | 1.83 μM |
| 25 | Catechin | -4.88 | 263.1 μM | -8.38 | 715.13 nM |
| 26 | Honokiol | -4.85 | 277.33 μM | -7.0 | 7.4 μM |
| 27 | Rosmarinic acid | -4.84 | 283.34 μM | -9.71 | 76.64 nM |
| 28 | Curcumin | -4.75 | 332.17 μM | -6.66 | 13.03 μM |
| 29 | Myricetin | -4.71 | 350.4 μM | -7.57 | 2.83 μM |
| 30 | Betulinic acid | -4.68 | 372.97 μM | -6.33 | 23.06 μM |
| 31 | Crocetin | -4.58 | 442.27 μM | -7.87 | 1.7 μM |
| 32 | Moringin | -4.58 | 440.4 μM | -7.5 | 3.17 μM |
| 33 | Phloretin | -4.54 | 467.86 μM | -6.94 | 8.21 μM |
| 34 | Hymenidin | -4.51 | 498.39 μM | -7.24 | 4.92 μM |
| 35 | Morphine | -4.11 | 977.09 μM | -6.80 | 10.41 μM |
| **36** | Arjunolic acid | -4.04 | 1.09 mM | -6.95 | 7.98 μM |
| **37** | Oleanolic acid | -3.78 | 1.68 mM | -7.39 | 3.82 μM |
| **38** | Ginsenosides | -3.71 | 1.89 mM | -6.12 | 32.87 μM |
| **39** | Mangiferin | -3.09 | 5.43 mM | -6.72 | 11.83 μM |
| **40** | Carindone | -2.27 | 21.85 mM | -7.17 | 5.59 μM |

**Table 1B;.**
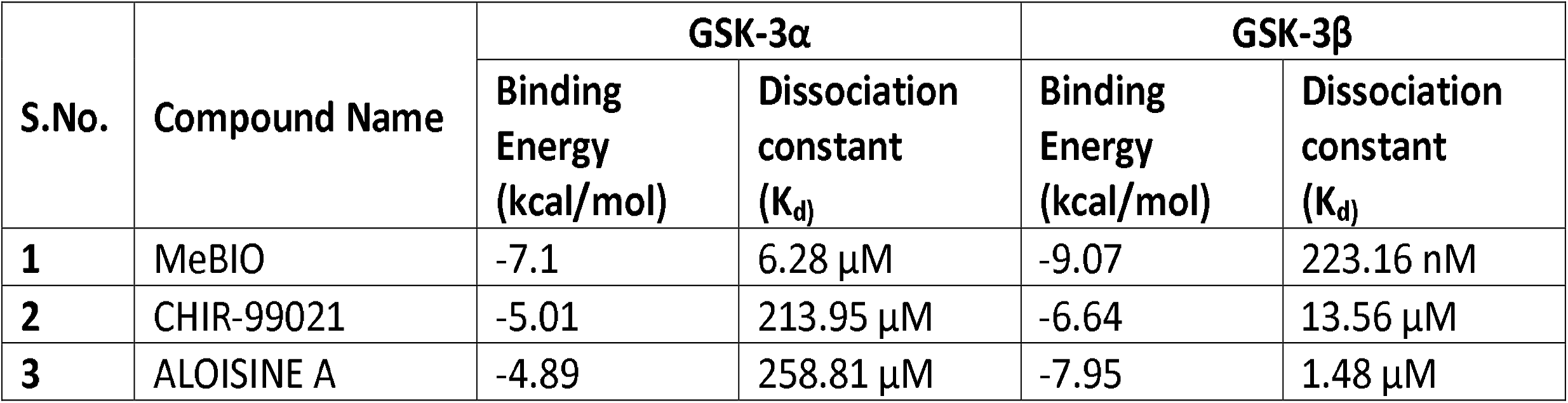
Computational comparative binding energies (kcal/mol) and dissociation constants (Kd) towards GSK-3α and GSK-3β against selected reference synthetic drugs using AutoDock version 4.2.6.

Considering the GSK-3α modeled protein structure, few phytoconstituents **(Table 1A, Suppl. Fig. 4)** displayed binding affinity in a descending chronology from Psoralidin, Sesamin, Withaferin A, Galangin, Piperine, Berberin, Serpentine, Daidzein, Zingiberene, and Apigenin. Psoralidin was predicted to have the highest binding affinity of −7.11 (kcal/mol), dissociation constant (6.18 µM) for GSK-3α among selected ligands. The docking pose of Psoralidin with GSK-3α is modeled in **Fig. 2A**. For GSK-3β, selected phytoconstituents showed the following trend of decreasing binding affinity with the highest ranging from Rosmarinic acid, Apigenin, Sesamin, Luteolin, Berberine, Resveratrol, Piperine, Psoralidin, Serpentine, Catechin, Costunolide, Crocetin, Quercetin, Daidzein, and Kaempferol being the least in the predicted list (Table 1A). Among the mentioned phytoconstituents Rosmarinic acid, Apigenin and Sesamin hasd a calculated highest binding energy of −9.71 (kcal/mol); K_d_ (76.64 nM), −9.16 (kcal/mol); K_d_ (194.29 nM) and −9.10 (kcal/mol); K_d_(212.96 nM) respectively against GSK-3β. The best docking pose of Rosmarinic acid with GSK-3β crystal structure was generated and is shown in **Fig. 2B**. Similar to these compounds, the reference synthetic small molecule inhibitors MeBIO, ALOISINE A and CHIR-99021 showed higher binding energy of (−7.1, −4.89 and −5.01 kcal/mol) against GSK-3α and (−9.07, −7.95 and −6.64 kcal/mol) against GSK-3β respectively **(Table 1B)**.

**Figure 2:**
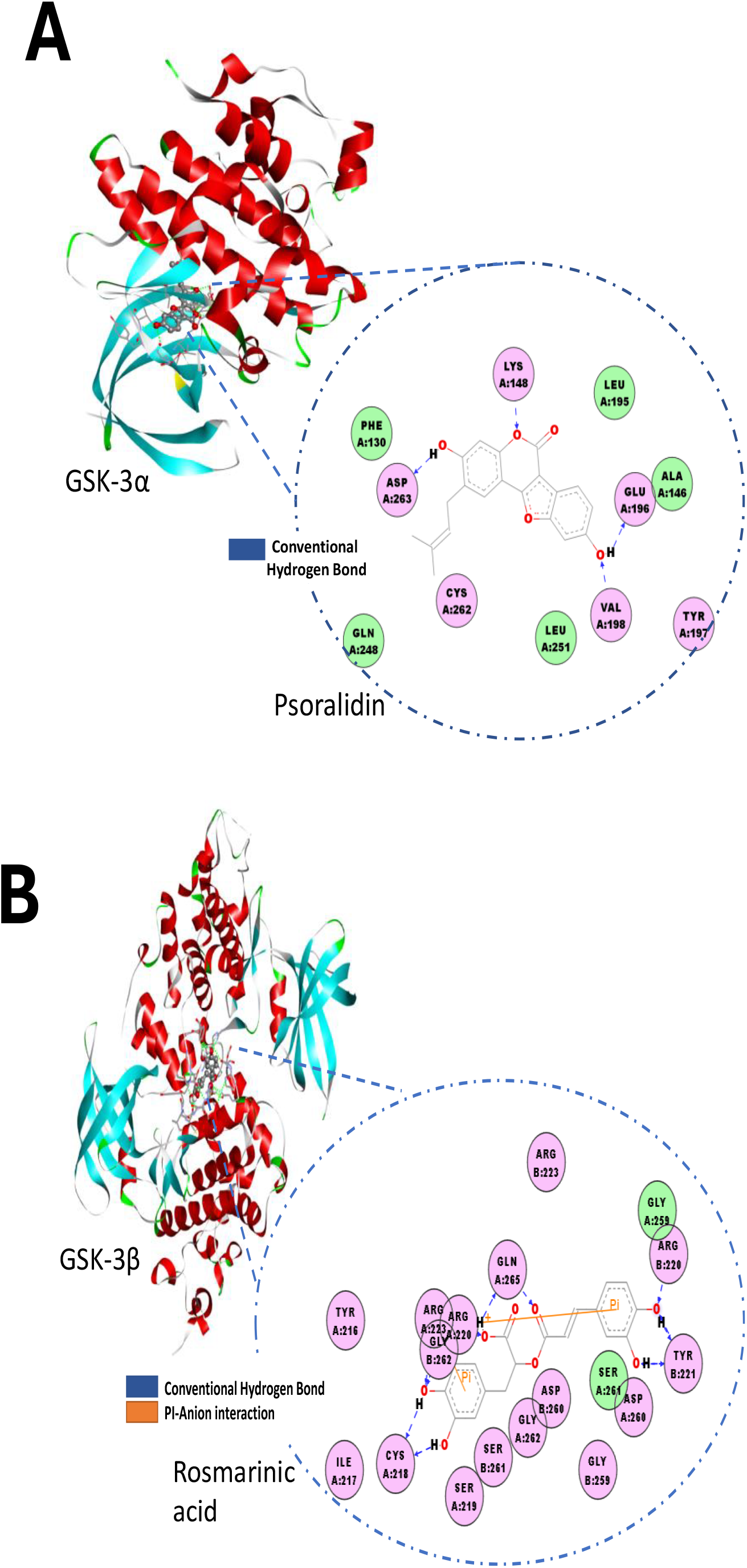
Molecular docking to identify the interaction between protein and ligands. **(A)** The molecular modeling shows the best docking pose and interacting amino acid of GSK-3α with Psoralidin in 3D and 2D views. **(B)** The molecular modeling shows the best docking pose and interacting amino acid of GSK-3β with Rosmarinic acid in 3D and 2D views.

The best docking poses and interacting amino acids of GSK-3α and GSK-3β with phytoconstituents and reference drugs were generated and summarized in **Suppl. Tables 1C-D**. However, to narrow down our study for further phytochemical screening, 17 phytoconstituents and three reference drugs were selected based on their binding affinity against target proteins **(Suppl. Fig. 4)**.

### 3.3 Pharmacological properties of included natural compounds

An orally active drug candidate must obey “Lipinski’s rule of five” without violating one or more parameters. The molecular properties including clogP, molecular weight, drug-likeness, topological polar surface area (TPSA), percentage absorption, Lipinski’s violations, hydrogen bond donor and acceptor of selected phytoconstituents and reference drugs are summarized in **Table 2**. In the present study, selected phytoconstituents were found to obey Lipinski’s parameters with no violation. Only Crocetin had clogP greater than 5.0. The included reference drugs displayed no violations of Lipinski’s rule of five except azithromycin which had two violations. The results obtained indicated that selected natural compounds followed all physiochemical properties to behave as potential drug candidates. Based on the mentioned parameters, the most effective identified natural compounds against GSK-3α and GSK-3β were Psoralidin and Rosmarinic acid.

**Table 2.** PASS analysis of selected natural compounds versus reference synthetic drugs (MeBIO, CHIR-99021, ALOISINE A and Azithromycin) calculated by OSIRIS Property Explorer.

| S. No. | Ligand | %Absorption (>50%)<sup>a</sup> | Topological polar surface area (Å)<sup>2</sup> (TPSA)<sup>b</sup> (<160 Å) | MW (<500) | c log P (<5)<sup>c</sup> | Heavy atom count | Hydrogen bond donors (nOHNH) (≤5) | Hydrogen bond acceptors (nON) (≤10) | Number of rotatable bonds (≤10) | Lipinski's violation |
| --- | --- | --- | --- | --- | --- | --- | --- | --- | --- | --- |
| 1 | Berberine | 94.92 | 40.82 | 336.37 | 0.5223 | 25 | 0 | 5 | 2 | 0 |
| 2 | Rosmarinic acid | 59.14 | 144.52 | 360.32 | 1.4502 | 26 | 5 | 8 | 7 | 0 |
| 3 | Luteolin | 70.66 | 111.12 | 286.24 | 1.99 | 21 | 4 | 6 | 1 | 0 |
| 4 | Apigenin | 77.64 | 90.89 | 270.24 | 2.3357 | 20 | 3 | 5 | 1 | 0 |
| 5 | Daidzein | 84.62 | 70.67 | 254.24 | 1.9729 | 19 | 2 | 4 | 1 | 0 |
| 6 | Galangin | 77.64 | 90.89 | 270.24 | 2.1816 | 20 | 3 | 5 | 1 | 0 |
| 7 | Psoralidin | 80.09 | 83.81 | 336.34 | 4.8632 | 25 | 2 | 5 | 2 | 0 |
| 8 | Catechin | 70.92 | 110.37 | 290.27 | 1.5087 | 21 | 5 | 6 | 1 | 0 |
| 9 | Crocetin | 83.26 | 74.60 | 328.41 | 5.4464 | 24 | 2 | 4 | 8 | 0 |
| 10 | Piperine | 95.62 | 38.78 | 285.34 | 3.5951 | 21 | 0 | 4 | 3 | 0 |
| 11 | Sesamin | 89.89 | 55.40 | 354.36 | 3.2248 | 26 | 0 | 6 | 2 | 0 |
| 12 | Quercetin | 63.68 | 131.35 | 302.24 | 1.4902 | 22 | 5 | 7 | 1 | 0 |
| 13 | Resveratrol | 88.07 | 60.68 | 228.25 | 2.8295 | 17 | 3 | 3 | 2 | 0 |
| 14 | Kaempferol | 70.66 | 111.12 | 286.24 | 1.8359 | 21 | 4 | 6 | 1 | 0 |
| 15 | Costunolide | 99.93 | 26.30 | 232.32 | 4.1833 | 17 | 0 | 2 | 0 | 0 |
| 16 | Withaferin A | 75.76 | 96.36 | 470.61 | 2.4938 | 34 | 2 | 6 | 3 | 0 |
| 17 | Serpentine | 90.59 | 53.37 | 348.40 | 1.331 | 26 | 0 | 5 | 2 | 0 |
| 18 | MeBIO | 84.72 | 70.39 | 370.21 | 4.1221 | 23 | 2 | 5 | 2 | 0 |
| 19 | ALOISINE A | 87.68 | 61.80 | 267.33 | 3.1754 | 20 | 2 | 4 | 4 | 0 |
| 20 | CHIR-99021 | 69.26 | 115.20 | 465.35 | 3.6043 | 32 | 3 | 8 | 7 | 0 |
| 21 | Azithromycin | 46.87 | 180.09 | 749.00 | 1.6569 | 52 | 5 | 14 | 7 | 2 |

<sup>a</sup>Percentage Absorption was calculated as: % Absorption = 109-[0.345\*Topological Polar Surface Area].
<sup>b</sup>Topological polar surface area (defined as a sum of surfaces of polar atoms in a molecule).
<sup>c</sup>Logarithm of compound partition coefficient between n-octanol and water.

### 3.4 Bioactivity score prediction and druglikeness

Bioactivity score is an important measure to predict the potency and/or molecular activity of drug-like compounds in biological receptor targets such as binding to the G protein-coupled receptor (GPCR) and nuclear receptor, ion channel modulation, kinase inhibition, protease inhibition, and enzyme activity inhibition. The higher the bioactivity score, the more likely the molecule will be biologically active and possess pharmacological effects. If the BAS is greater than 0.0 then the potential target drug is considered physiologically active and if the BAS score is between −5.0 to 0.0 then the target drug is moderately active and if⍰−5.0 then the drug is inactive [35]. In the present study, most of the included phytoconstituents and reference drugs were found to be active or moderately active, as summarized in Suppl. Table 2. The *in silico* analysis predicted that the selected phytoconstituents including Psoralidin and Rosmarinic acid may serve as an inhibitor against GSK-3α and GSK-3β.

### 3.5 Toxicity Risk Assessment

Toxicity evaluation of potential drug-like compounds is essential during the drug discovery and development process. Here, the toxicity potential of selected phytoconstituents was evaluated against parameters such as mutagenic, tumorigenic, reproductive and irritant effects. The results of the toxicity prediction are color-coded and are represented in **Suppl. Table 3**. Red color indicates high risk, green color predicts drug-like behavior (no effect) and yellow color denotes moderate risk. We observed that Berberine, Rosmarinic acid, Luteolin, Catechin, Crocetin, Sesamin, Serpentine, ALOISINE A, CHIR-99021 and Azithromycin had no toxicity effects. Quercetin and MeBIO showed high mutagenic and tumorigenic effects. Whereas, Apigenin, Galangin, Resveratrol and Kaempferol showed a high mutagenic potential. Only Costunolide was found to have irritant properties. Only Costunolide was found to show an irritant effect. Daidzein, Psoralidin, Piperine, and Resveratrol were predicted to show high reproductive effects except for Withaferin A with low reproductive effects.

### 3.6 Assessment of pharmacokinetic properties

Several factors determine the pharmacokinetic properties of drugs. Compounds with higher MW are unlikely to be absorbed and digested in the GI tract; thereby low levels are detected in plasma. Our analysis predicted 59.14% absorption for Rosmarinic acid while it was 80.09% for Psoralidin **(Table 2)**. Another important parameter for characterizing drug efficacy is the blood-brain barrier (BBB) which allows highly selective permeability of drugs to the brain [43]. Berberine, Daidzein, Piperine, Sesamin, Resveratrol, Costunolide, and Serpentine were predicted to be capable of penetrating the BBB. The rest of the compounds were predicted to be incapable of penetrating the BBB membrane.

Drug transporter, P-glycoprotein (P-gp) is an ATP-dependent drug efflux pump that determines uptake and efflux of compounds [44]. Unlike Berberine, Catechin, Withaferin A, ALOISINE a, and Azithromycin, the rest of the compounds were predicted not to behave as P-gp substrates. As a result, these drugs are less likely to be pumped out of the cell by the glycoprotein. CYP1A2 is a member of the cytochrome P450 superfamily of enzymes that catalyzes many reactions involved in drug metabolism and oxidizes the organic molecules to eliminate them from circulation [45]. In the present study, Catechin, Crocetin, Rosmarinic acid, Resveratrol, Sesamin, Withaferin A, Costunolide, Serpentine, and Azithromycin were predicted not to inhibit CYP1A2, thus making them more likely to be metabolized by enzymes **(Suppl. Table 5)**. Moreover, CYP2D6 is also responsible for the metabolism and elimination of toxins from the body [46]. Rosmarinic acid, Psoralidin, Catechin, Crocetin, Piperine, and Costunolide were predicted not to inhibit CYP2D6 and CYP3A4 inhibitors. Hence, it can easily be metabolized and eliminated from the body, as compared to the rest of the compounds. CYP2C19 is another family of P450 liver enzymes and inhibiting it may lead to antiepileptic toxicity. Psoralidin, Crocetin, Piperine, Costunolide, Sesamin and Serpentine were predicted to inhibit CYP2C19 and CYP2C9 metabolizing enzymes **(Suppl. Table 5)**.

### 3.7 Mapping the targets and characterization of medicinal properties

Swiss Target Prediction, an online tool [47] was used to predict the similarity between bioactive molecules with identical biochemical targets. The frequency of protein targets such as kinases, oxidoreductases, phosphatases, proteases, phosphodiesterases and lyases, were predicted for selected phytoconstituents versus reference synthetic drugs as shown in the pie chart **(Fig. 3a-t)**. Our results show that all the phytoconstituents listed were found to be active toward several classes of soluble enzymes such as kinases, cytochrome p450, proteases, membrane-bound enzymes and transporter proteins. Thus, it is evident that these compounds possess broad-spectrum bioactivity against protein targets in humans.

**Figure 3:**
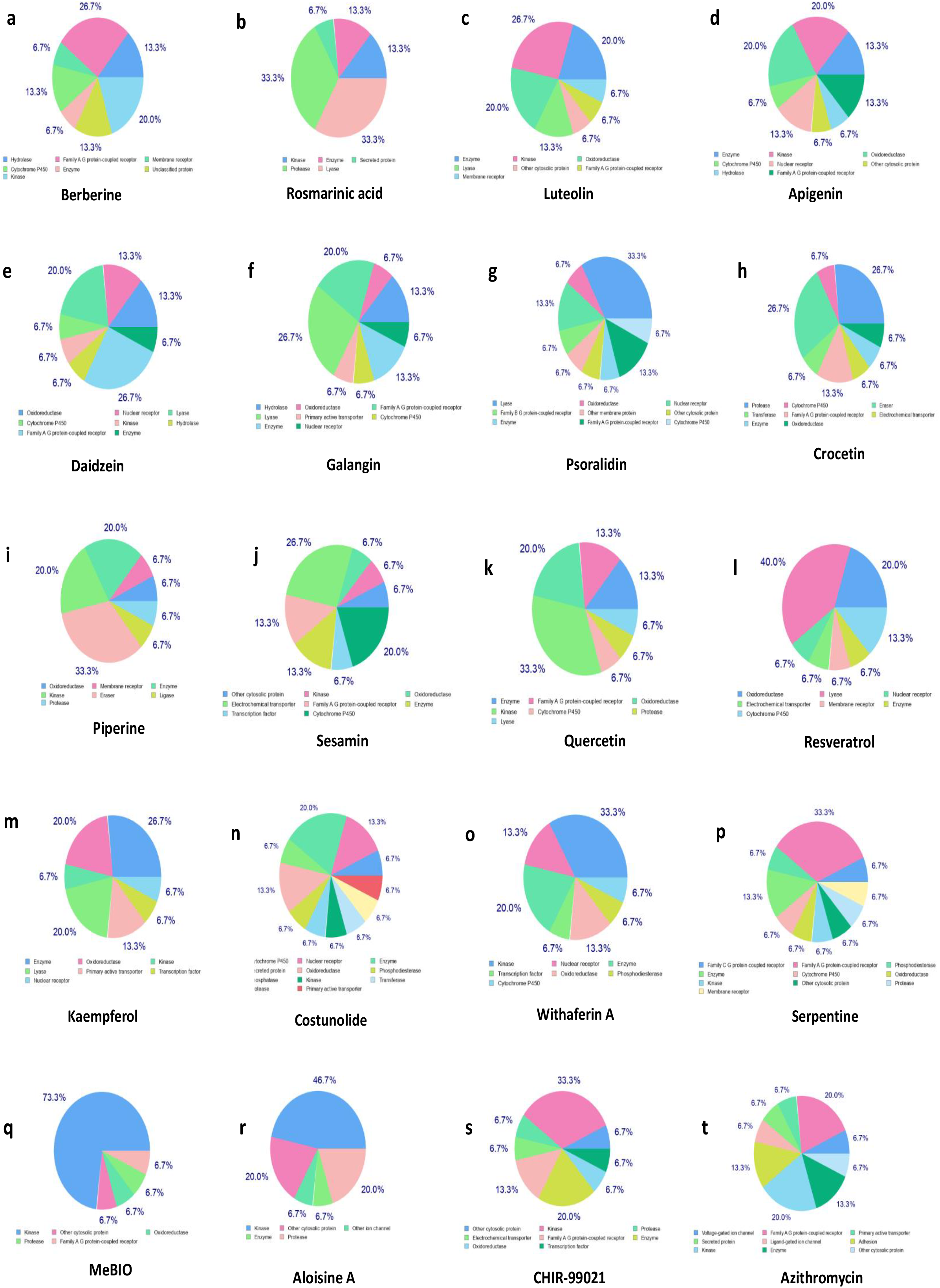
Protein target prediction. **(a-p)** Swiss target prediction of selected phytoconstituents. **(q-u)** Swiss target prediction of included reference drugs.

Graphical representation or bioavailability radar plot or ‘spider’ or ‘cobweb’ plots provide a visual assessment of the drug-likeness parameter of an orally active biomolecule. Bioavailability radar plots provide a preliminary assessment of a potential drug candidate’s medicinal properties. The pink zone (see **Fig 4a-u)** represents physiochemical properties such as lipophilicity (XLOGP3 range between 0.7 to 5.0), size (MW range between 150 to 500), polarity (TPSA range between 20 to 130⍰Å^2^), solubility range (log S⍰≤ ⍰6), saturation range (fraction of carbons in sp^3^ hybridization ≤0.25), and flexibility range (≤9) **(Suppl. Table 6)**. Our analysis showed that all of the selected phytoconstituents exhibited a significant bioavailability radar and score as comparable to standard reference drugs. Berberine, Piperine, Sesamin, Costunolide, Withaferin A, Serpentine and ALOISINE fall within the optimal range of all the above-mentioned six properties, consistent with chemotherapeutic potential.

**Figure 4:**
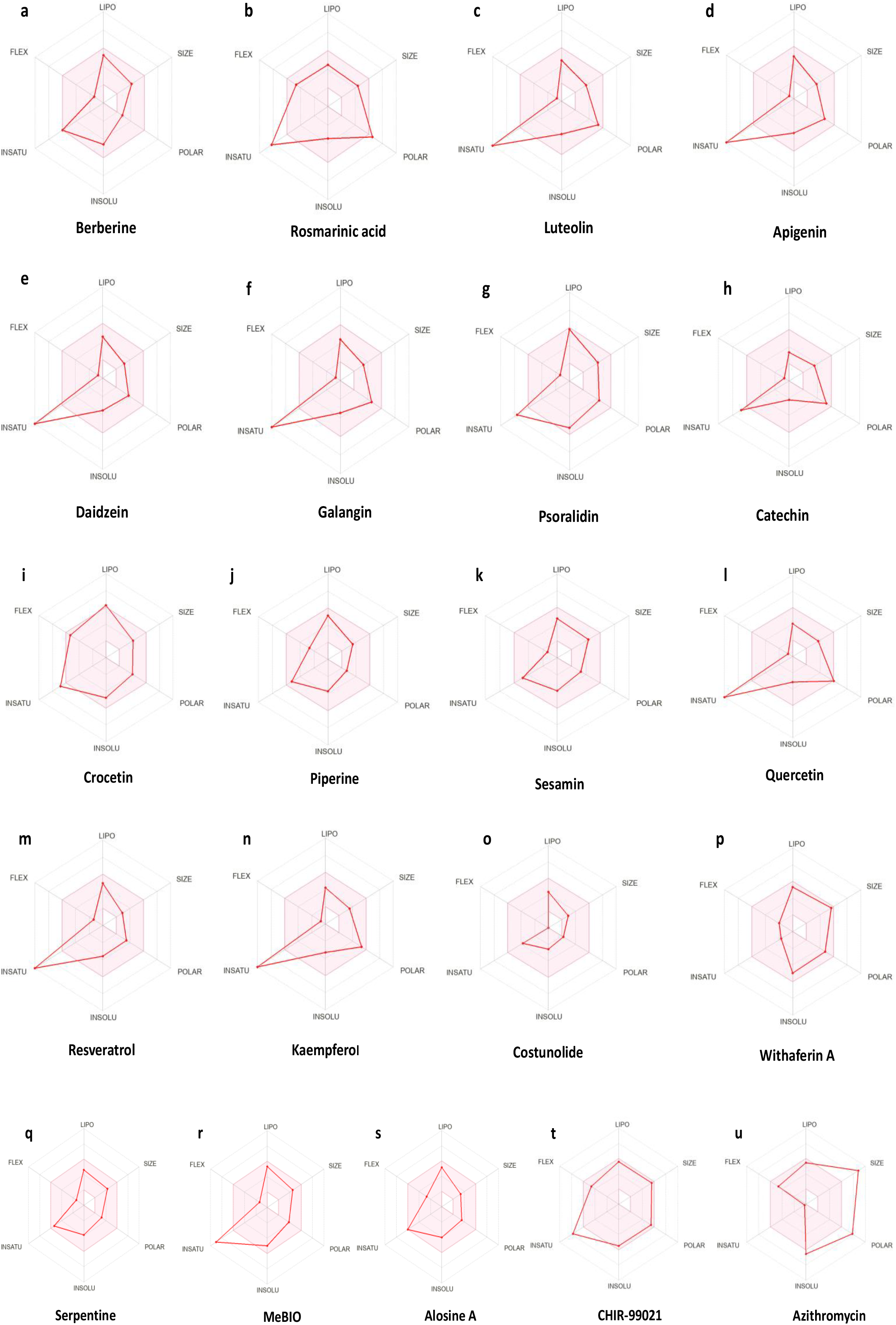
Bioavailability radar plots of natural compounds. **(a-q**) Bioavailability radar plots of selected phytoconstituents and **(r-u)** selected reference control drugs.

### 3.8 Characterization of Psoralidin and Rosmarinic acid for inhibitory potency

Based on the highest predicted binding energy against GSK-3α and GSK-3β, Psoralidin and Rosmarinic acid were selected for further characterization through *in vitro* kinase assays. We used purified enzymes and substrates for the respective GSK-3α and GSK-3β. First, the capacity of Psoralidin to inhibit GSK-3α and GSK-3β was evaluated. A dose-response curve was established for both the kinases post-normalizing the maximum ATP-ADP conversion with DMSO controls **(Suppl. Fig. 5A-D)**. The kinase activity of GSK-3 isoforms decreased upon increasing concentration of Psoralidin **(Fig. 5A-B)**. Similarly, the kinase assay showed Rosmarinic acid to inhibit both GSK3 isoforms **(Fig. 5C-D)**.

**Figure 5:**
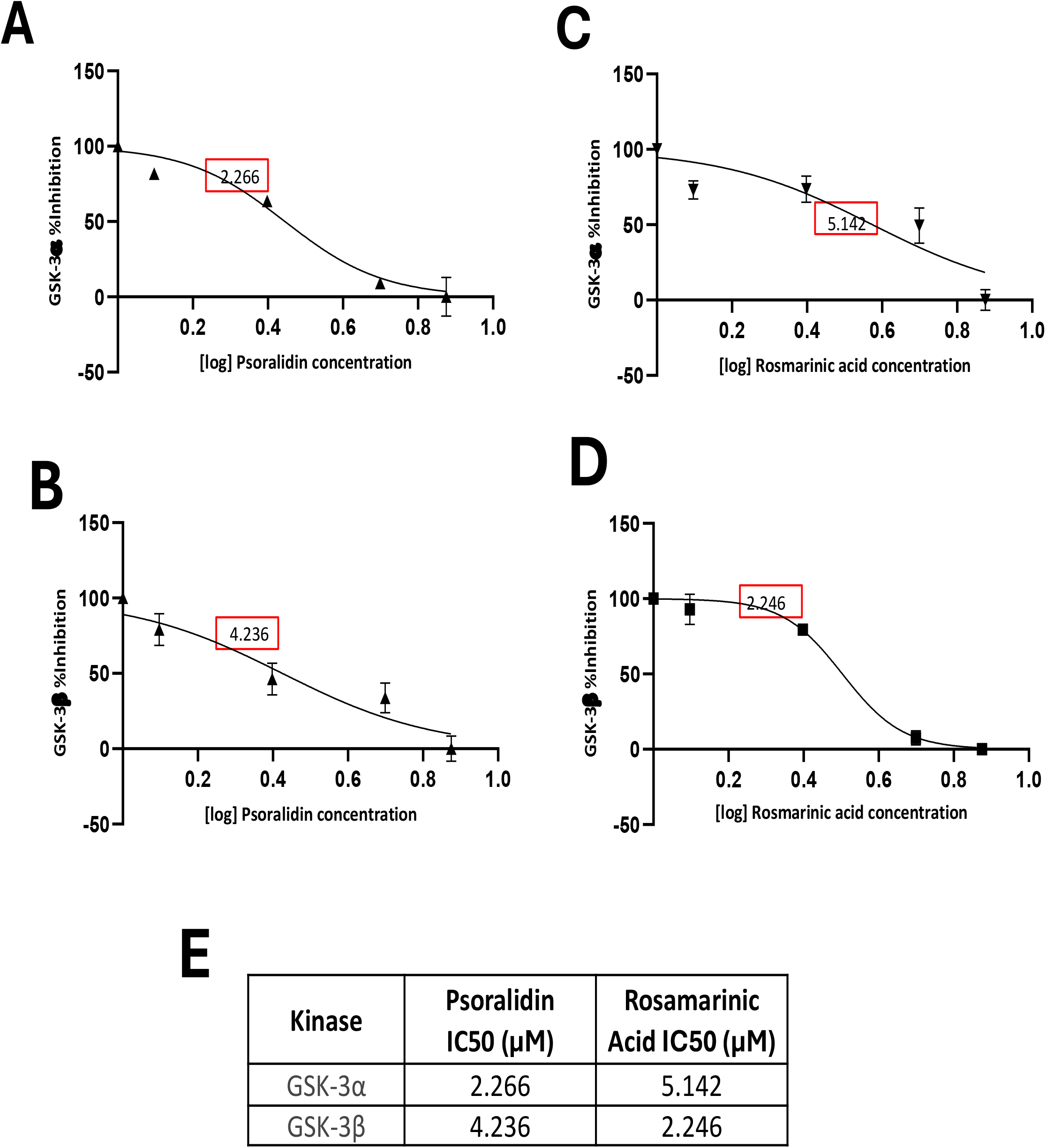
Kinetics and effects of the selected compounds on GSK-3 isoforms. **(A-D)** Kinase assay shows the percent inhibition profile of Rosmarinic acid and Psoralidin IC50 (µM) plotted against 8nM concentration of GSK-3α and GSK-3β. **(E)** IC50 values for respective inhibitors against GSK-3α and GSK-3β.

To further determine the inhibitory characteristics of the compounds, IC50 values were determined **(Fig. 5E)** from the kinase assay as this better indicates inhibitory potency (lower IC50 reflects better inhibitory potency). From the range of selected concentrations tested *in vitro* for both GSK-3α and GSK-3β kinases, we determined 8 nM as an ideal concentration of these kinases to explain the effect of the compounds. At this concentration, Psoralidin showed a better inhibitory effect against GSK-3α with an IC50 of 2.27 µM compared to GSK-3β with an IC50 of 4.24 µM. In contrast, Rosmarinic acid displayed better inhibitory effect against GSK-3β with an IC50 of 2.25 µM compared to GSK-3α with an IC50 of 5.14 µM. These results show that these compounds may serve as GSK-3 inhibitors with ∼2-fold isoform selectivity.

## 4. Discussion

Though it is established that inhibiting GSK-3 isoforms affects the prognosis of various disease conditions, natural inhibitors with minimal toxicity are largely undefined. Here, we screened the natural compounds employing *in silico* and biochemical approaches and identified several potentially novel GSK-3 inhibitors. Rosmarinic acid, Psoralidin, Apigenin, Sesamin, Luteolin, Berberine, Resveratrol, and Piperine were among several other identified potentially novel inhibitors **(Table 1A)**. Of these, Psoralidin and Rosmarinic acid were identified to show better physicochemical and drug-like properties and the highest binding affinities for GSK-3α and GSK-3β respectively **(Fig. 6)**. These observations were further attested by in vitro kinase assay where the kinase assay-driven IC50 for Rosmarinic acid was 2-fold lower for GSK-3β compared to GSK-3α while the IC50 for Psoralidin was 2-fold lower for GSK-3α vs. GSK-3β.

**Figure 6:**
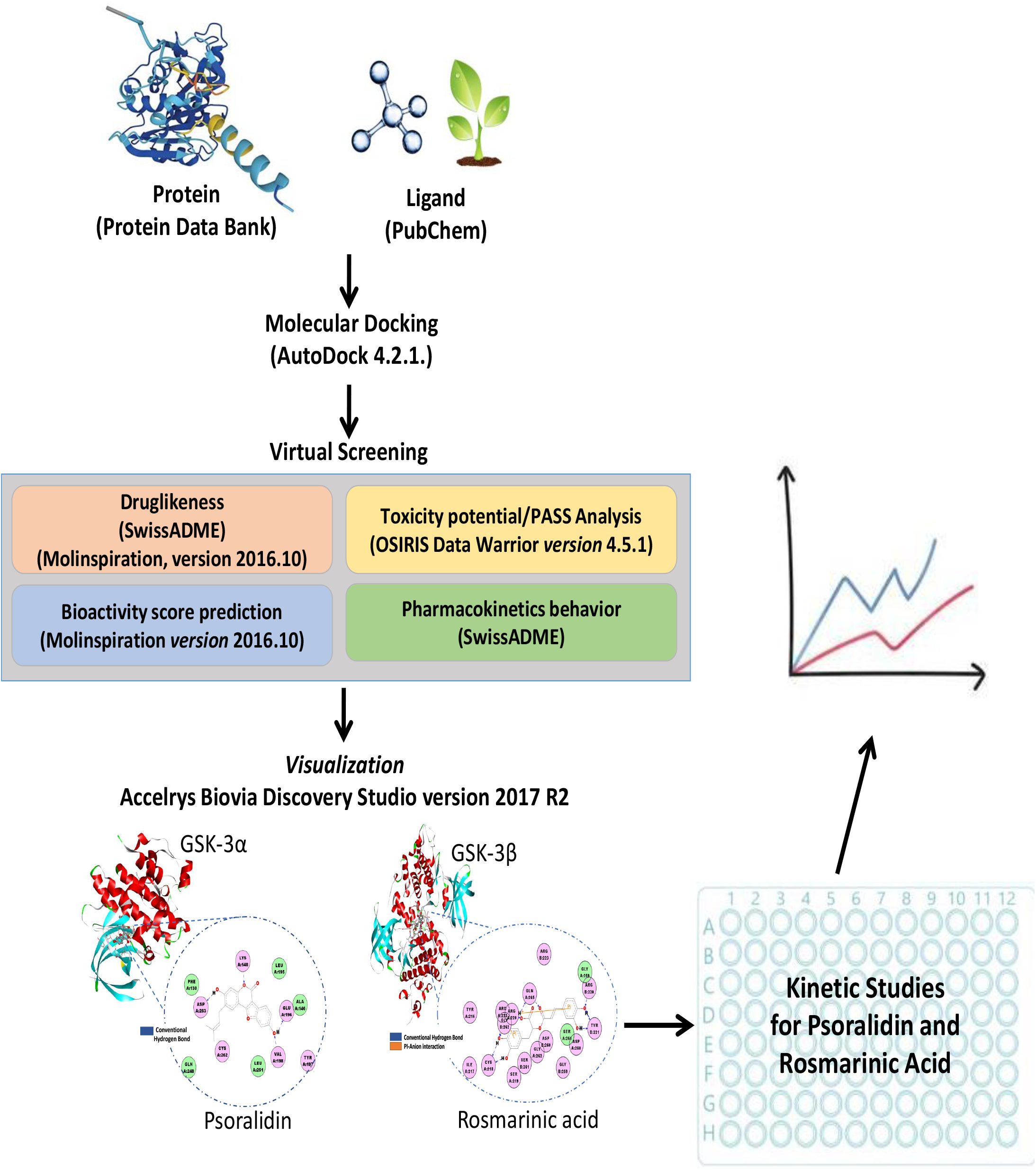
Schematic diagram shows the molecular docking to simulate how natural compounds might bind to the crystal structure of GSK-3 enzymes. This was followed by additional in silico analyses, including assessments of druglikeness, potential toxicity, bioactivity scores, and pharmacokinetic properties. Through this multi-step screening process, Psoralidin and Rosmarinic acid emerged as potential inhibitors of GSK-3α and GSK-3β isoforms, respectively. Finally, based on their predicted high binding energy with both GSK-3α and GSK-3β, Psoralidin and Rosmarinic acid were selected for further testing using a kinase assay. This assay confirmed their inhibitory potential, demonstrating higher potency against GSK-3α and GSK-3β, respectively.

Most of the available synthetic GSK-3 small molecule inhibitors target both isoforms non-specifically. The GSK-3 inhibitor LiCl has been tested in Phase-II and Phase-III clinical trials (ClinicalTrials.gov; NCT00870311, NCT01096082) in patients with neurological disorders. Other phase-II clinical trials (ClinicalTrials.gov; NCT00948259, NCT01049399) have tested the efficacy of GSK-3 inhibitor Tideglusib (NP031112) in patients with Alzheimer’s disease. However, none of them has yet been successfully translated into clinical application and, importantly, systemic prolonged treatment using available synthetic GSK-3 inhibitors might further risk the patients for developing additional clinical conditions. Our *in silico* and *in vitro* studies reveal an interesting interplay between GSK-3 isoforms and Psoralidin. With an IC50 of 2.27 µM against GSK-3α compared to 4.23 µM against GSK-3β, Psoralidin exhibits a marginal yet statistically significant preference for binding with GSK-3α isoform. This higher selectivity of Psoralidin against GSK-3α holds therapeutic promise, as GSK-3α is implicated in a range of human diseases including diabetes [48], ischemic and hypertensive cardiac diseases [7, 11, 19], fatty acid metabolism [49], fibrosis [50], neurological disorders [51], atherosclerosis [52], malignancies [53] and aging [15]. The precise mechanism of higher selectivity of Psoralidin against GSK-3α remains unknown. However, there is a possibility that Psoralidin interacts with the ATP binding pocket of two isoforms differently due to slight variations in amino acid residues lining the pocket which favors GSK-3α isoform. Further investigations employing X-ray crystallography and molecular modeling could enhance our understanding of these complex molecular interactions.

The higher selectivity of Psoralidin for GSK-3α has larger implications. In ischemic cardiac diseases, for instance, where GSK-3α promotes cardiac injury in acute as well as chronic myocardial infarction through mitochondrial dysfunction and activating the pro-apoptotic pathways [8, 11], Psoralidin could offer a more targeted therapeutic approach with potentially fewer side effects associated with pan GSK-3 small molecule inhibitors. While Psoralidin shows a subtle preferential effect for GSK-3α, Rosmarinic acid paints a converse picture by showing higher selectivity for GSK-3β. This opens exciting avenues and makes Rosmarinic acid an attractive natural compound for therapeutic exploration.

While our study identifies Psoralidin and Rosmarinic acid as potential GSK-3 isoform inhibitors, further investigation is required to better understand their broader impact on cellular signaling pathways. While not extensively explored in this context, Psoralidin has been shown to regulate NR1H3-AMPK signaling to limit septic myocardial injury [54]. Additionally, emerging evidence suggests its ability to regulate focal adhesion kinase (FAK), PI3K/Akt, and NF-κB pathways, potentially contributing to its anti-cancer effects [54, 55]. Similarly, Rosmarinic acid influences cell survival through the PI3K/AKT/mTOR pathway and regulates cell proliferation and apoptosis via p53, p21, and cell cyclins in cancer cells [56, 57]. Beyond its potential in cancer, Rosmarinic acid exhibits cardioprotective effects. Studies report its ability to limit ischemia/reperfusion (I/R)-induced inflammation and cardiomyocyte apoptosis through activation of PPARγ and overexpression of SERCA2 and RyR2 genes, both crucial for Ca2+ homeostasis [58, 59]. Additionally, *in vivo* studies demonstrate its role in inhibiting NF-κB inflammatory signaling, reducing ROS production, and limiting mitochondrial impairment *via* the Sirt1/PGC-1α pathway, ultimately protecting against cardiac I/R injury [60]. Given the established role of GSK-3 isoforms in most of the discussed signaling pathways, the observed protective effects of Psoralidin and Rosmarinic acid may involve modulation of these isoforms.

Emerging evidence suggests that GSK-3β plays critical roles in the pathophysiology of numerous cancers, where modified GSK-3β function promotes tumor growth by inducing cell proliferation and survival [61]. Moreover, GSK-3β regulates the pathogenesis of several neurological disorders including the synaptic deficit, neuro-inflammation and Parkinson’s disease [62-64]. Preferential inhibition of GSK-3β by Rosmarinic acid suggests potential novel anti-cancer agent and, as an agent to treat specific neurological disorders. Furthermore, its selectivity for GSK-3β might show beneficial effects in inflammatory diseases like rheumatoid arthritis and psoriasis, where GSK-3β plays a key role in inflammatory pathways and immune cell activation [65].

It is important to mention that while IC50 value provides valuable insights into the relative potencies of inhibitors, it represents only one aspect of a compound’s inhibitory potential. Other properties such as cellular uptake, tissue distribution and metabolic stability also play a crucial role in determining their *in vivo* efficacy. We identified favorable bioactivity and drug-likeness scores for Psoralidin and Rosmarinic acid against GSK-3α and GSK-3β that further suggest their therapeutic potential. Moreover, Rosmarinic acid was predicted to have no mutagenic, tumorigenic and irritant effects and no adverse effects on the reproductive system while Psoralidin was predicted to have no mutagenic, tumorigenic or irritant effects however, predicted to show an adverse effect on the reproductive system. Our finding on Psoralidin is consistent with previous reports where germline global genetic targeting of GSK-3α causes male infertility due to altered ATP levels and impaired sperm motility [66]. Further analysis predicted Rosmarinic acid to have low GI absorption while Psoralidin to have high absorption and both were predicted to be incapable of penetrating the BBB. Therefore, future studies to investigate these pharmacokinetic parameters for these compounds are important to better assess their therapeutic potential.

In summary, here we have computationally screened a candidate series of natural compounds for GSK-3 isoform-specific natural inhibition and identified Psoralidin and Rosmarinic acid. The natural origins of these compounds may offer a better safety profile with minimal side effects compared to the available synthetic small molecule inhibitors. Further detailed studies related to *in vivo* efficacy and safety assessments, structure-activity relationship and mechanistic insight are required to translate these promising findings for patients suffering from GSK-3-driven pathological conditions. Moreover, investigating the cellular and molecular regulatory mechanisms of Psoralidin and Rosmarinic acid will help better understand GSK-3-driven actions.

## 5. Declaration

### 5.2 Authors’ contributions

FA designed the study, acquired funding, supervised the project, data interpretation, and wrote the manuscript, AG and HM performed experiments, collected data and helped with manuscript writing, MAS and JRW helped with data interpretation and manuscript editing, RQ helped with data analysis, interpretation and manuscript writing.

### 5.4 Availability of Data

Data is included in the manuscript and supplementary files. Additional information is available from the corresponding author upon a reasonable request.

## References

[1] Lal H, Ahmad F, Woodgett J, Force T. The GSK-3 family as therapeutic target for myocardial diseases. Circ Res. 2015;116:138–49.

[2] Gupte M, Lal H, Ahmad F, Sawyer DB, Hill MF. Chronic Neuregulin-1beta Treatment Mitigates the Progression of Postmyocardial Infarction Heart Failure in the Setting of Type 1 Diabetes Mellitus by Suppressing Myocardial Apoptosis, Fibrosis, and Key Oxidant-Producing Enzymes. J Card Fail. 2017;23:887–99.

[3] Yusuf AM, Qaisar R, Al-Tamimi AO, Jayakumar MN, Woodgett JR, Koch WJ, et al. Cardiomyocyte-GSK-3beta deficiency induces cardiac progenitor cell proliferation in the ischemic heart through paracrine mechanisms. J Cell Physiol. 2022;237:1804–17.

[4] Force T, Woodgett JR. Unique and overlapping functions of GSK-3 isoforms in cell differentiation and proliferation and cardiovascular development. J Biol Chem. 2009;284:9643–7.

[5] Umbarkar P, Ruiz Ramirez SY, Toro Cora A, Tousif S, Lal H. GSK-3 at the heart of cardiometabolic diseases: Isoform-specific targeting is critical to therapeutic benefit. Biochim Biophys Acta Mol Basis Dis. 2023;1869:166724.

[6] Woodgett JR. Regulation and functions of the glycogen synthase kinase-3 subfamily. Sem Cancer Biol. 1994;5:269–75.

[7] Ahmad F, Singh AP, Tomar D, Rahmani M, Zhang Q, Woodgett JR, et al. Cardiomyocyte-GSK-3α promotes mPTP opening and heart failure in mice with chronic pressure overload. J Mol Cell Cardiol. 2019;130:65–75.

[8] Ahmad F, Lal H, Zhou J, Vagnozzi RJ, Yu JE, Shang X, et al. Cardiomyocyte-specific deletion of gsk3alpha mitigates post-myocardial infarction remodeling, contractile dysfunction, and heart failure. J Am Coll Cardiol. 2014;64:696–706.

[9] Matsuda T, Zhai P, Maejima Y, Hong C, Gao S, Tian B, et al. Distinct roles of GSK-3alpha and GSK-3beta phosphorylation in the heart under pressure overload. Proc Natl Acad Sci U S A. 2008;105:20900–5.

[10] Lal H, Ahmad F, Zhou J, Yu JE, Vagnozzi RJ, Guo Y, et al. Cardiac fibroblast glycogen synthase kinase-3beta regulates ventricular remodeling and dysfunction in ischemic heart. Circulation. 2014;130:419–30.

[11] Ahmad F, Marzook H, Gupta A, Aref A, Patil K, Khan AA, et al. GSK-3alpha aggravates inflammation, metabolic derangement, and cardiac injury post-ischemia/reperfusion. J Mol Med (Berl). 2023;101:1379–96.

[12] Woodgett JR, Cohen P. Multisite phosphorylation of glycogen synthase. Molecular basis for the substrate specificity of glycogen synthase kinase-3 and casein kinase-II (glycogen synthase kinase-5). Biodhim Biophys Acta. 1984;788:339–47.

[13] Kerkela R, Kockeritz L, Macaulay K, Zhou J, Doble BW, Beahm C, et al. Deletion of GSK-3beta in mice leads to hypertrophic cardiomyopathy secondary to cardiomyoblast hyperproliferation. J Clin Invest. 2008;118:3609–18.

[14] Lal H, Zhou J, Ahmad F, Zaka R, Vagnozzi RJ, Decaul M, et al. Glycogen synthase kinase-3alpha limits ischemic injury, cardiac rupture, post-myocardial infarction remodeling and death. Circulation. 2012;125:65–75.

[15] Zhou J, Freeman TA, Ahmad F, Shang X, Mangano E, Gao E, et al. GSK-3alpha is a central regulator of age-related pathologies in mice. J Clin Invest. 2013;123:1821–32.

[16] Zhou J, Ahmad F, Lal H, Force T. Response by Zhou et al to Letter Regarding Article, “Loss of Adult Cardiac Myocyte GSK-3 Leads to Mitotic Catastrophe Resulting in Fatal Dilated Cardiomyopathy”. Circ Res. 2016;119:e29–e30.

[17] Zhou J, Ahmad F, Parikh S, Hoffman NE, Rajan S, Verma VK, et al. Loss of Adult Cardiac Myocyte GSK-3 Leads to Mitotic Catastrophe Resulting in Fatal Dilated Cardiomyopathy. Circ Res. 2016;118:1208–22.

[18] Papadopoli D, Pollak M, Topisirovic I. The role of GSK3 in metabolic pathway perturbations in cancer. Biochim Biophys Acta Mol Cell Res. 2021;1868:119059.

[19] Ahmad F, Woodgett JR. Emerging roles of GSK-3α in pathophysiology: Emphasis on cardio-metabolic disorders. Biochim Biophys Acta Mol Cell Res. 2020;1867:118616.

[20] Kramer T, Schmidt B, Lo Monte F. Small-Molecule Inhibitors of GSK-3: Structural Insights and Their Application to Alzheimer’s Disease Models. Int J Alzheimers Dis. 2012;2012:381029.

[21] Dajani R, Fraser E, Roe SM, Young N, Good V, Dale TC, et al. Crystal structure of glycogen synthase kinase 3ß: a structural basis for phosphate-primed substrate specificity and autoinhibition. Cell. 2001;105:721–32.

[22] Beurel E, Grieco SF, Jope RS. Glycogen synthase kinase-3 (GSK3): regulation, actions, and diseases. Pharmacol Ther. 2015;148:114–31.

[23] Duda P, Akula SM, Abrams SL, Steelman LS, Martelli AM, Cocco L, et al. Targeting GSK3 and Associated Signaling Pathways Involved in Cancer. Cells. 2020;9:1110.

[24] Xin Z, Wu X, Yu Z, Shang J, Xu B, Jiang S, et al. Mechanisms explaining the efficacy of psoralidin in cancer and osteoporosis, a review. Pharmacol Res. 2019;147:104334.

[25] Sharifi-Rad J, Kamiloglu S, Yeskaliyeva B, Beyatli A, Alfred MA, Salehi B, et al. Pharmacological Activities of Psoralidin: A Comprehensive Review of the Molecular Mechanisms of Action. Front Pharmacol. 2020;11:571459.

[26] Liang Z, Chen Y, Wang Z, Wu X, Deng C, Wang C, et al. Protective effects and mechanisms of psoralidin against adriamycin-induced cardiotoxicity. J Adv Res. 2022;40:249–61.

[27] Jin Z, Yan W, Jin H, Ge C, Xu Y. Differential effect of psoralidin in enhancing apoptosis of colon cancer cells via nuclear factor-kappaB and B-cell lymphoma-2/B-cell lymphoma-2-associated X protein signaling pathways. Oncol Lett. 2016;11:267–72.

[28] Luo C, Zou L, Sun H, Peng J, Gao C, Bao L, et al. A Review of the Anti-Inflammatory Effects of Rosmarinic Acid on Inflammatory Diseases. Front Pharmacol. 2020;11:153.

[29] Dotolo S, Cervellera C, Russo M, Russo GL, Facchiano A. Virtual Screening of Natural Compounds as Potential PI(3)K-AKT1 Signaling Pathway Inhibitors and Experimental Validation. Molecules. 2021;26.

[30] Gupta A, Ahmad R, Siddiqui S, Yadav K, Srivastava A, Trivedi A, et al. Flavonol morin targets host ACE2, IMP-α, PARP-1 and viral proteins of SARS-CoV-2, SARS-CoV and MERS-CoV critical for infection and survival: a computational analysis. J Biomol Struct Dyn. 2022;40:5515–46.

[31] Ahmad R. Steroidal glycoalkaloids from Solanum nigrum target cytoskeletal proteins: an in silico analysis. PeerJ. 2019;7:e6012.

[32] Schwede T, Kopp J, Guex N, Peitsch MC. SWISS-MODEL: An automated protein homology-modeling server. Nucleic Acids Res. 2003;31:3381–5.

[33] Davis IW, Leaver-Fay A, Chen VB, Block JN, Kapral GJ, Wang X, et al. MolProbity: all-atom contacts and structure validation for proteins and nucleic acids. Nucleic Acids Res. 2007;35:W375–83.

[34] Lipinski CA, Lombardo F, Dominy BW, Feeney PJ. Experimental and computational approaches to estimate solubility and permeability in drug discovery and development settings. Adv Drug Deliv Rev. 2001;46:3–26.

[35] Khan T, Dixit S, Ahmad R, Raza S, Azad I, Joshi S, et al. Molecular docking, PASS analysis, bioactivity score prediction, synthesis, characterization and biological activity evaluation of a functionalized 2-butanone thiosemicarbazone ligand and its complexes. J Chem Biol. 2017;10:91–104.

[36] Veber DF, Johnson SR, Cheng HY, Smith BR, Ward KW, Kopple KD. Molecular properties that influence the oral bioavailability of drug candidates. J Med Chem. 2002;45:2615–23.

[37] Ghose AK, Viswanadhan VN, Wendoloski JJ. A knowledge-based approach in designing combinatorial or medicinal chemistry libraries for drug discovery. 1. A qualitative and quantitative characterization of known drug databases. J Comb Chem. 1999;1:55–68.

[38] Egan WJ, Merz KM, Jr., Baldwin JJ. Prediction of drug absorption using multivariate statistics. J Med Chem. 2000;43:3867–77.

[39] Muegge I, Heald SL, Brittelli D. Simple selection criteria for drug-like chemical matter. J Med Chem. 2001;44:1841–6.

[40] Daina A, Michielin O, Zoete V. SwissADME: a free web tool to evaluate pharmacokinetics, drug-likeness and medicinal chemistry friendliness of small molecules. Sci Rep. 2017;7:42717.

[41] Ritchie TJ, Ertl P, Lewis R. The graphical representation of ADME-related molecule properties for medicinal chemists. Drug Discov Today. 2011;16:65–72.

[42] Ansari JA, Ahmad MK, Fatima N, Azad I, Mahdi AA, Satyanarayan GNV, et al. Chemical Characterization, In-silico Evaluation, and Molecular Docking Analysis of Antiproliferative Compounds Isolated from the Bark of Anthocephalus cadamba Miq. Anticancer Agents Med Chem. 2022;22:3416–37.

[43] Geldenhuys WJ, Mohammad AS, Adkins CE, Lockman PR. Molecular determinants of blood-brain barrier permeation. Ther Deliv. 2015;6:961–71.

[44] Lin JH, Yamazaki M. Role of P-glycoprotein in pharmacokinetics: clinical implications. Clin Pharmacokinet. 2003;42:59–98.

[45] Dai Z, Wu Y, Xiong Y, Wu J, Wang M, Sun X, et al. CYP1A inhibitors: Recent progress, current challenges, and future perspectives. Med Res Rev. 2024;44:169–234.

[46] Wang JS, DeVane CL. Involvement of CYP3A4, CYP2C8, and CYP2D6 in the metabolism of (R)- and (S)-methadone in vitro. Drug Metab Dispos. 2003;31:742–7.

[47] Gfeller D, Grosdidier A, Wirth M, Daina A, Michielin O, Zoete V. SwissTargetPrediction: a web server for target prediction of bioactive small molecules. Nucleic Acids Res. 2014;42:W32–8.

[48] MacAulay K, Doble BW, Patel S, Hansotia T, Sinclair EM, Drucker DJ, et al. Glycogen synthase kinase 3alpha-specific regulation of murine hepatic glycogen metabolism. Cell Metab. 2007;6:329–37.

[49] Nakamura M, Liu T, Husain S, Zhai P, Warren JS, Hsu CP, et al. Glycogen Synthase Kinase-3alpha Promotes Fatty Acid Uptake and Lipotoxic Cardiomyopathy. Cell Metab. 2019.

[50] Umbarkar P, Ejantkar S, Ruiz Ramirez SY, Toro Cora A, Zhang Q, Tousif S, et al. Cardiac fibroblast GSK-3alpha aggravates ischemic cardiac injury by promoting fibrosis, inflammation, and impairing angiogenesis. Basic Res Cardiol. 2023;118:35.

[51] Ma T. GSK3 in Alzheimer’s disease: mind the isoforms. J Alzheimers Dis. 2014;39:707–10.

[52] McAlpine CS, Huang A, Emdin A, Banko NS, Beriault DR, Shi Y, et al. Deletion of Myeloid GSK3alpha Attenuates Atherosclerosis and Promotes an M2 Macrophage Phenotype. Arterioscler Thromb Vasc Biol. 2015;35:1113–22.

[53] Banerji V, Frumm SM, Ross KN, Li LS, Schinzel AC, Hahn CK, et al. The intersection of genetic and chemical genomic screens identifies GSK-3alpha as a target in human acute myeloid leukemia. J Clin Invest. 2012;122:935–47.

[54] Yang Y, Lei W, Qian L, Zhang S, Yang W, Lu C, et al. Activation of NR1H3 signaling pathways by psoralidin attenuates septic myocardial injury. Free Radic Biol Med. 2023;204:8–19.

[55] Jin Z, Yan W, Jin H, Ge C, Xu Y. Psoralidin inhibits proliferation and enhances apoptosis of human esophageal carcinoma cells via NF-kappaB and PI3K/Akt signaling pathways. Oncol Lett. 2016;12:971–6.

[56] Wang L, Yang H, Wang C, Shi X, Li K. Rosmarinic acid inhibits proliferation and invasion of hepatocellular carcinoma cells SMMC 7721 via PI3K/AKT/mTOR signal pathway. Biomed Pharmacother. 2019;120:109443.

[57] Sirajudeen F, Bou Malhab LJ, Bustanji Y, Shahwan M, Alzoubi KH, Semreen MH, et al. Exploring the Potential of Rosemary Derived Compounds (Rosmarinic and Carnosic Acids) as Cancer Therapeutics: Current Knowledge and Future Perspectives. Biomol Ther (Seoul). 2024;32:38–55.

[58] Han J, Wang D, Ye L, Li P, Hao W, Chen X, et al. Rosmarinic Acid Protects against Inflammation and Cardiomyocyte Apoptosis during Myocardial Ischemia/Reperfusion Injury by Activating Peroxisome Proliferator-Activated Receptor Gamma. Front Pharmacol. 2017;8:456.

[59] Javidanpour S, Dianat M, Badavi M, Mard SA. The cardioprotective effect of rosmarinic acid on acute myocardial infarction and genes involved in Ca(2+) homeostasis. Free Radic Res. 2017;51:911–23.

[60] Quan W, Liu HX, Zhang W, Lou WJ, Gong YZ, Yuan C, et al. Cardioprotective effect of rosmarinic acid against myocardial ischaemia/reperfusion injury via suppression of the NF-kappaB inflammatory signalling pathway and ROS production in mice. Pharm Biol. 2021;59:222–31.

[61] Domoto T, Uehara M, Bolidong D, Minamoto T. Glycogen Synthase Kinase 3beta in Cancer Biology and Treatment. Cells. 2020;9.

[62] Li J, Ma S, Chen J, Hu K, Li Y, Zhang Z, et al. GSK-3beta Contributes to Parkinsonian Dopaminergic Neuron Death: Evidence From Conditional Knockout Mice and Tideglusib. Front Mol Neurosci. 2020;13:81.

[63] Banach E, Jaworski T, Urban-Ciecko J. Early synaptic deficits in GSK-3beta overexpressing mice. Neurosci Lett. 2022;784:136744.

[64] Lei P, Ayton S, Bush AI, Adlard PA. GSK-3 in Neurodegenerative Diseases. Int J Alzheimers Dis. 2011;2011:189246.

[65] Arioka M, Takahashi-Yanaga F. Glycogen synthase kinase-3 inhibitor as a multi-targeting anti-rheumatoid drug. Biochem Pharmacol. 2019;165:207–13.

[66] Bhattacharjee R, Goswami S, Dey S, Gangoda M, Brothag C, Eisa A, et al. Isoform-specific requirement for GSK3alpha in sperm for male fertility. Biol Reprod. 2018;99:384–94.

