## Supplementary material for "Natural compound screening predicts novel GSK-3 isoform-specific inhibitors": Suppl. Table 1A-B

**Supplementary table 1A;** List of Natural ligands

| S.No. | Natural Compound Name | Pubchem ID | Molecular Formula | Molecular Weight (g/mol) | Structure |
|---|---|---|---|---|---|
| 1 | Silymarin | 5213 | C25H22O10 | 482.4 | 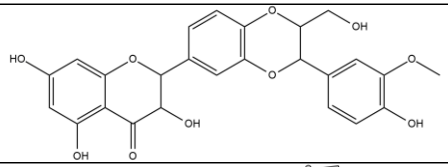 |
| 2 | Berberine | 2353 | C20H18NO4+ | 336.4 | 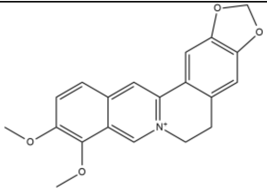 |
| 3 | Rosmarinic acid | 5281792 | C18H16O8 | 360.3 | 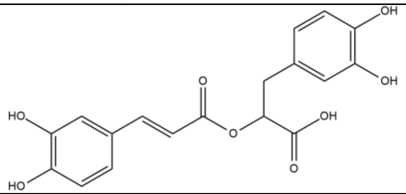 |
| 4 | Rabdosin A | 102034411 | C21H28O6 | 376.4 | 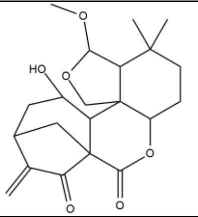 |
| 5 | Arjunolic acid | 73641 | C30H48O5 | 488.7 | 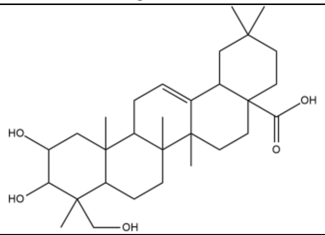 |

| | | | | | |
|---|---|---|---|---|---|
| 6 | Myricetin | 5281672 | C15H10O8 | 318.23 | 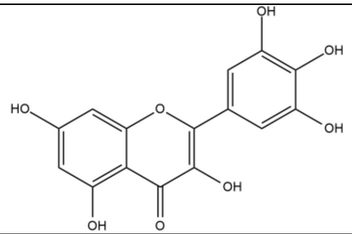 |
| 7 | Honokiol | 72303 | C18H18O2 | 266.3 | 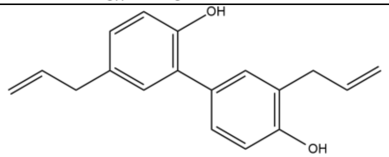 |
| 8 | Lutein | 5281243 | C40H56O2 | 568.9 | 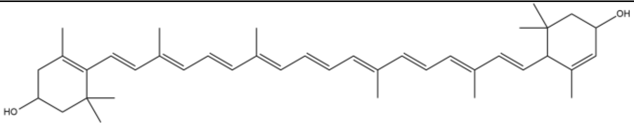 |
| 9 | Luteolin | 5280445 | C15H10O6 | 286.24 | 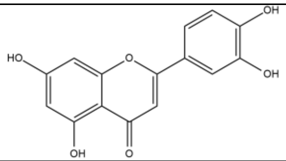 |
| 10 | Anethole | 637563 | C10H12O | 148.2 | 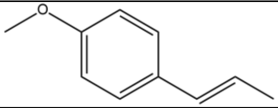 |
| 11 | Apigenin | 5280443 | C15H10O5 | 270.24 | 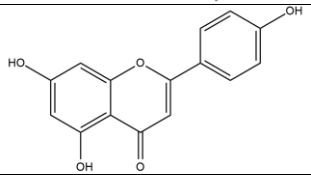 |
| 12 | Betulinic acid | 64971 | C30H48O3 | 456.7 | 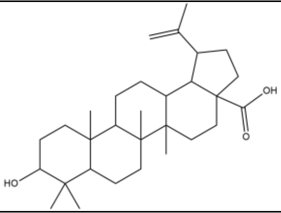 |

| | | | | | |
|---|---|---|---|---|---|
| 13 | Caffeic acid | 689043 | C9H8O4 | 180.16 | 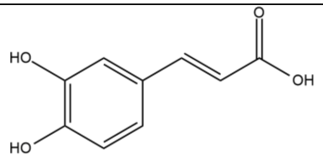 |
| 14 | Cinnamaldehyde | 637511 | C9H8O | 132.16 | 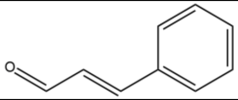 |
| 15 | Cinnamic acid | 444539 | C9H8O2 | 148.16 | 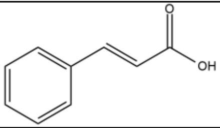 |
| 16 | Coumarin | 323 | C9H6O2 | 146.14 | 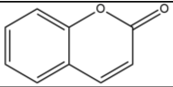 |
| 17 | Daidzein | 5281708 | C15H10O4 | 254.24 | 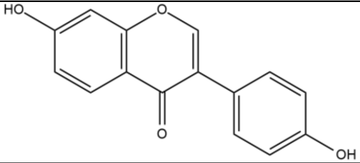 |
| 18 | Emodin | 3220 | C15H10O5 | 270.24 | 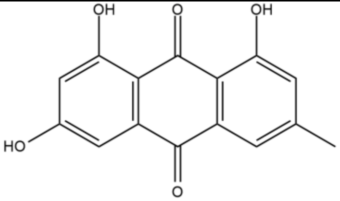 |
| 19 | Eugenol | 3314 | C10H12O2 | 164.2 | 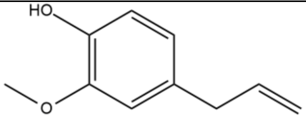 |
| 20 | Galangin | 5281616 | C15H10O5 | 270.24 | 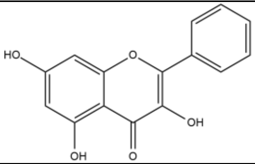 |

| | | | | | |
|---|---|---|---|---|---|
| 21 | Ginsenosides | 3086007 | C30H52O2 | 444.7 | 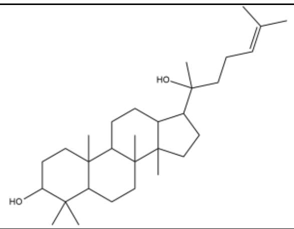 |
| 22 | Morphine | 5288826 | C17H19NO3 | 285.34 | 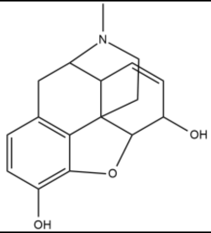 |
| 23 | Oleanolic acid | 10494 | C30H48O3 | 456.7 | 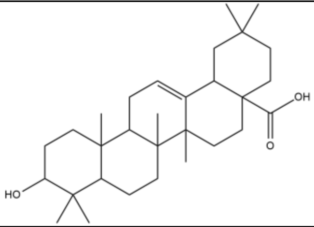 |
| 24 | Psoralidin | 5281806 | C20H16O5 | 336.3 | 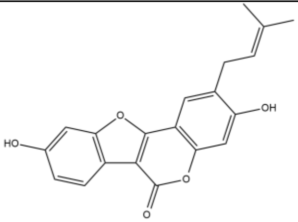 |
| 25 | Thymoquinone | 10281 | C10H12O2 | 164.2 | 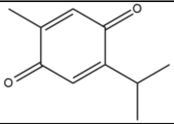 |
| 26 | Tocotrienol | 9929901 | C26H38O2 | 382.6 | 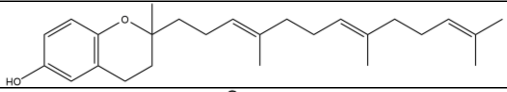 |
| 27 | Allicin | 65036 | C6H10OS2 | 162.3 | 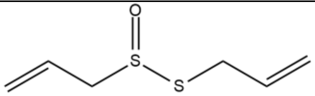 |

| | | | | | |
|---|---|---|---|---|---|
| 28 | alpha-Pinene | 6654 | C10H16 | 136.23 | 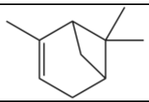 |
| 29 | Beta-caryophyllene | 5281515 | C15H24 | 204.35 | 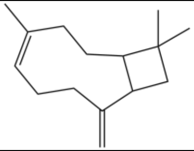 |
| 30 | Carvone | 7439 | C10H14O | 150.22 | 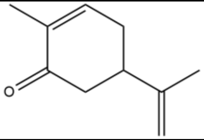  |
| 31 | Catechin | 9064 | C15H14O6 | 290.27 | 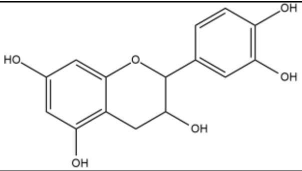  |
| 32 | Crocetin | 5281232 | C20H24O4 | 328.4 | 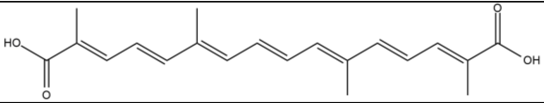  |
| 33 | Cuminaldehyde | 326 | C10H12O | 148.20 | 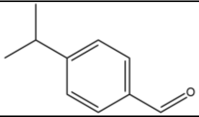  |
| 34 | Curcumin | 969516 | C21H20O6 | 368.4 | 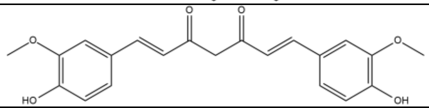  |
| 35 | Gingerol | 442793 | C17H26O4 | 294.4 | 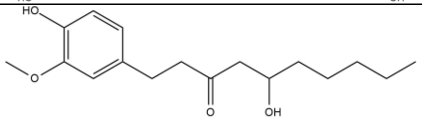  |
| 36 | Limonene | 22311 | C10H16 | 136.23 | 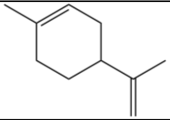 |
| 37 | Myrcene | 31253 | C10H16 | 136.23 | 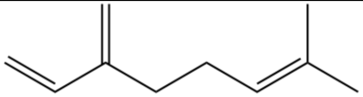 |

| | | | | | |
|---|---|---|---|---|---|
| 38 | Piperine | 638024 | C17H19NO3 | 285.34 | 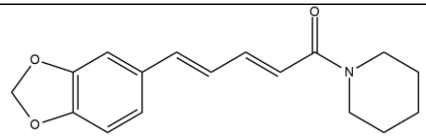 |
| 39 | Sabinene | 18818 | 10H16 | 136.23 | 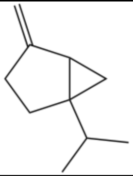 |
| 40 | Shogaol | 5281794 | C17H24O3 | 276.4 | 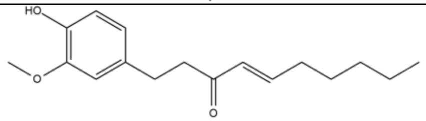 |
| 41 | Thymol | 6989 | C10H14O | 150.22 | 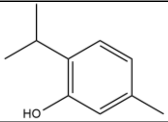 |
| 42 | Trigonelline | 5570 | C7H7NO2 | 137.14 | 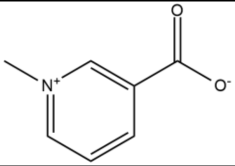 |
| 43 | Zingiberene | 92776 | C15H24 | 204.35 | 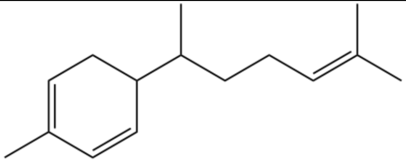 |
| 44 | turmerone | 14367555 | C15H22O | 218.33 | 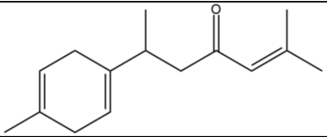 |
| 45 | Carbazole | 6854 | C12H9N | 167.21 | 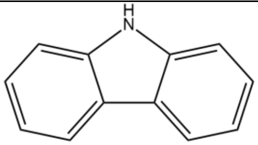 |

| | | | | | |
|---|---|---|---|---|---|
| 46 | Carindone | 101316738 | C31H44O6 | 512.7 | 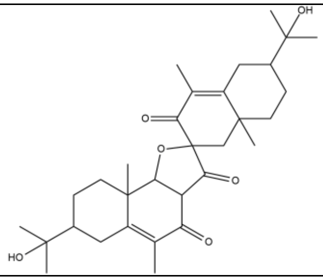 |
| 47 | Sesamin | 72307 | C20H18O6 | 354.4 | 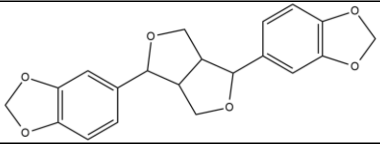 |
| 48 | Linalool | 6549 | C10H18O | 154.25 | 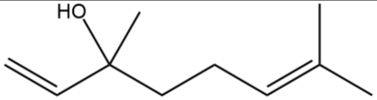 |
| 49 | Moringin | 14865502 | C14H17NO5S | 311.36 | 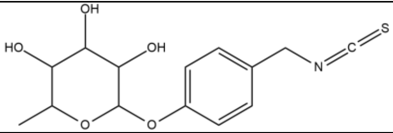 |
| 50 | Isoquercitrin | 5280804 | C21H20O12 | 464.4 | 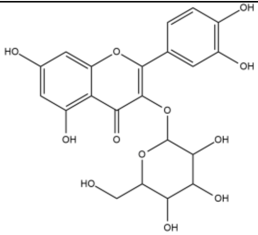 |
| 51 | Quercetin | 5280343 | C15H10O7 | 302.23 | 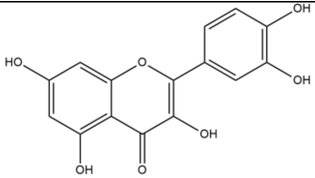 |
| 52 | Resveratrol | 445154 | C14H12O3 | 228.24 | 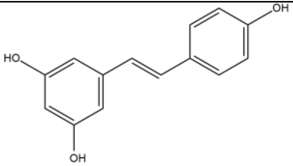 |

| | | | | | |
|---|---|---|---|---|---|
| 53 | Kaempferol | 5280863 | C15H10O6 | 286.24 | 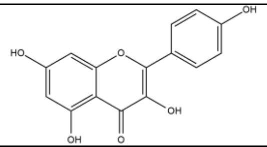 |
| 54 | Costunolide | 5281437 | C15H20O2 | 232.32 | 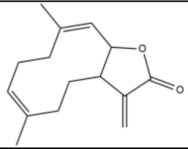 |
| 55 | Kumatakenin | 5318869 | C17H14O6 | 314.29 | 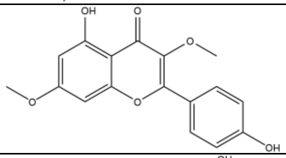 |
| 56 | Mangiferin | 5281647 | C19H18O11 | 422.3 | 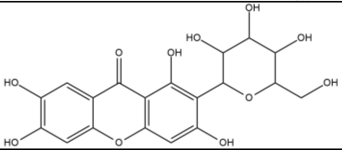 |
| 57 | Tinosporaside | 14194109 | C25H32O10 | 492.5 | 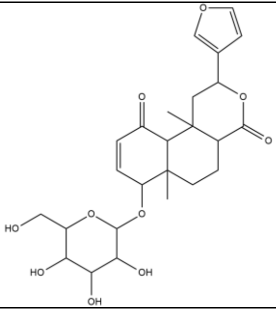 |
| 58 | Epigallocatechin gallate | 65064 | C22H18O11 | 458.4 | 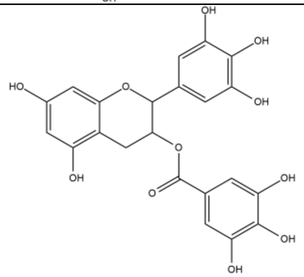 |

| | | | | | |
|---|---|---|---|---|---|
| 59 | Ellagic acid | 5281855 | C14H6O8 | 302.19 | 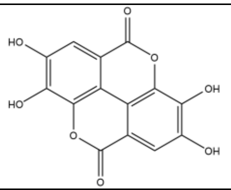 |
| 60 | Withaferin A | 265237 | C28H38O6 | 470.6 | 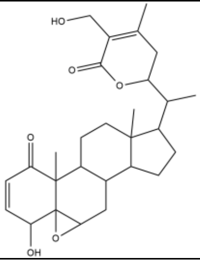 |
| 61 | withanolide A | 11294368 | C28H38O6 | 470.6 | 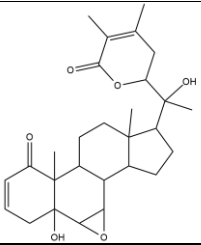 |
| 62 | Serpentine | 73073 | C21H20N2O3 | 348.4 | 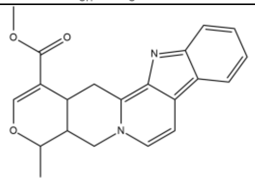 |
| 63 | Naringin | 442428 | C27H32O14 | 580.5 | 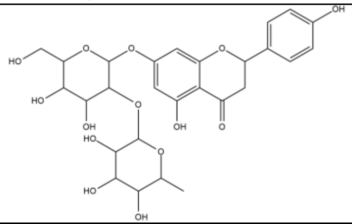 |

| | | | | | |
|---|---|---|---|---|---|
| 64 | Hesperidin | 10621 | C28H34O15 | 610.6 | 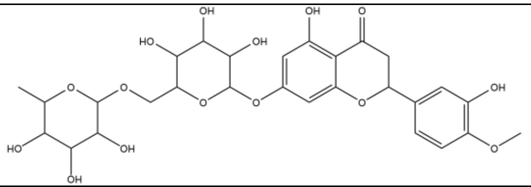 |
| 65 | Phloretin | 4788 | C15H14O5 | 274.27 | 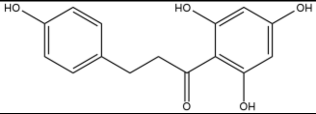 |
| 66 | Neral | 643779 | C10H16O | 152.23 | 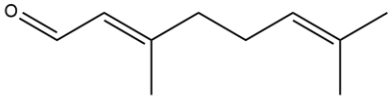 |
| 67 | Oleic acid | 445639 | C18H34O2 | 282.5 | 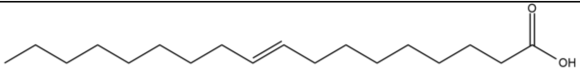 |
| 68 | Citric acid | 311 | C6H8O7 | 192.12 | 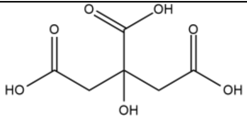 |
| 69 | Rutin | 5280805 | C27H30O16 | 610.5 | 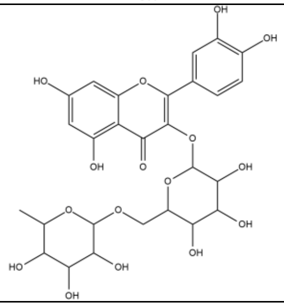 |
| 70 | Hymenidin | 6439099 | C11H12BrN5O | 310.15 | 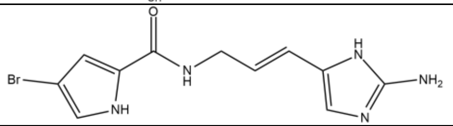 |

**Supplementary Table 1B:** List of synthetic ligands.

| S. No. | Synthetic Compound Name | PUBCHEM ID | Molecular Formula | Molecular Weight (g/mol) | Structure |
|---|---|---|---|---|---|
| 1 | MeBIO | 135524908 | C17H12BrN3O2 | 370.2 | 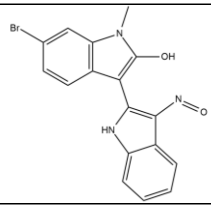 |
| 2 | CHIR-98014 | 53396311 | C20H17Cl2N9O2 | 486.3 | 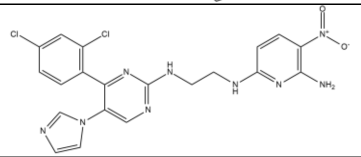 |
| 3 | ALOISINE A | 448912 | C16H17N3O | 267.33 | 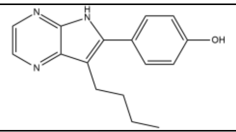 |
| 4 | SB 216763 | 176158 | C19H12Cl2N2O2 | 371.2 | 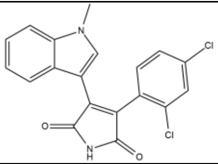 |
| 5 | CHIR-99021 | 9956119 | C22H18Cl2N8 | 465.3 | 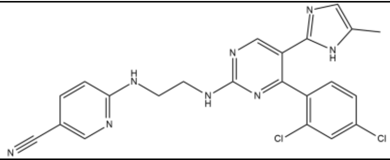 |
| 6 | Azithromycin | 447043 | C38H72N2O12 | 749.0 | 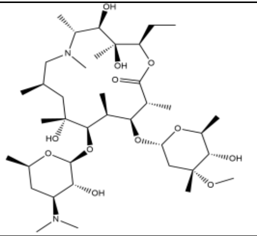 |
