## Supplementary material for "Natural compound screening predicts novel GSK-3 isoform-specific inhibitors": Suppl. Table 1C-D

**Supplementary table 1C:** Binding energies (kcal/mol) and dissociation constants (Kd) of natural compounds towards GSK-3α using AutoDock version 4.2.6.

| S.No. | Protein | Ligand | Binding Energy (kcal/mol) | Dissociation constant $(K_{d})$ | Interacting amino acid | Structure |
|---|---|---|---|---|---|---|
| 1 | GSK-3α | Berberine | -6.2 | 28.66 µM | Glu196, Leu195, Ala146, Lys148, Leu251, Cys262, As263, Asn249, Gln248, Asp244, Lys246 | 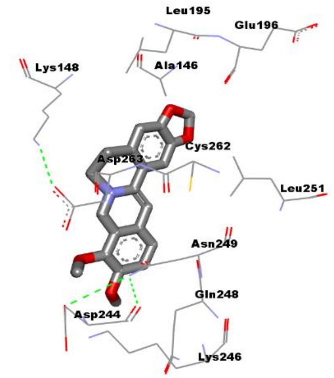 |
| 2 | GSK-3α | Rosmarinic acid | -4.84 | 283.34 µM | Phe130, Lys148, Glu160, Leu195, Gly2565, Asp263, Val173, Cys262, Asn249, Lys246, Ser129 | 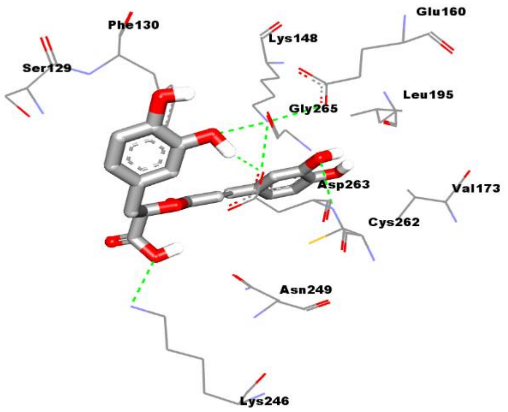 |

| | | | | | | |
|---|---|---|---|---|---|---|
| 3 | GSK-3α | Myricetin | **-4.71** | **350.4 µM** | Ser129, Gly128, Phe130, Asp263, Lys148, Cys262, Gln248, Asn249, Lys246 | 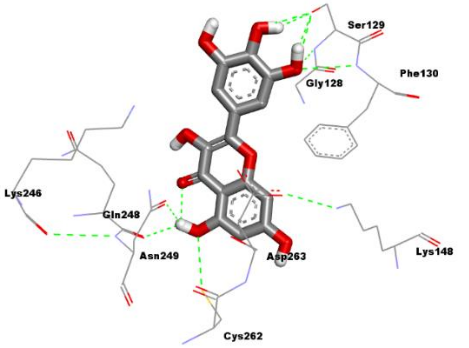 |
| 4 | GSK-3α | Honokiol | **-4.85** | **277.33 µM** | Phe130, Val133, Lys148, Leu195, Glu196, Val173, Asp263, Cys262, Leu261, Asn249, Gln248, Lys246 | 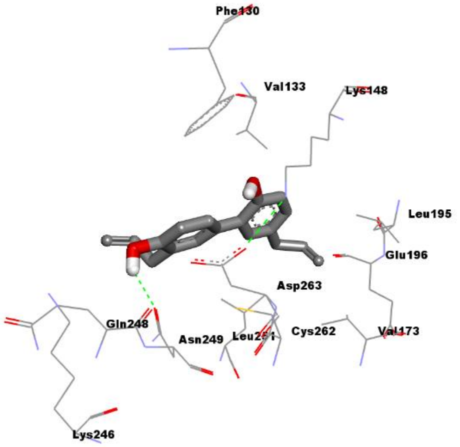 |
| 5 | GSK-3α | Luteolin | **-5.66** | **70.86 µM** | Gln248, Asn249, Asp263, Cys262, Val173, Lys148, Lau251, Phe130, Ala146, Tyr197, Glu196, Leu195, Val173 | 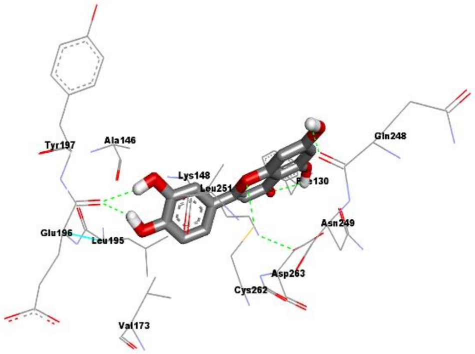 |

| | | | | | | |
|---|---|---|---|---|---|---|
| 6 | GSK-3α | Apigenin | **-5.69** | **67.68 µM** | Glu196, Val173, Leu251, Gln248, Asn249, Cys262, Asp263, Phe130, Lys148, Leu195, Ala146 | 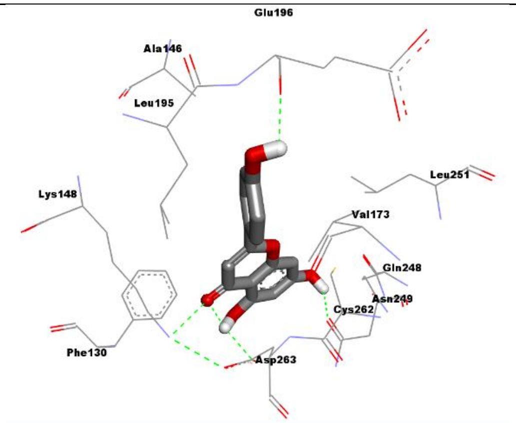 |
| 7 | GSK-3α | Caffeic acid | **-5.57** | **82.1 µM** | Lys148, Glu160, Phe264, Asp263, Cys262, Met164, Leu251, Val173, Val198, Tyr197, Glu196, Leu195, Ala146 | 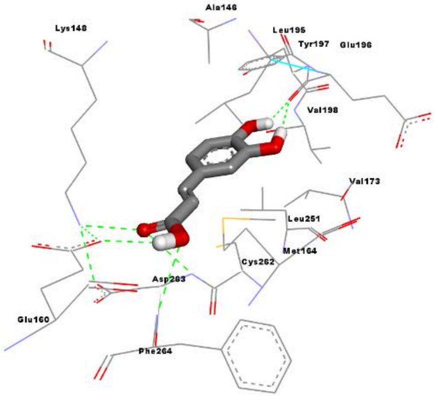 |
| 8 | GSK-3α | Daidzein | **-5.81** | **54.73 µM** | Lys148, Leu195, Glu196, Val173, Leu251, Cys262, Asn249, Asp263, Phe130 | 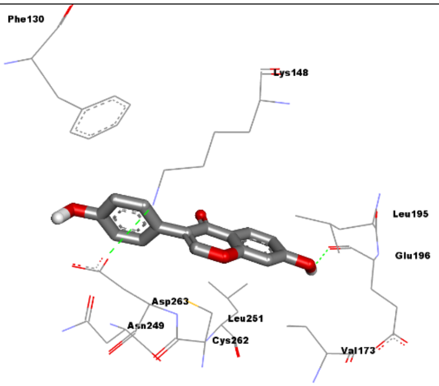 |

| | | | | | | |
|---|---|---|---|---|---|---|
| 9 | GSK-3α | Emodin | **-5.44** | **103.66 μM** | Lys148, Asp263, Leu195, Cys262, Glu196, Asn249, Leu251, Thr201, Gln248, Ala146 | 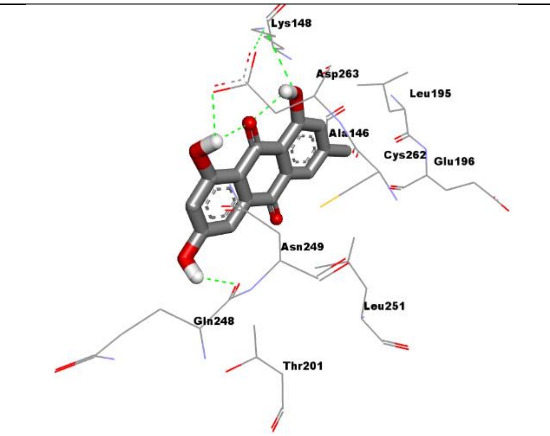 |
| 10 | GSK-3α | Galangin | **-6.37** | **21.43 μM** | Phe130, Glu160, Lys148, Phe264, Asp263, Cys262, Leu251, Val173, Leu195, Glu196, Val198, Tyr197, Ala146 | 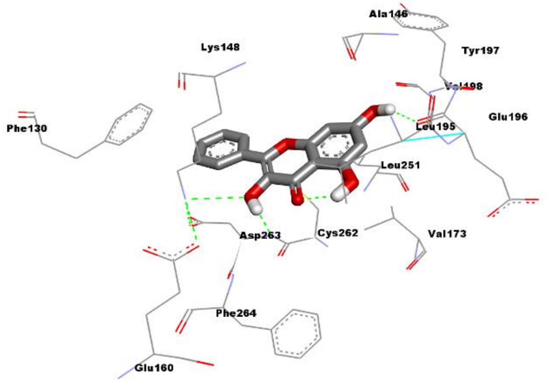 |
| 11 | GSK-3α | Oleanolic acid | **-3.78** | **1.68 mM** | Arg283, Ser282, Lys246, Asp244, Ser266, Asn249, Asp263, Gly265, Phe130, Ser129 | 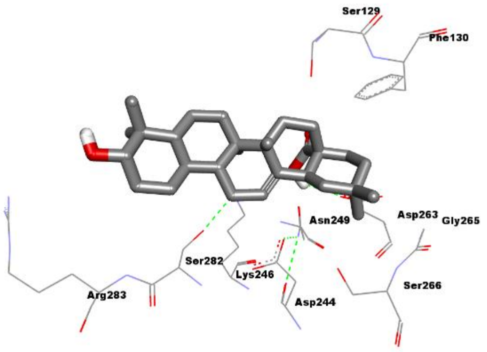 |

| | | | | | | |
|---|---|---|---|---|---|---|
| 12 | GSK-3α | Psoralidin | **-7.11** | **6.18 µM** | Lys148, Ala146, Tyr197, Val198, Leu195, Glu196, Leu251, Asp163, Cys262, Gln248, Phe130 | 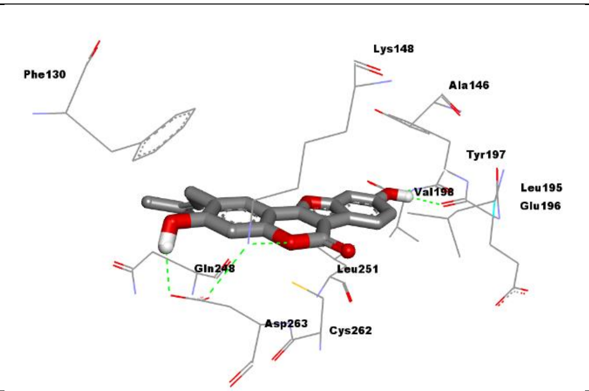 |
| 13 | GSK-3α | Beta-caryophyllene | **-5.65** | **71.7 µM** | Ala146, Leu195, Leu251, Asn249, Asp263, Cys262, Val173, Lys148, Val133, Phe130 | 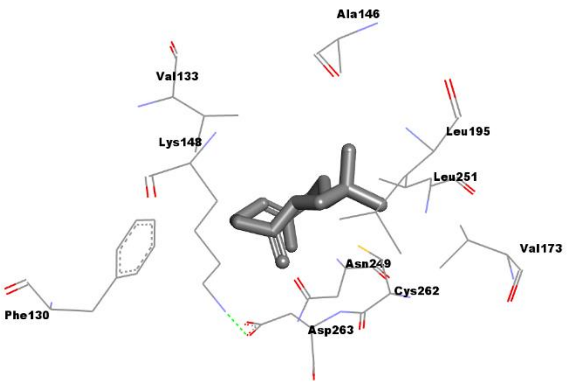 |
| 14 | GSK-3α | Catechin | **-4.88** | **263.1 µM** | Asp244, Lys246, Gln248, Asn249, Cys262, Leu251, Asp263, Lys148, Leu195, Glu160 | 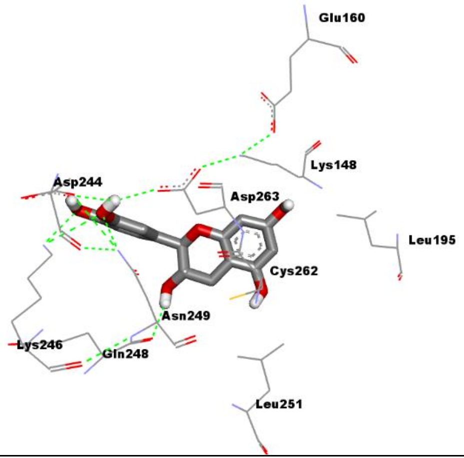 |

| | | | | | | |
|---|---|---|---|---|---|---|
| 15 | GSK-3α | Crocetin | **-4.58** | **442.27 µM** | Phe130, Val133, Lys148, Asp263, Asn249, Cys262, Ser266, Ser282, Cys281, Arg286, Arg283, Tyr279, Asp244, Lys246 | 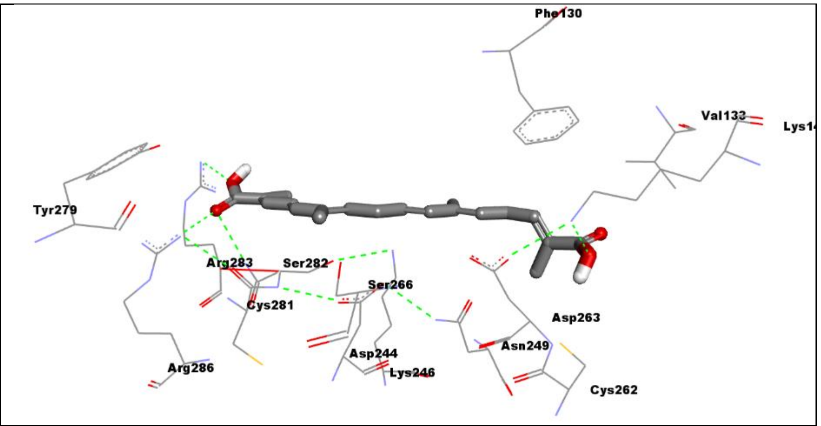 |
| 16 | GSK-3α | Piperine | **-6.29** | **24.4 µM** | Phe130, Val133, Lys148, Leu195, Glu196, Val173, Asp263, Cys262, Leu251, Asn249, Lys246 | 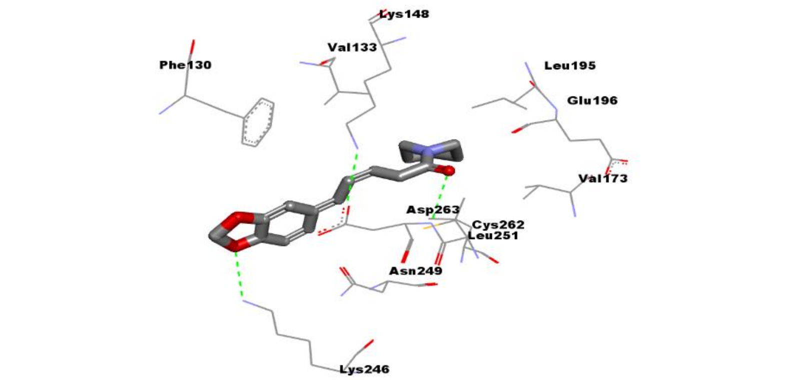 |
| 17 | GSK-3α | Zingibere e | **-5.70** | **65.99 µM** | Lys148, Glu160, Ala146, Leu195, Tyr197, Glu196, Val198, Val173, Cys262, Leu251, Asn249, Gln248, Asp263 | 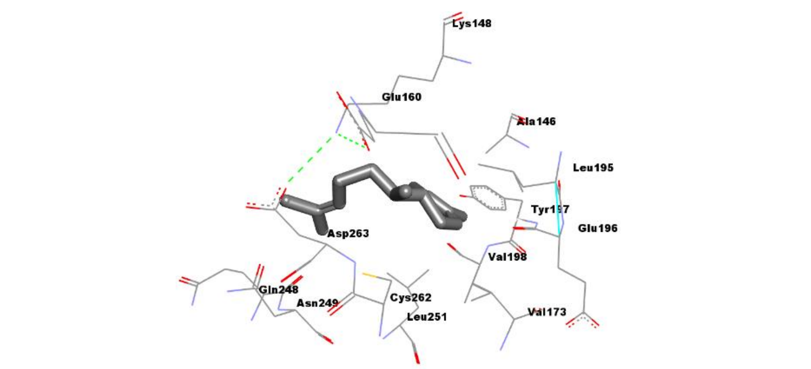 |

| | | | | | | |
|---|---|---|---|---|---|---|
| 18 | GSK-3α | turmerone | **-516** | **164.93 µM** | Phe130, Lys148, Glu160, Leu195, Glu196, Val173, Cys262, Asn249, Gln248, Asp263 | 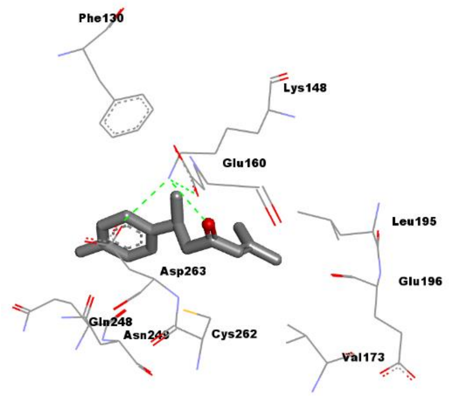 |
| 19 | GSK-3α | Carbazole | **-5.21** | **151.26 µM** | Leu251, Val173, Glu196, Ala146, Leu195, Lys148, Met164, Glu160, Asp263, Cys262, Phe264 | 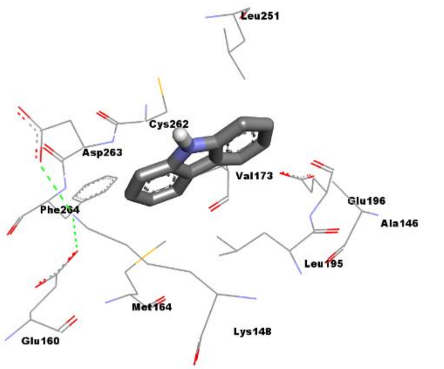 |
| 20 | GSK-3α | Carindone | **-2.27** | **21.85 mM** | Glu160, Gly265, Ser266, Asp263, Asp244, Lys246, Ser282, Lys148, Phe130, Ser129 | 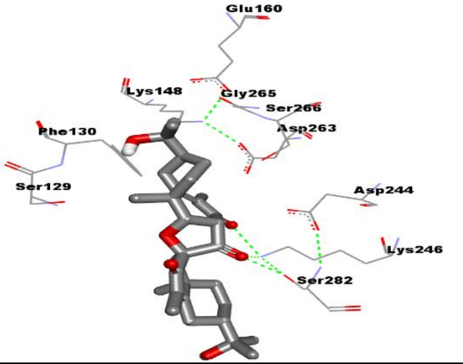 |

| | | | | | | |
|---|---|---|---|---|---|---|
| 21 | GSK-3α | Sesamin | **-6.47** | **18.02 µM** | Lys246, Asp244, Asn249, Leu251, Cys262, Asp263, Val173, Glu196, Leu195, Ala146, Lys148, Val133, Asn127, Phe130 | 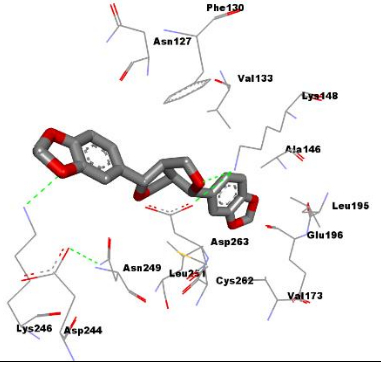 |
| 22 | GSK-3α | Moringin | **-4.58** | **440.4 µM** | Asn127, Gly128, Phe130, Ser129, Asn249, Asp263, Lys246, Asp244, Leu195, Lys148 | 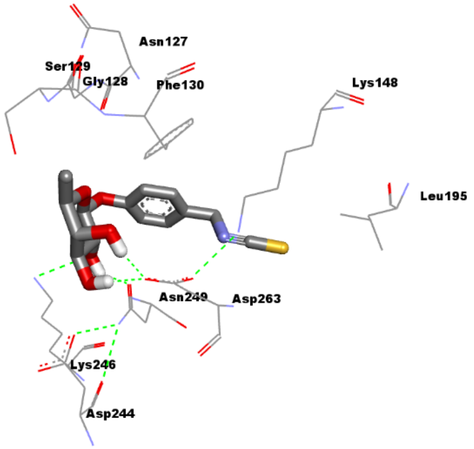 |
| 23 | GSK-3α | Quercetin | **-4.94** | **241.29 µM** | Leu195, Ala146, Glu196, Tyr197, Val198, Cys262, Leu251, Thr201, Gln248, Asp263, Phe130 | 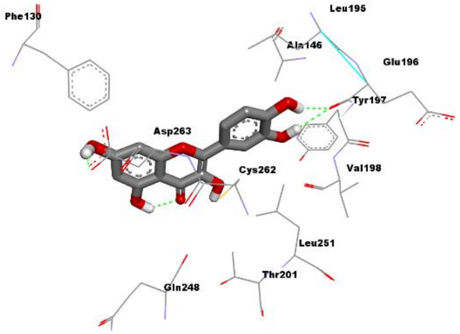 |

| | | | | | | |
|---|---|---|---|---|---|---|
| 24 | GSK-3α | Resveratrol | **-5.12** | **176.52 μM** | Phe130, Gln248, Asn249, Asp263, Cys262, Leu251, Val173, Glu196, Leu195, Ala146 | 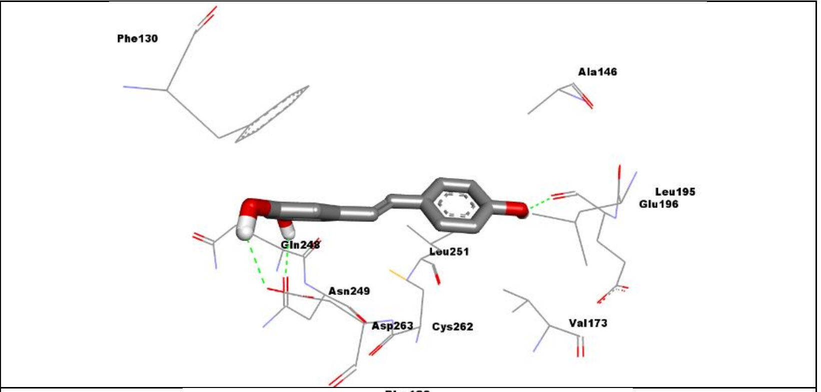 |
| 25 | GSK-3α | Kaempferol | **-5.47** | **98.58 μM** | Phe130, Lys148, Leu195, Glu196, Val173, Cys262, Asp263, Asn249, Thr201, Leu251, Gln248 | 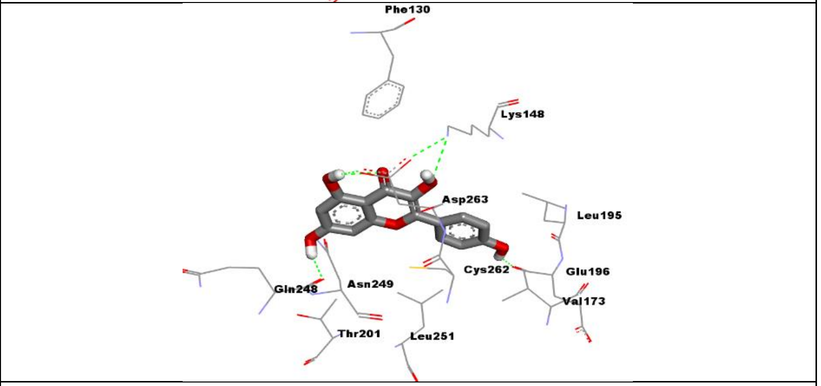 |
| 26 | GSK-3α | Costunolide | **-5.54** | **86.65 μM** | Thr201, Leu251, Cys262, Asp263, Leu195, Gln248, Val133, Phe130, Lys148 | 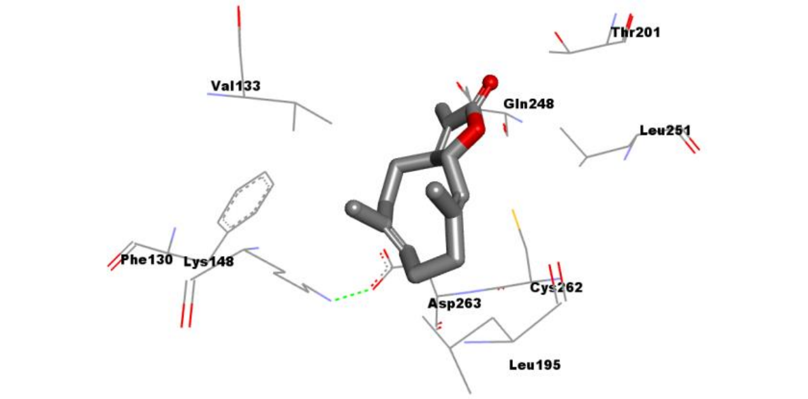 |

| | | | | | | |
|---|---|---|---|---|---|---|
| 27 | GSK-3α | Kumatakenin | **-5.27** | **137.77 µM** | Lys148, Leu195, Ala146, Glu196, Asp263, Cys262, Asn249, Gln248, Thr201, Leu251 | 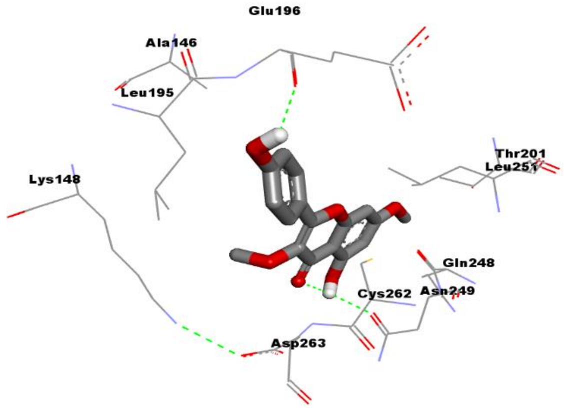 |
| 28 | GSK-3α | Withaferin A | **-6.47** | **18.14 µM** | Ser129, Phe130, Gly265, Asp263, Ser266, Asp244, Gln248, Tyr285, Lys246, Ser282, Arg283, Tyr284 | 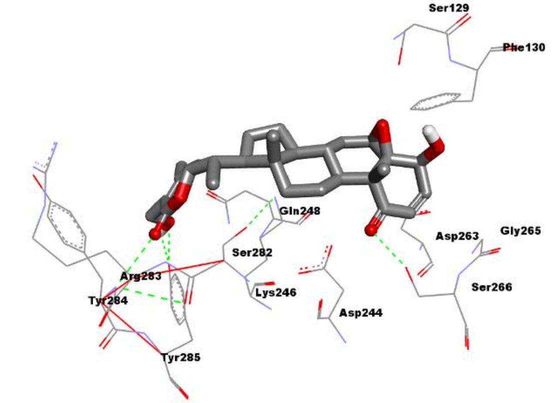 |
| 29 | GSK-3α | withanolide A | **-5.06** | **194.76 µM** | Ser129, Phe130, Gln248, Asp263, Asn249, Lys246, Asp244, Cys281, Arg283, Ser282 | 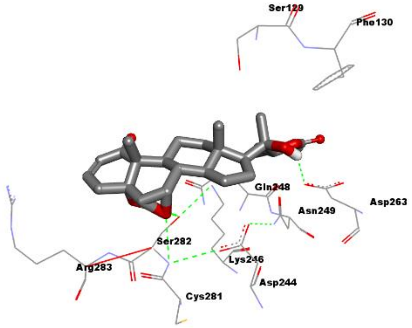 |

| | | | | | | |
|---|---|---|---|---|---|---|
| 30 | GSK-3α | Serpentine | **-6.13** | **31.86 µM** | Thr201, Leu251, Gln248, Asn249, Cys262, Lys246, Asp244, Asp263, Phe130, Lys148, Leu195 | 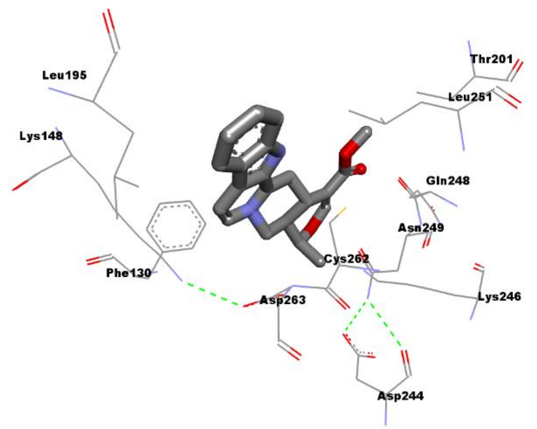 |
| 31 | GSK-3α | Arjunolic acid | -4.04 | 1.09 mM | Ser282, Lys246, Asp244, Ser266, Gly265, Asp263, Phe130, Ser129, Lys148 | 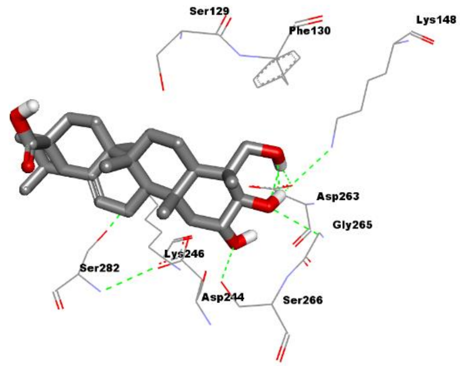 |
| 32 | GSK-3α | Betulinic acid | -4.68 | 372.97 µM | Ser129, Gly128, Phe130, Asn249, Asp263, Ser266, Asp244, Cys281, Arg286, Arg283, Ser282, Lys246 | 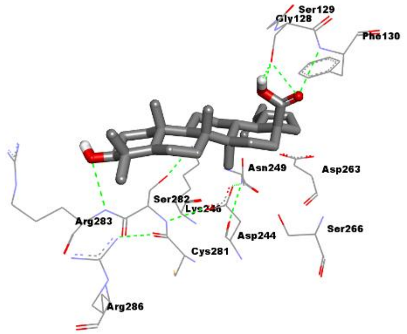 |

| | | | | | | |
|---|---|---|---|---|---|---|
| 33 | GSK-3α | Cinnamic acid | -5.29 | 132.98 µM | Lys148, Ala146, Leu195, Glu196, Val173, Glu160, Met164, Phe264, Cys262, Asp263, Leu251 | 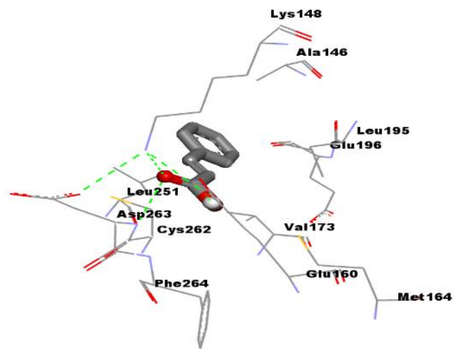 |
| 34 | GSK-3α | Ginsenosides | -3.71 | 1.89 mM | Phe130, Ser129, Gly128, Gly265, Asp263, Gln248, Asp244, Lys246, Ser282, Arg283, Tyr284, Tyr285 | 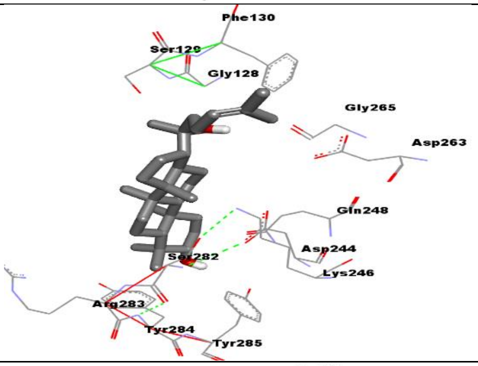 |
| 35 | GSK-3α | Morphine | -4.11 | 977.09 µM | Ser129, Phe130, Ser282, Lys246, Asp263, Gly265, Ser266, Asp244 | 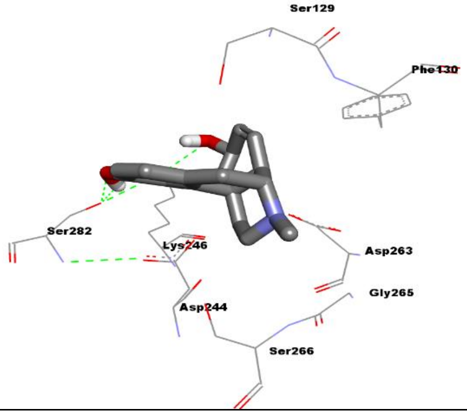 |

| | | | | | | |
|---|---|---|---|---|---|---|
| 36 | GSK-3α | Thymoquinone | -5.39 | 112.78 μM | Glu196, Lys148, Leu195, Leu251, Val173, Cys262, Asp263, Phe264, Met164, Glu160 | 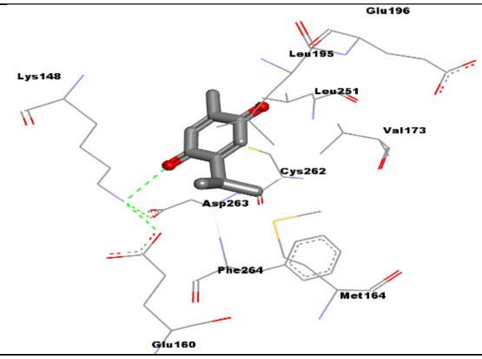 |
| 37 | GSK-3α | Curcumin | -4.75 | 332.17 μM | Ser129, Phe130, Val133, Ala146, Tyr197, Leu195, Glu196, Val198, Leu251, Asp263, Cys262, Val173, Asn249 | 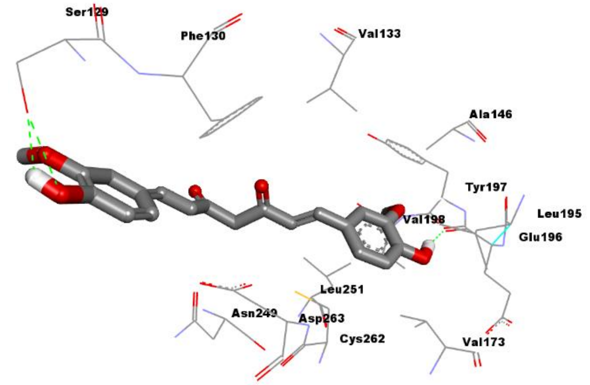 |
| 38 | GSK-3α | Mangiferin | -3.09 | 5.43 mM | Lys148, Leu195, Asp263, Asn249, Leu251, Cys262, Lys246, Asp244, Ser282, Ser129, Phe130, Gly128 | 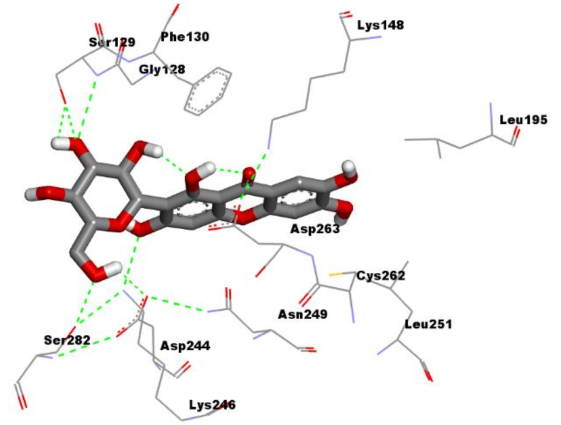 |

| | | | | | | |
|---|---|---|---|---|---|---|
| 39 | GSK-3α | Phloretin | -4.54 | 467.86 μM | Gln248, Lys246, Asn249, Leu251, Asp263, Cys262, Glu196, Leu195, Ala146, Lys148, Val133 | 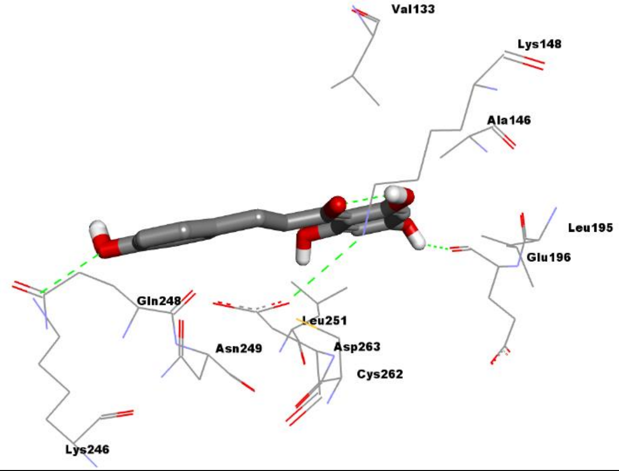 |
| 40 | GSK-3α | Hymenidin | -4.51 | 498.39 μM | Phe130, Ser266, Gly265, Asp244, Asp263, Lys246, Cys262, Asn249, Gln248 | 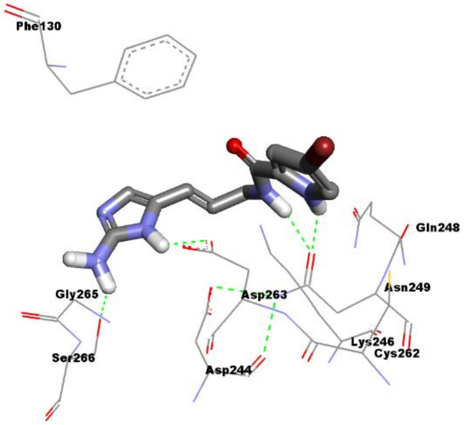 |

**Supplementary Table 1D;** Binding energies (kcal/mol) and dissociation constants (Kd) of natural compounds towards GSK-3β using AutoDock version 4.2.6.

| **S.No.** | **Protein** | **Ligand** | **Binding Energy (kcal/mol)** | **Dissociation constant ($K_d$)** | **Interacting amino acid** | **Structure** |
|---|---|---|---|---|---|---|
| 1 | GSK-3β | Berberine | **-8.65** | **452.71 nM** | Gln265, Gly259, Gly262, Tyr221, Arg223, Arg220, Tyr216, Asp260, Ile228, Ser261 | 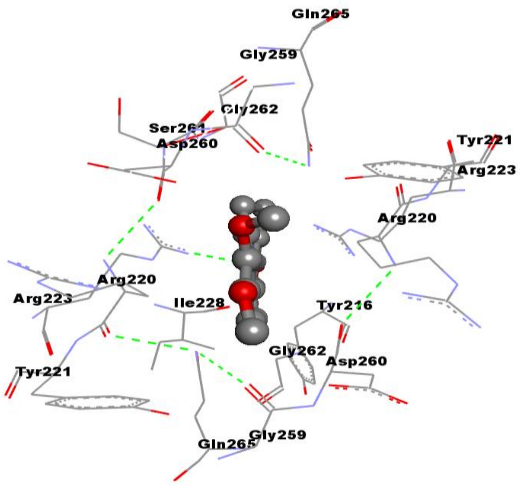 |

| | | | | | | |
|---|---|---|---|---|---|---|
| 2 | GSK-3β | Rosmarinic acid | **-9.71** | **76.64 nM** | Ile217, Tyr216, Cys218, Gly262, Ser261, Ser219, Gly259, Asp260, Tyr221, Arg220, Arg223, Gln265 | 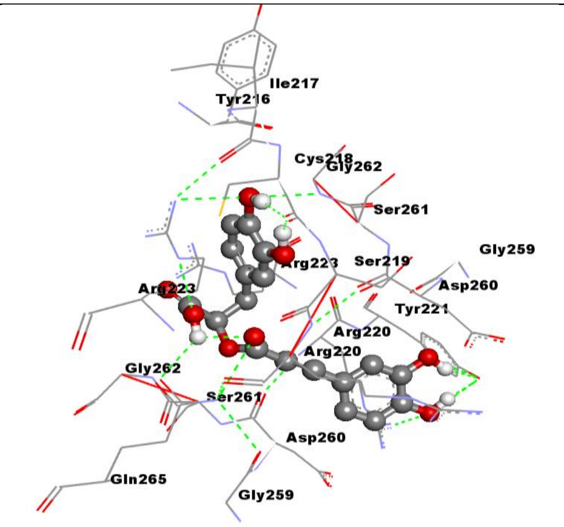  |
| 3 | GSK-3β | Myricetin | **-7.57** | **2.83 µM** | Tyr221, Arg223, Ile228, Arg220, Ser261, Asp260, Gly262, Gly259, Gln265 | 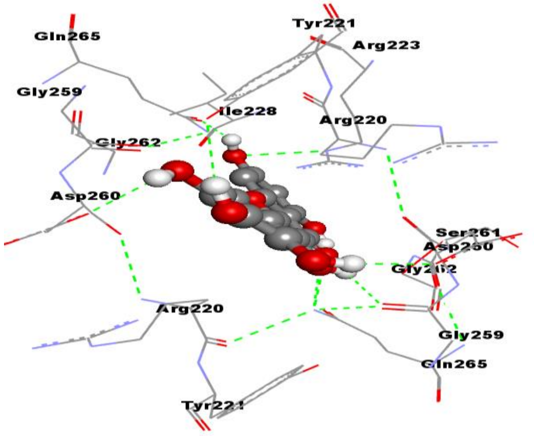  |

| | | | | | | |
|---|---|---|---|---|---|---|
| 4 | GSK-3β | Honokiol | **-7.0** | **7.4 µM** | Gln265, Gly262, Ser215, Arg223, Leu227, Ile228, Gly230, Ser261, Cys218, Asp260 | 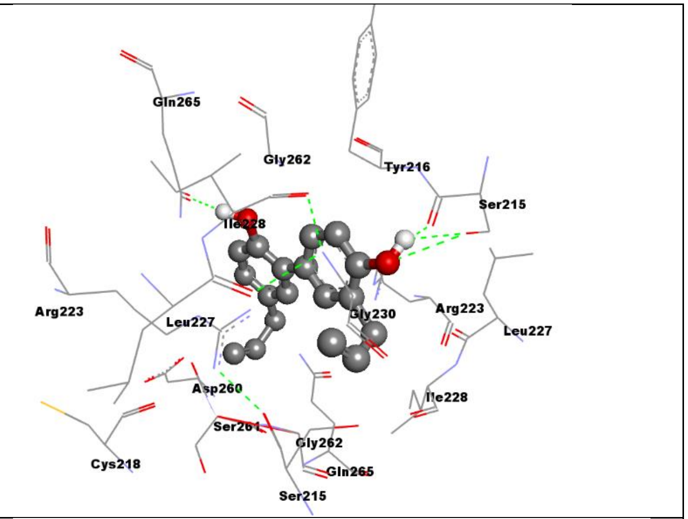 |
| 5 | GSK-3β | Luteolin | **-8.80** | **353.7 nM** | Gln265,Arg223, Gly259, Asp260, Arg220, Ser261, Gly262, Cys218, Ile228, Ile217, Tyr216 | 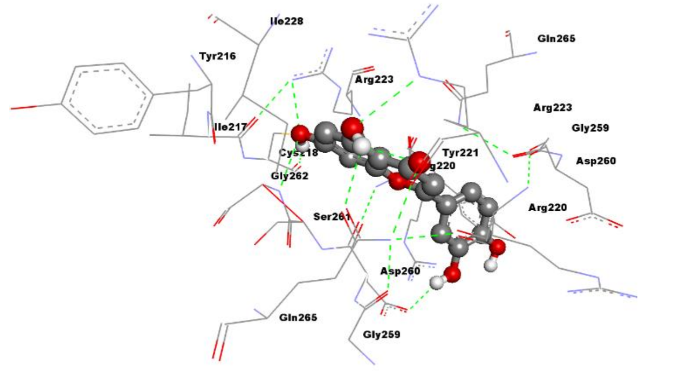 |

| | | | | | | |
|---|---|---|---|---|---|---|
| 6 | GSK-3β | Apigenin | **-9.16** | **194.29 nM** | Gln265, Arg223, Tyr221, Gly259, Asp260, Arg220, Ser261, Gly262, Ile217, Tyr216, Cys218, Ile228 | 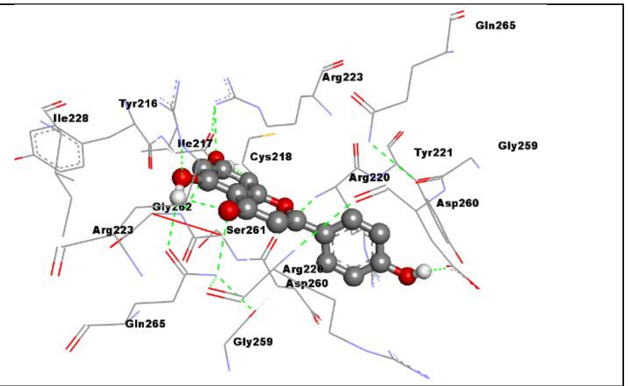 |
| 7 | GSK-3β | Caffeic acid | **-6.55** | **15.85 µM** | Ile217, Cys218, Gly262, Ser261, Ser219, Asp260, Arg220, Ser219, Arg223, Tyr216 | 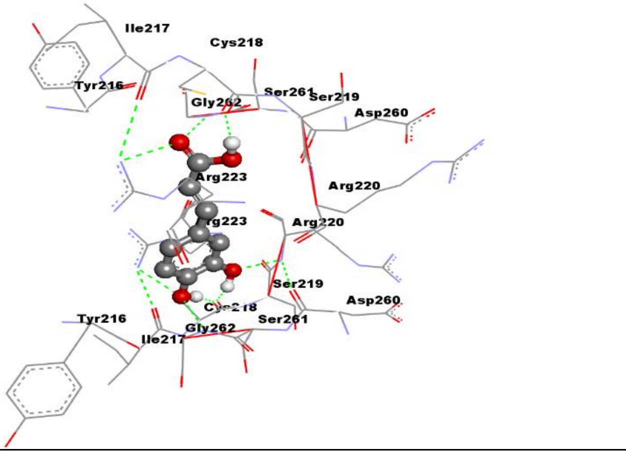 |

| | | | | | | |
|---|---|---|---|---|---|---|
| 8 | GSK-3β | Daidzein | **-7.78** | **1.99 µM** | Gln265, Gly259, Asp260, Tyr221, Arg220, Ser261, Gly262, Ile228, Arg223 | 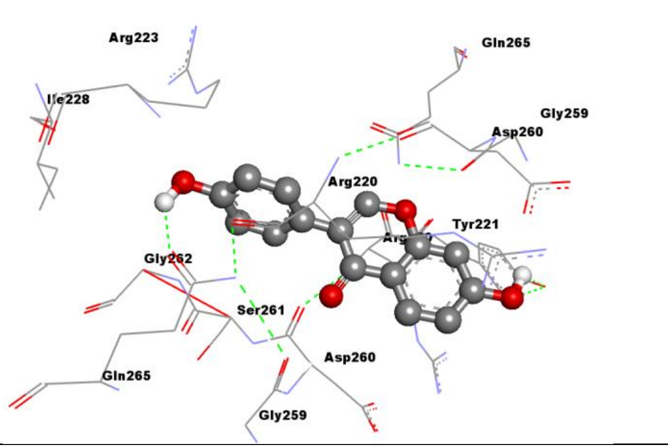  |
| 9 | GSK-3β | Emodin | **-7.27** | **4.69 µM** | Gly262, Gln265, Ser261, Gly259, Asp260, Tyr221, Arg220, Arg223 | 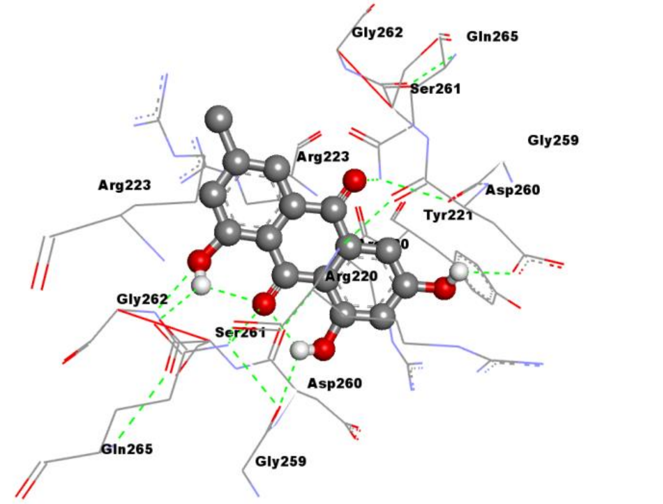  |

| | | | | | | |
|---|---|---|---|---|---|---|
| 10 | GSK-3β | Galangin | **-7.48** | **3.28 μM** | Gly262, Gln265, Ser261, Gly259, Asp260, Arg223, Tyr221, Arg220, Ser261 | 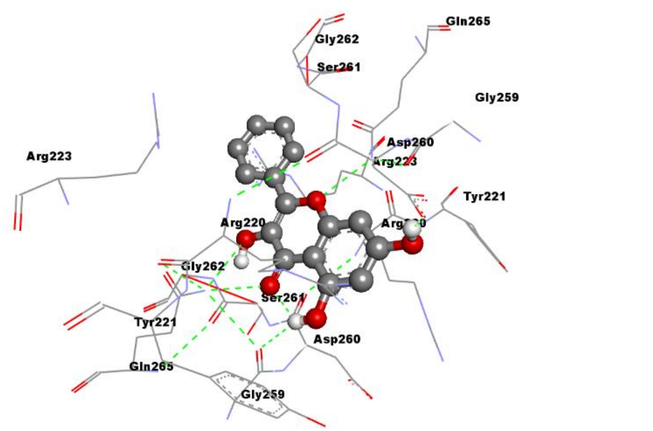 |
| 11 | GSK-3β | Oleanolic acid | **-7.39** | **3.82 μM** | Ser147, Arg144, Tyr140, Gly253, Gln254, Glu249, Pro255, Tyr221, Arg220, Asp260 | 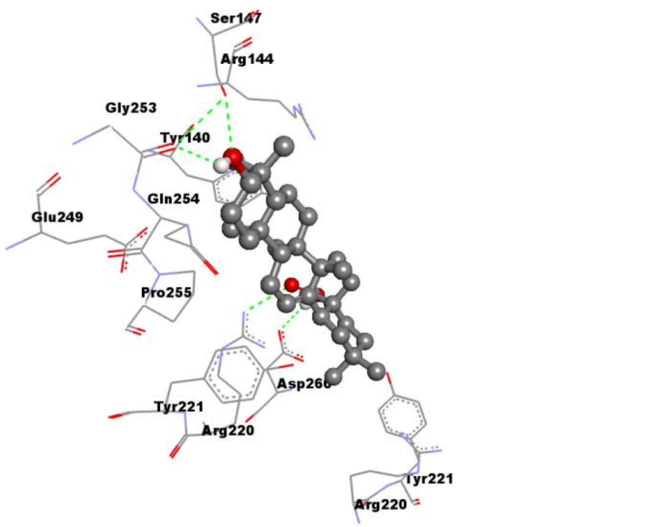 |

| | | | | | | |
|---|---|---|---|---|---|---|
| 12 | GSK-3β | Psoralidin | **-8.43** | **661.74 nM** | Arg223, Ser261, Tyr221, Gly262, Gln265, Asp260, Gly259, Arg220, Ile228 | 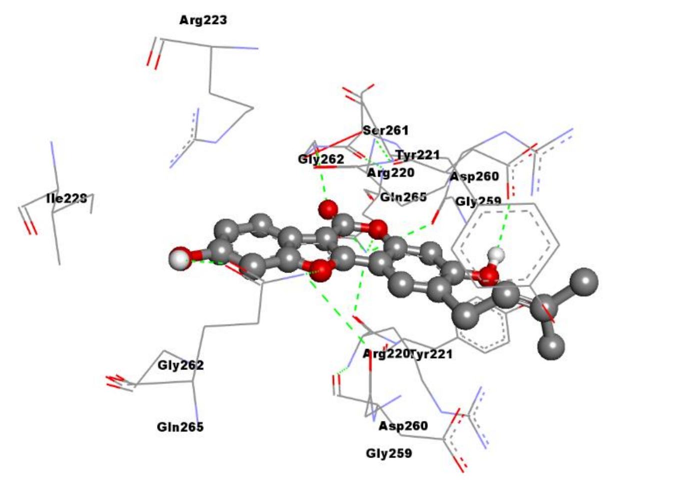 |
| 13 | GSK-3β | Beta-caryophylle ne | **-7.22** | **5.14 µM** | Tyr216, Ser261, Gly262, Gln265, Ile228, Arg223, Arg220, Cys218, Asp260 | 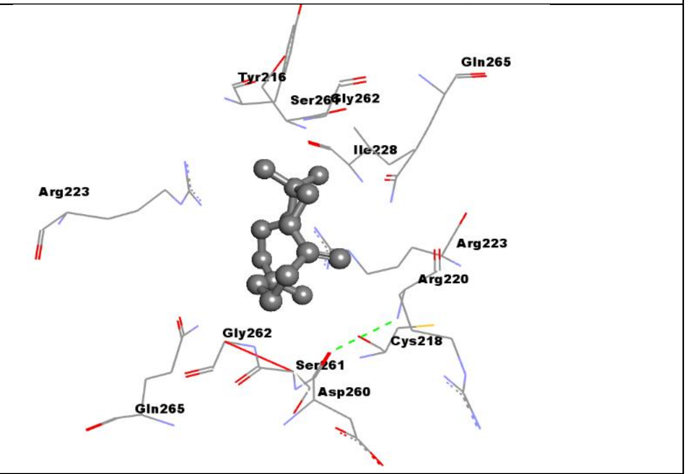 |

| | | | | | | |
|---|---|---|---|---|---|---|
| 14 | GSK-3β | Catechin | **-8.38** | **715.12 nM** | Arg223, Tyr221, Arg220, Asp260, Gly259, Gln265, Ser219, Cys218, Ile217, Gly262, Ser261, Tyr216, Ser215 | 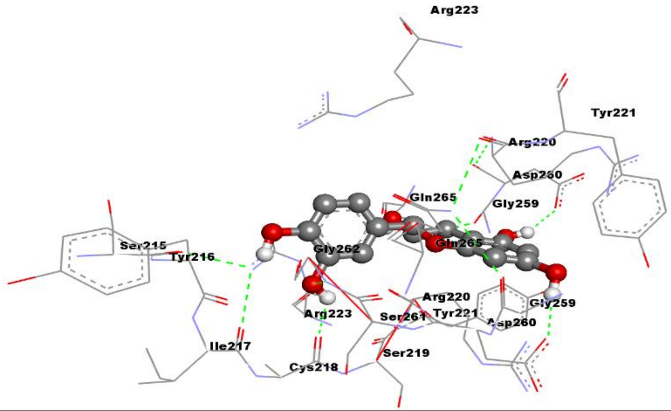 |
| 15 | GSK-3β | Crocetin | **-7.87** | **1.7 μM** | Ser215, Gly230, Ile228, Gly262, Ser261, Arg223, Gln265, Gly259, Tyr221, Asp260, Arg220, | 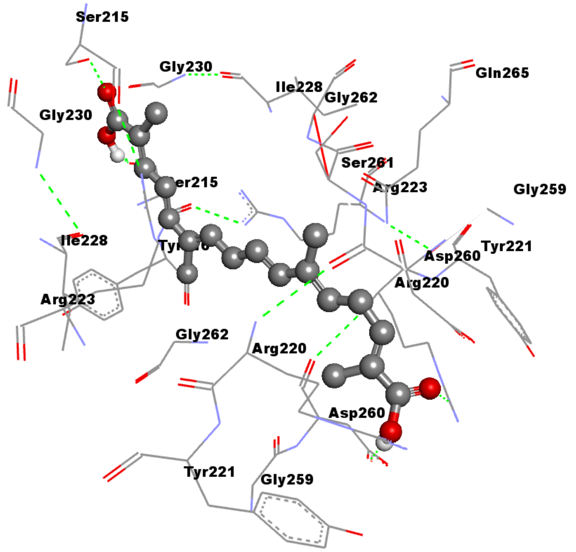 |

<table>
<tr><td></td><td></td><td></td><td></td><td></td><td></td><td></td></tr>
<tr><td>16</td><td>GSK-3β</td><td>Piperine</td><td>-8.48</td><td>608.02 nM</td><td>Arg223, Arg220, Tyr221, Asp260, Gly259, Cys218, Ser261, Gln265, Ile228, Tyr216, Gly262, Ser215</td><td>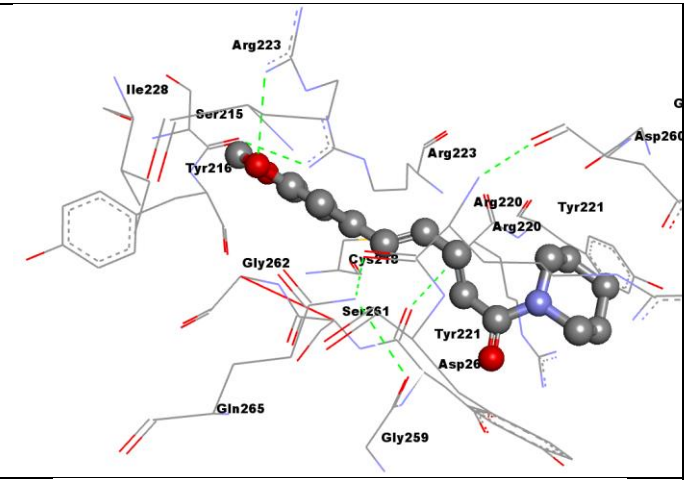
</td></tr>
<tr><td>17</td><td>GSK-3β</td><td>Zingiberene</td><td>-6.96</td><td>7.93 µM</td><td>Gln265, Gly259, Arg223, Asp260, Tyr221, Tyr216, Ser261, Gly262</td><td>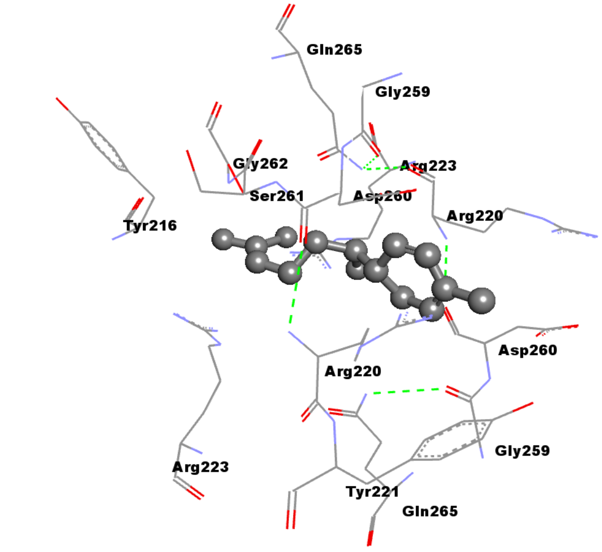
</td></tr>
</table>

| | | | | | | |
|---|---|---|---|---|---|---|
| 18 | GSK-3β | turmerone | **-6.84** | **9.67 μM** | Arg223, Gln265, Gly259, Asp260, Arg220, Gly262, Ile228, Tyr216 | 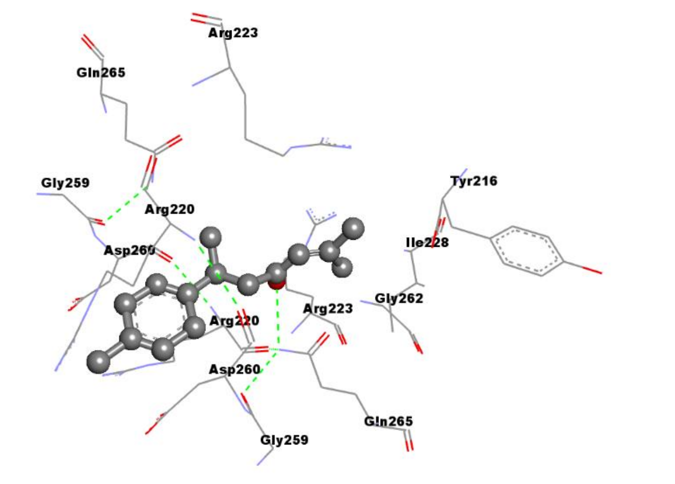 |
| 19 | GSK-3β | Carbazole | **-6.4** | **20.39 μM** | Gln265, Gly259, Asp260, Arg220, Ser261 | 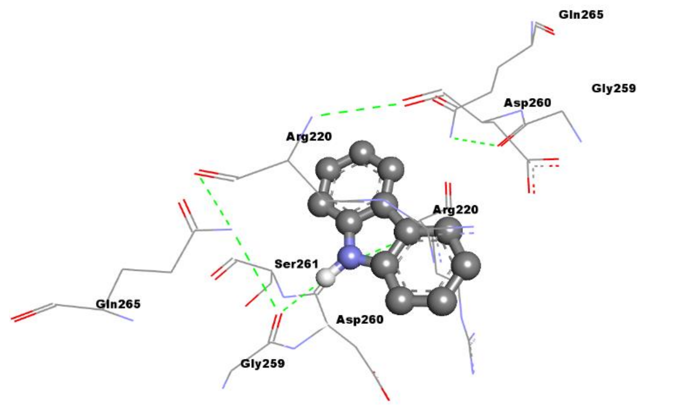 |

| | | | | | | |
|---|---|---|---|---|---|---|
| 20 | GSK-3β | Carindone | **-7.17** | **5.59 µM** | Tyr221, Tyr222, Pro255,Glu249, Arg220,Tyr140, Ala143, Arg144 | 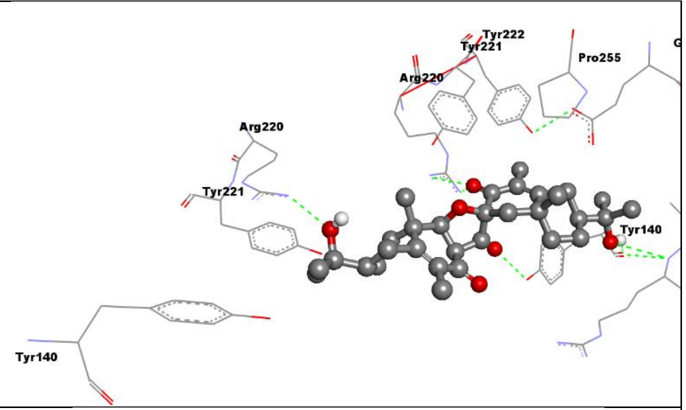 |
| 21 | GSK-3β | Sesamin | **-9.10** | **212.96 nM** | Ser215, Tyr216, Arg223, Leu227, Gln265, Gly259, Asp260, Arg220, Ser261, Gly262, Ile228, | 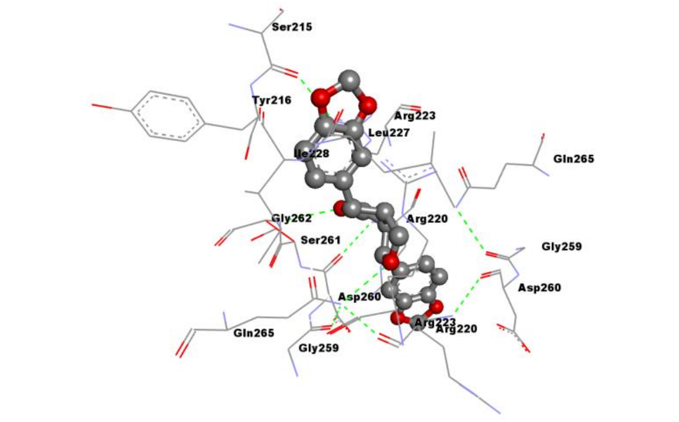 |

| | | | | | | |
|---|---|---|---|---|---|---|
| 22 | GSK-3β | Moringin | **-7.50** | **3.17 µM** | Ser215, Tyr216, Ile228, Arg223, Gly230, Arg220, Asp260, Gln265, Gly262, Ser261, Phe229, Leu227. Gly230 | 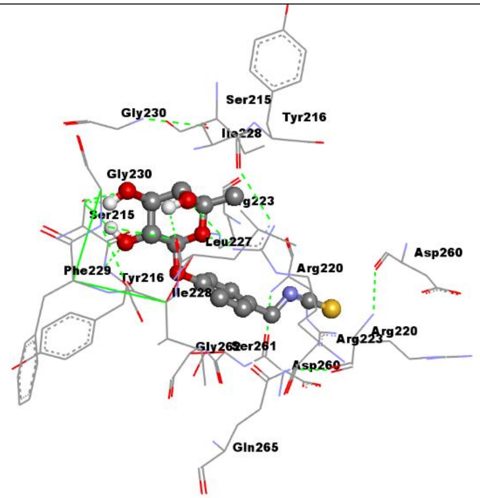 |
| 23 | GSK-3β | Quercetin | **-7.83** | **1.83 µM** | Gly262, Ser261, Gln265, Gly259, Asp260, Tyr221, Arg220, Arg223, Ile228 | 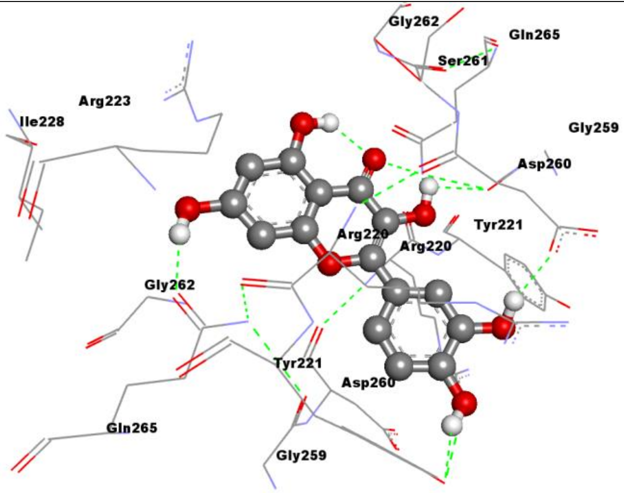 |

| | | | | | | |
|---|---|---|---|---|---|---|
| 24 | GSK-3β | Resveratrol | **-8.54** | **545.64 nM** | Tyr216, Ile228, Gln265, Arg223, Ile217, Cys218, Arg220, Tyr221, Gly259, Asp260, Ser261, Gly262 | 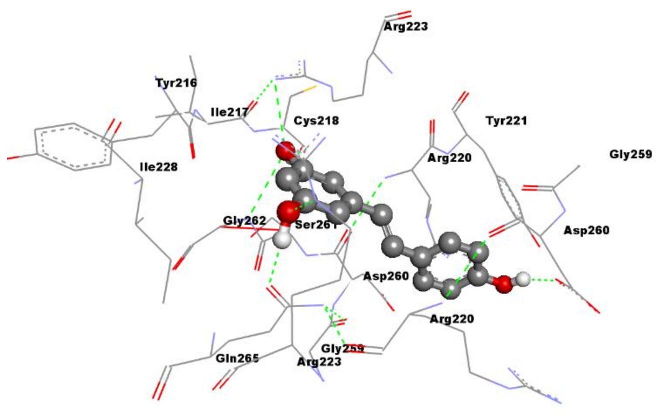 |
| 25 | GSK-3β | Kaempferol | **-7.77** | **2.81 µM** | Ile228, Gly262, Gln265, Gly259, Asp260, Tyr221, Arg220, Arg223, Ser261 | 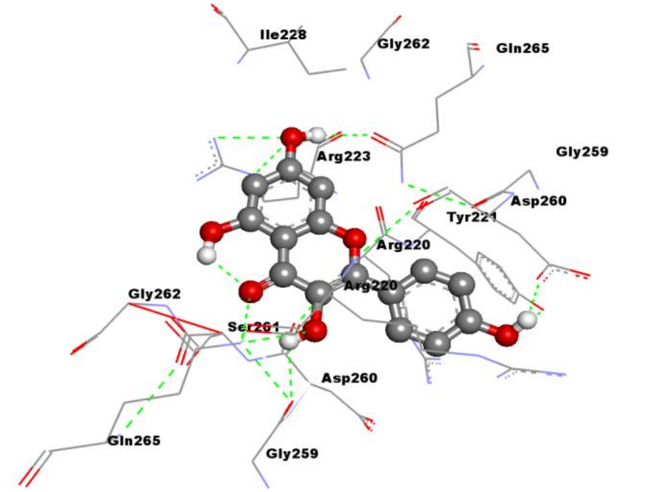 |

| | | | | | | |
|---|---|---|---|---|---|---|
| 26 | GSK-3β | Costunolide | **-7.92** | **1.57 µM** | Cys218, Ile217, Tyr216, Ser261, Asp260, Gly262, Gln265, Ile228, Arg220, Arg223, Ser261, Asp260, Gly259 | 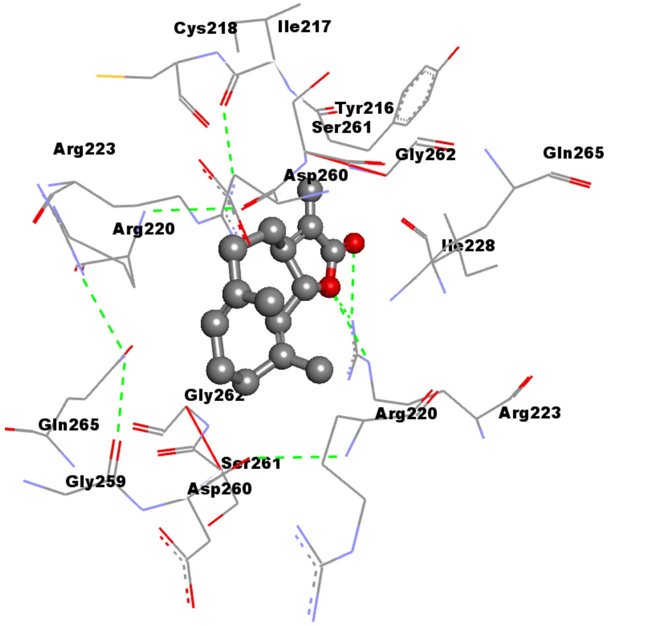  |
| 27 | GSK-3β | Kumatakeni n | **-7.54** | **2.99 µM** | Arg223, Gln265, Leu227, Tyr221, Gly259, Arg220, Gly262, Tyr216, Asp260, Ser261, | 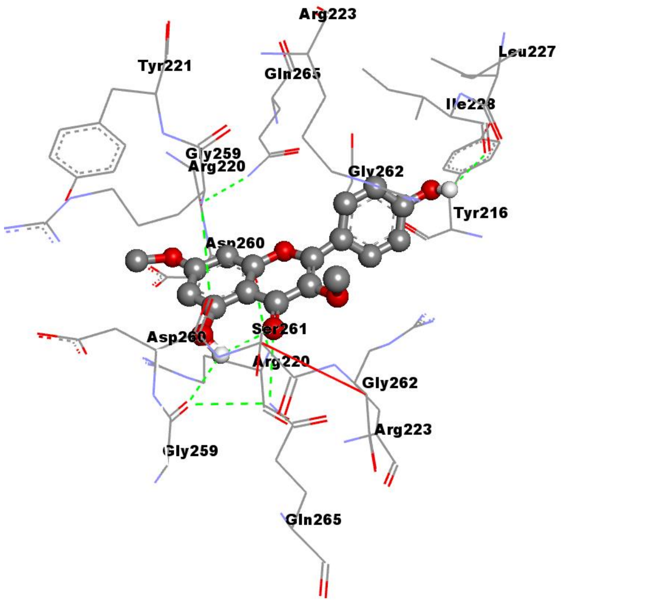  |

| | | | | | | |
|---|---|---|---|---|---|---|
| 28 | GSK-3β | Withaferin A | **-7.23** | **5.05 µM** | Cys218, Arg223, Tyr221, Arg220, Gly262, Gln265, Gly259, Ser261, Asp260, | 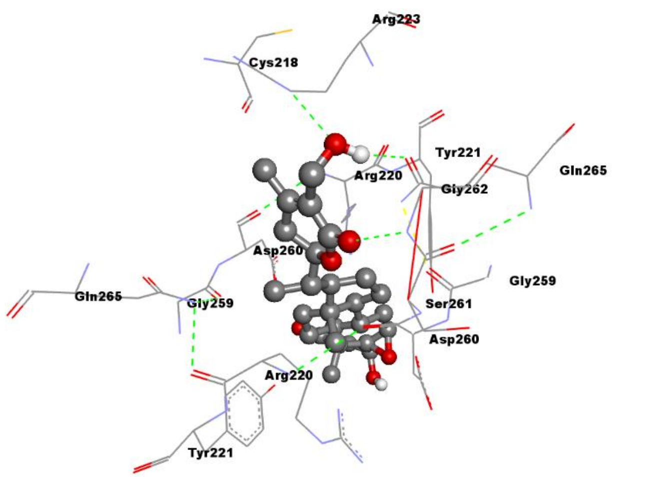 |
| 29 | GSK-3β | withanolide A | **-7.32** | **4.34 µM** | Gln265, Gly259, Asp260, Tyr221, Gln185, Tyr140, Gly259, Arg220, | 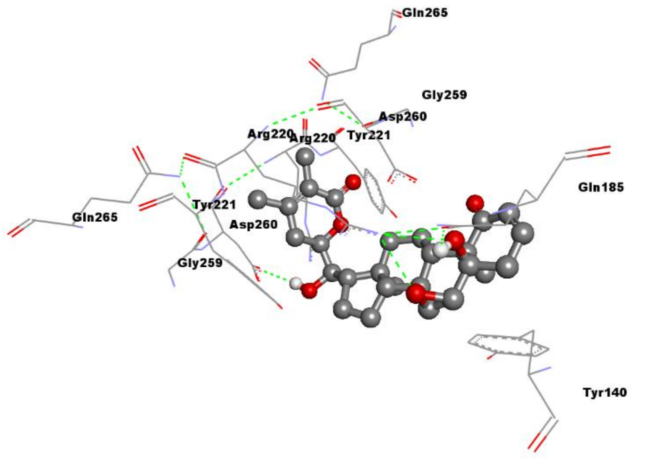 |

| | | | | | | |
|---|---|---|---|---|---|---|
| 30 | GSK-3β | Serpentine | **-8.39** | **704.95 nM** | Gly262, Gln265, Ser261, Cys218, Gly259, Arg220, Arg223, Tyr221, Asp260, Ile228 | 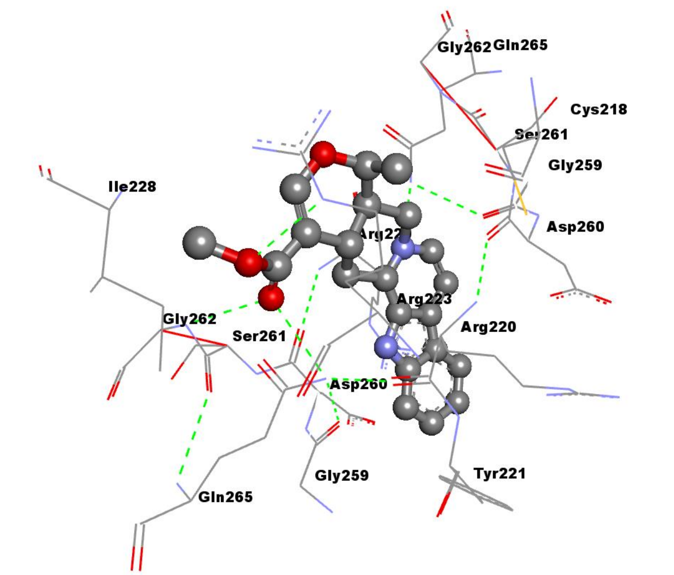 |
| 31 | GSK-3β | **Arjunolic acid** | **-6.95** | **7.98 µM** | Asp260, Tyr140, Tyr221, Arg220, Arg144, Pro255, Gln254, Gly253, Ser147 | 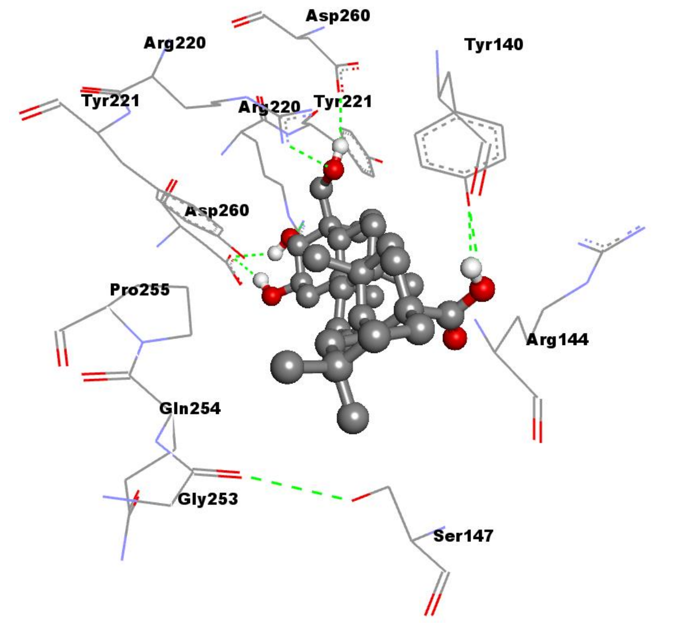 |

| | | | | | | |
|---|---|---|---|---|---|---|
| 32 | GSK-3β | **Betulinic acid** | **-6.33** | **23.06 µM** | Tyr221, Arg220, Asp260, Tyr140, Glu249, Pro255, Gln254, Gly253, Ser147, Arg144 | 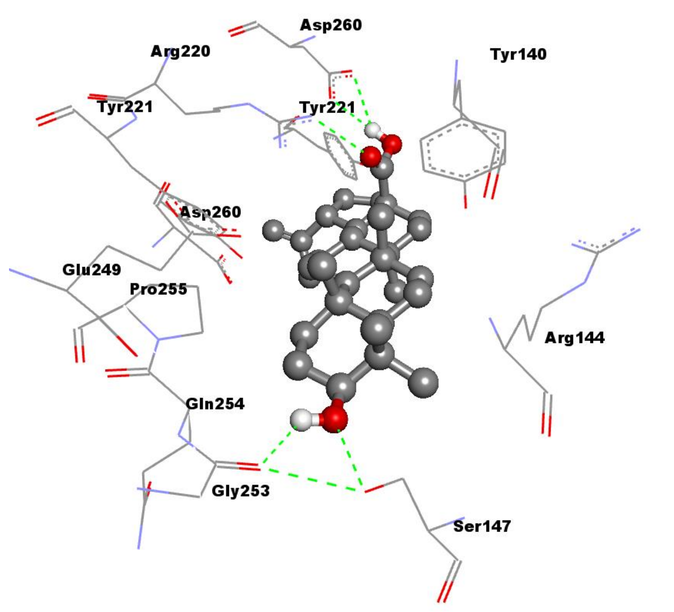 |
| 33 | GSK-3β | **Cinnamic acid** | **-6.08** | **35.23 µM** | Arg223, Ile228, Arg220, Gly262, Gln265, Asp260 | 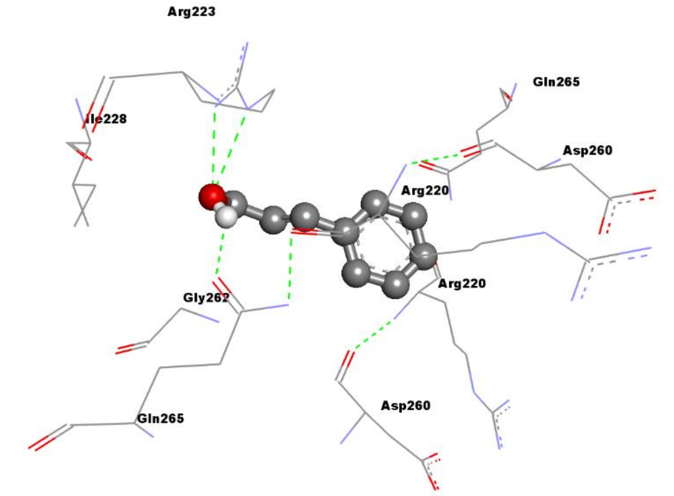 |

| | | | | | | |
|---|---|---|---|---|---|---|
| 34 | GSK-3β | **Ginsenosides** | **-6.12** | **32.87 µM** | Arg220, Tyr221, Pro255, Glu249, Tyr140, Ala143, Arg144, Asp260 | 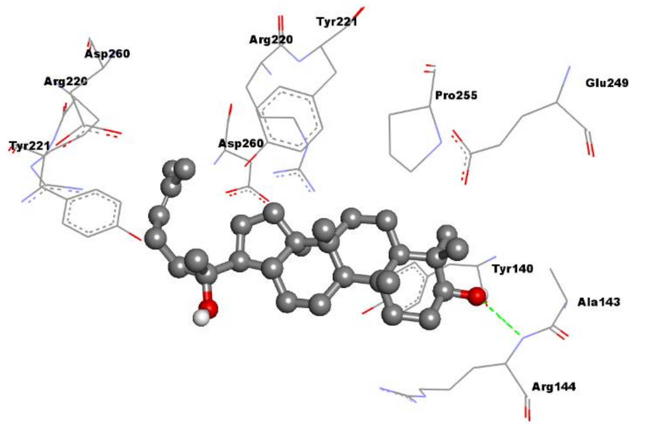 |
| 35 | GSK-3β | **Morphine** | **-6.80** | **10.41 µM** | Ser215, Tyr216, Cys218, Gly262, Gln265, Ser261, Arg223, Gly259, Asp260, Arg220 | 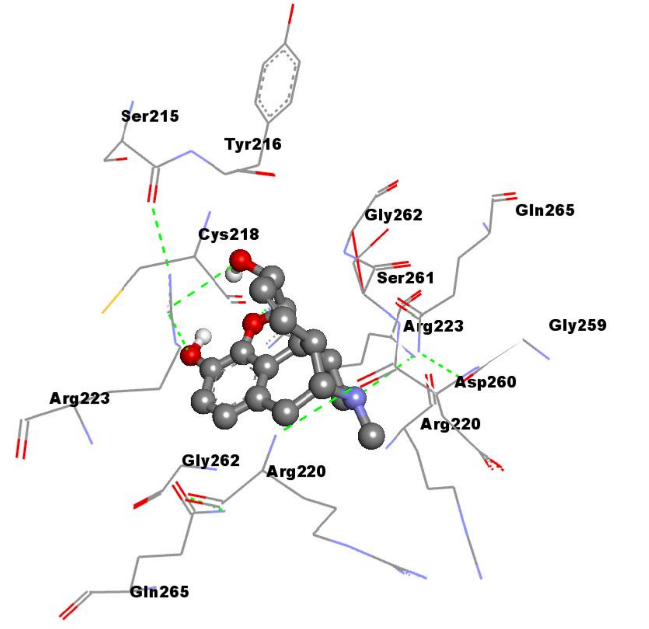 |

| | | | | | | |
|---|---|---|---|---|---|---|
| 36 | GSK-3β | **Thymoquinone** | **-6.13** | **31.98 µM** | Arg223, Arg220, Gln265, Asp260, Ile217, Cys218, Ser261, Gly259, Gly262, Tyr216 | 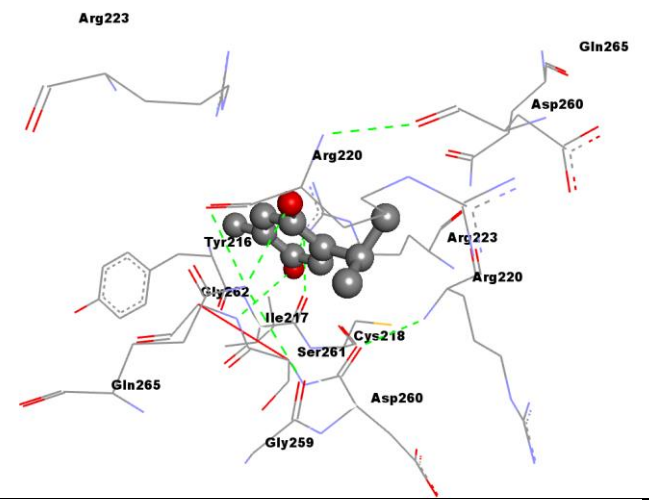 |
| 37 | GSK-3β | **Curcumin** | **-6.66** | **13.03 µM** | Ser215, Tyr216, Ile217, Cys218, Arg223, Gln265, Tyr221, Ser261, Asp260, Arg220, Gly259, Gly262, Ile228 | 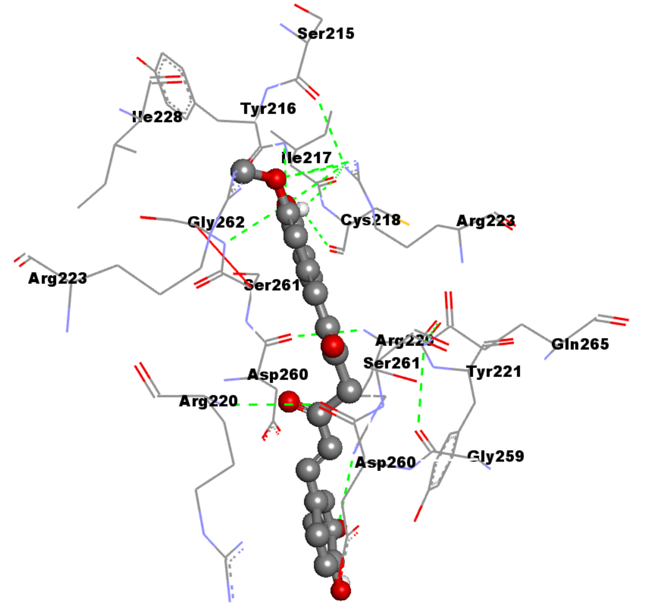 |

| | | | | | | |
|---|---|---|---|---|---|---|
| 38 | GSK-3β | **Mangiferin** | **-6.72** | **11.83 µM** | Gln265, Gly259, Asp260, Tyr221, Arg220, Arg223, Ser261, Gly262 | 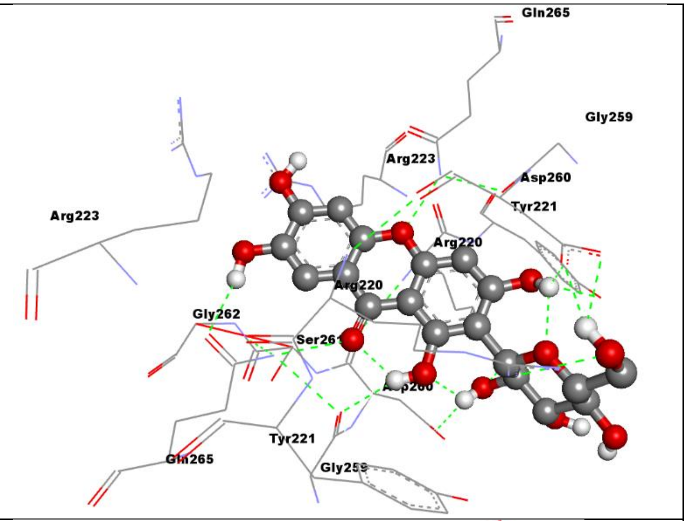  |
| 39 | GSK-3β | **Phloretin** | **-6.94** | **8.21 µM** | Gln265, Gly259, Arg223, Arg220, Asp260, Ser261, Tyr221, Gly262, Ile228 | 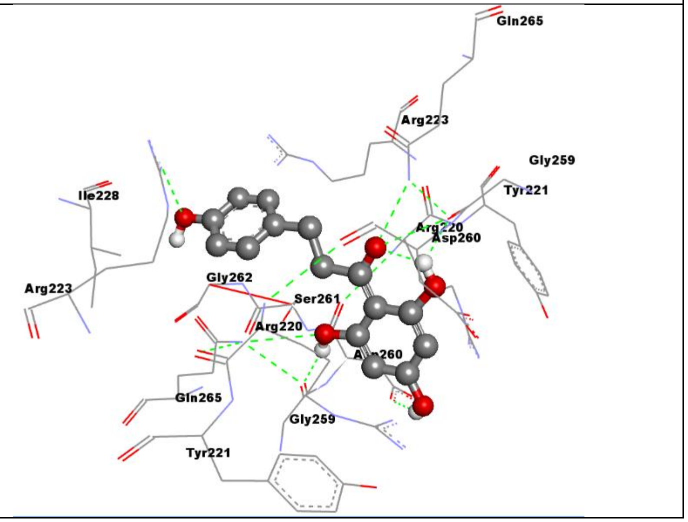  |

| | | | | | | |
|---|---|---|---|---|---|---|
| 40 | GSK-3β | **Hymenidin** | **-7.24** | **4.92 µM** | Tyr216, Cys218, Gly262, Ser261, Ser219, Asp260, Arg220, Tyr221, Gln265, Gly259, Arg223 | 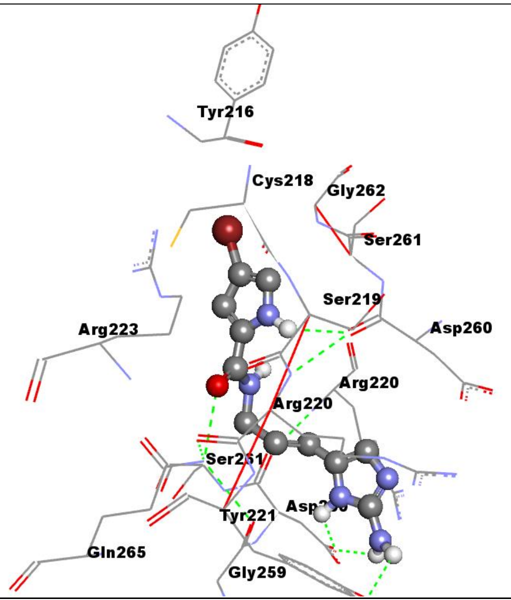 |
