## Supplementary material for "Natural compound screening predicts novel GSK-3 isoform-specific inhibitors": Suppl. Table 2

**Suppl. Table 2.** Bioactivity score (BAS) and Druglikeness of selected natural compounds versus reference synthetic drugs calculated by Molinspiration. BAS>0.0 (active), BAS -5.0- 0.0 (moderately active), BAS<-5.0 (inactive).

| S.No. | Ligand | GPCR ligand | Ion channel modulator | Kinase inhibitor | Nuclear receptor ligand | Protease inhibitor | Enzyme inhibitor | Druglikeness |
|---|---|---|---|---|---|---|---|---|
| 1 | Berberine | -0.11 | 0.71 | -0.27 | -0.78 | -0.35 | 0.82 | -2.2467 |
| 2 | **Rosmarinic acid** | **-0.11** | **0.71** | **-0.27** | **-0.78** | **-0.35** | **0.82** | **-3.8118** |
| 3 | Luteolin | -0.02 | -0.07 | 0.26 | 0.39 | -0.22 | 0.28 | 0.28194 |
| 4 | Apigenin | -0.07 | -0.09 | 0.18 | 0.34 | -0.25 | 0.26 | 0.28194 |
| 5 | Daidzein | -0.30 | -0.64 | -0.20 | 0.04 | -0.83 | 0.02 | -0.093853 |
| 6 | Galangin | -0.13 | -0.21 | 0.19 | 0.28 | -0.32 | 0.28 | -0.082832 |
| 7 | **Psoralidin** | **-0.20** | **-0.09** | **-0.17** | **0.53** | **-0.15** | **0.21** | **-0.53359** |
| 8 | Catechin | 0.41 | 0.14 | 0.09 | 0.60 | 0.26 | 0.47 | 0.31525 |
| 9 | Crocetin | 0.13 | 0.19 | -0.01 | 0.68 | -0.04 | 0.40 | 1.2391 |
| 10 | Piperine | 0.15 | -0.18 | -0.13 | -0.13 | -0.10 | 0.04 | 0.43286 |
| 11 | Sesamin | 0.02 | -0.31 | -0.27 | -0.09 | -0.15 | 0.03 | -1.0557 |
| 12 | Quercetin | -0.06 | -0.19 | 0.28 | 0.36 | -0.25 | 0.28 | -0.082832 |
| 13 | Resveratrol | -0.20 | 0.02 | -0.20 | 0.01 | -0.41 | 0.02 | -1.6732 |
| 14 | Kaempferol | -0.10 | -0.21 | 0.21 | 0.32 | -0.27 | 0.26 | -0.082832 |
| 15 | Costunolide | 0.24 | 0.07 | -0.52 | 0.96 | -0.28 | 0.77 | -9.162 |
| 16 | Withaferin A | 0.07 | 0.14 | -0.49 | 0.76 | 0.15 | 0.94 | 1.6889 |
| 17 | Serpentine | 0.21 | 0.35 | -0.07 | 0.05 | -0.06 | 0.16 | 2.3376 |
| 18 | MeBIO | -0.34 | 0.01 | 0.57 | -0.35 | -0.47 | -0.04 | -0.65189 |
| 19 | ALOISINE A | 0.15 | -0.07 | 0.58 | -0.09 | -0.34 | 0.28 | -4.945 |
| 20 | CHIR-99021 | 0.45 | 0.22 | 0.85 | -0.11 | 0.07 | 0.30 | -2.2161 |
| 21 | Azithromycin | -0.60 | -1.50 | -1.35 | -1.40 | -0.28 | -0.82 | 13.854 |
