## Supplementary material for "Natural compound screening predicts novel GSK-3 isoform-specific inhibitors": Suppl. Table 3

**Supplementary table 3.** Toxicity potential of selected natural compounds versus reference synthetic drugs calculated by Osiris Property Explorer.

| S.No. | Ligand name | Mutagenic | Tumorigenic | Irritant | Reproductive effect |
|---|---|---|---|---|---|
| 1 | Berberine | None | None | None | None |
| 2 | Rosmarinic acid | None | None | None | None |
| 3 | Luteolin | None | None | None | None |
| 4 | Apigenin | High | None | None | None |
| 5 | Daidzein | None | None | None | High |
| 6 | Galangin | High | None | None | None |
| 7 | Psoralidin | None | None | None | High |
| 8 | Catechin | None | None | None | None |
| 9 | Crocetin | None | None | None | None |
| 10 | Piperine | None | None | None | High |
| 11 | Sesamin | None | None | None | None |
| 12 | Quercetin | High | High | None | None |
| 13 | Resveratrol | High | None | None | High |
| 14 | Kaempferol | High | None | None | None |
| 15 | Costunolide | None | None | High | None |
| 16 | Withaferin A | None | None | None | Low |
| 17 | Serpentine | None | None | None | None |
| 18 | MeBIO | High | High | None | None |
| 19 | ALOISINE A | None | None | None | None |
| 20 | CHIR-99021 | None | None | None | None |
| 21 | Azithromycin | None | None | None | None |

High Risk

None Risk

Mild Risk
