## Supplementary material for "Natural compound screening predicts novel GSK-3 isoform-specific inhibitors": Suppl. Table 5

**Supplementary table 5.** Abbot Bioavailability Score of selected natural compounds versus reference synthetic drugs calculated using SwissADME.

| S.No. | Compounds | GI absorption (Gatrointestinal Absorption) | BBB permeant | P-gp substrate (P-glycoprotein substrate) | CYP1A2 inhibitor (Cytochrome P450 1A2 Inhibitor) | CYP2C19 inhibitor (Cytochrome P450 2C19 Inhibitor) | CYP2C9 inhibitor (Cytochrome P450 2C9 Inhibitor) | CYP2D6 inhibitor (Cytochrome P450 2D6 Inhibitor) | CYP3A4 inhibitor (Cytochrome P450 3A4 Inhibitor) | Log Kp (cm/s) (skin permeation) |
|---|---|---|---|---|---|---|---|---|---|---|
| 1 | Berberine | High | Yes | Yes | Yes | No | No | Yes | Yes | -5.78 |
| 2 | Rosmarinic acid | Low | No | No | No | No | No | No | No | -6.82 |
| 3 | Luteolin | High | No | No | Yes | No | No | Yes | Yes | -6.25 |
| 4 | Apigenin | High | No | No | Yes | No | No | Yes | Yes | -5.80 |
| 5 | Daidzein | High | Yes | No | Yes | No | No | Yes | Yes | -6.10 |
| 6 | Galangin | High | No | No | Yes | No | No | Yes | Yes | -6.35 |
| 7 | Psoralidin | High | No | No | Yes | Yes | Yes | No | No | -5.02 |
| 8 | Catechin | High | No | Yes | No | No | No | No | No | -7.82 |
| 9 | Crocetin | High | No | No | No | Yes | Yes | No | No | -4.46 |
| 10 | Piperine | High | Yes | No | Yes | Yes | Yes | No | No | -5.58 |
| 11 | Sesamin | High | Yes | No | No | Yes | No | Yes | Yes | -6.56 |
| 12 | Quercetin | High | No | No | Yes | No | No | Yes | Yes | -7.05 |
| 13 | Resveratrol | High | Yes | No | No | No | No | No | Yes | -5.47 |
| 14 | Kaempferol | High | No | No | Yes | No | No | Yes | Yes | -6.70 |
| 15 | Costunolide | High | Yes | No | No | Yes | Yes | No | No | -6.23 |
| 16 | Withaferin A | High | No | Yes | No | No | No | No | No | -6.45 |
| 17 | Serpentine | High | Yes | No | No | Yes | No | Yes | Yes | -6.59 |
| 18 | MeBIO | High | No | No | Yes | Yes | Yes | Yes | No | -5.84 |
| 19 | ALOISINE A | High | Yes | Yes | Yes | Yes | No | Yes | Yes | -5.43 |
| 20 | CHIR-99021 | High | No | No | Yes | Yes | Yes | Yes | Yes | -6.09 |
| 21 | Azithromycin | Low | No | Yes | No | No | No | No | No | -8.01 |
