## Supplementary material for "Natural compound screening predicts novel GSK-3 isoform-specific inhibitors": Suppl. Table 6

**Supplementary Table 6:** Bioavailability radar graphs of natural compounds vs. reference drugs.

| S.No. | Compounds | LIPO | SIZE | POLAR | INSOLU | INSATU | FLEX |
|---|---|---|---|---|---|---|---|
| 1 | Berberine | 3.62 | 336.36 g/mol | 40.80 $Å^2$ | -4.55 | 0.25 | 2 |
| 2 | Rosmarinic acid | 2.36 | 360.31 g/mol | 144.52 $Å^2$ | -3.44 | 0.11 | 7 |
| 3 | Luteolin | 2.53 | 286.24 g/mol | 111.13 $Å^2$ | -3.71 | 0.00 | 1 |
| 4 | Apigenin | 3.02 | 270.24 g/mol | 90.90 $Å^2$ | -3.94 | 0.00 | 1 |
| 5 | Daidzein | 2.47 | 254.24 g/mol | 70.67 $Å^2$ | -3.53 | 0.00 | 1 |
| 6 | Galangin | 2.25 | 270.24 g/mol | 90.90 $Å^2$ | -3.46 | 0.00 | 1 |
| 7 | Psoralidin | 4.69 | 336.34 g/mol | 83.81 $Å^2$ | -5.25 | 0.15 | 2 |
| 8 | Catechin | 0.36 | 290.27 g/mol | 110.38 $Å^2$ | -2.22 | 0.20 | 1 |
| 9 | Crocetin | 5.41 | 328.40 g/mol | 74.60 $Å^2$ | -4.76 | 0.20 | 8 |
| 10 | Piperine | 3.46 | 285.34 g/mol | 38.77 $Å^2$ | -3.74 | 0.35 | 4 |
| 11 | Sesamin | 2.68 | 354.35 g/mol | 55.38 $Å^2$ | -3.93 | 0.40 | 2 |
| 12 | Quercetin | 1.54 | 302.24 g/mol | 131.36 $Å^2$ | -3.16 | 0.00 | 1 |
| 13 | Resveratrol | 3.13 | 228.24 g/mol | 60.69 $Å^2$ | -3.62 | 0.00 | 2 |
| 14 | Kaempferol | 1.90 | 286.24 g/mol | 111.13 $Å^2$ | -3.31 | 0.00 | 1 |
| 15 | Costunolide | 2.09 | 232.32 g/mol | 26.30 $Å^2$ | -2.60 | 0.53 | 0 |
| 16 | Withaferin A | 3.83 | 470.60 g/mol | 96.36 $Å^2$ | -4.97 | 0.79 | 3 |
| 17 | Serpentine | 2.58 | 348.40 g/mol | 53.35 $Å^2$ | -3.86 | 0.33 | 2 |
| 18 | MeBIO | 3.83 | 370.20 g/mol | 70.38 $Å^2$ | -5.00 | 0.06 | 2 |
| 19 | ALOISINE A | 3.52 | 267.33 g/mol | 61.80 $Å^2$ | -4.01 | 0.25 | 4 |

| | | | | | | | |
|---|---|---|---|---|---|---|---|
| 20 | CHIR-99021 | 4.30 | 465.34 g/mol | 115.20 Å$^2$ | -5.50 | 0.14 | 7 |
| 21 | Azithromycin | 4.02 | 748.98 g/mol | 180.08 Å$^2$ | -6.55 | 0.97 | 7 |

**Note:**

**LIPO (Lipophility):** -0.7 < XLOGP3 < + 5.0
**SIZE:** 150 g/mol < MW <500 g/mol
**POLAR (Polarity):** 20 Å$^2$ < TPSA < 130 Å$^2$
**INSOLU (Insolubility):** -6 < Log S (ESOL) <0
**INSATU (Instauration):** 0.25 < Fraction Csp3 < 1
**FLEX (Flexibility):** 0 < No. rotatable bonds < 9
The colored space is the suitable physicochemical space for oral bioavailability
