## Supplementary material for "Natural compound screening predicts novel GSK-3 isoform-specific inhibitors": Suppl. Figures

**Supplementary Figure 1;** Homology modeled GSK-3α protein structure (SwissADME software).

| Model #01 | File | Built with | Oligo-State | Ligands | GMQE | QMEANDisCo Global |
|---|---|---|---|---|---|---|
| 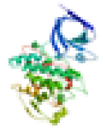 | PDB | ProMod3 3.2.0 | monomer | 1 x MLA: MALONIC ACID; | 0.70 | 0.87 ± 0.05 |

| Template | Seq Identity | Oligo-state | QSQE | Found by | Method | Resolution | Seq Similarity | Range | Coverage | Description |
|---|---|---|---|---|---|---|---|---|---|---|
| 3say.1.A | 82.97 | monomer | 0.00 | BLAST | X-ray | 2.23Å | 0.56 | 98 - 447 | 0.77 | Glycogen synthase kinase-3 beta |

**Included Ligands**

| Ligand | Description |
|---|---|
| 1 x MLA | MALONIC ACID |

**Excluded ligands**

| Ligand Name.Number | Reason for Exclusion | Description |
|---|---|---|
| FMT.3 | Not in contact with model. | FORMIC ACID |
| OFT.1 | Binding site not conserved. | (3Z)-N,N-DIETHYL-3-[(3E)-3-(HYDROXYIMINO)-1,3-DIHYDRO-2H-INDOL-2-YLIDENE]-2-OXO-2,3-DIHYDRO-1H-INDOLE-5-SULFONAMIDE |

**Supplementary Figure 2;** MolProbity analysis of modeled GSK-3α protein structure.

#### Summary statistics

| | | | | |
|---|---|---|---|---|
| All-Atom Contacts | Clashscore, all atoms: | 1.24 | | $99^{th}$ percentile* (N=1784, all resolutions) |
| | Clashscore is the number of serious steric overlaps (> 0.4 Å) per 1000 atoms. | | | |
| Protein Geometry | Poor rotamers | 11 | 3.54% | Goal: <0.3% |
| | Favored rotamers | 292 | 93.89% | Goal: >98% |
| | Ramachandran outliers | 2 | 0.57% | Goal: <0.05% |
| | Ramachandran favored | 333 | 95.69% | Goal: >98% |
| | Rama distribution Z-score | -0.11 ± 0.42 | | Goal: abs(Z score) < 2 |
| | MolProbity score^ | 1.56 | | $94^{th}$ percentile* (N=27675, 0Å - 99Å) |
| | Cβ deviations >0.25Å | 5 | 1.51% | Goal: 0 |
| | Bad bonds: | 0 / 2864 | 0.00% | Goal: 0% |
| | Bad angles: | 31 / 3890 | 0.80% | Goal: <0.1% |
| Peptide Omegas | Cis Prolines: | 0 / 24 | 0.00% | Expected: ≤1 per chain, or ≤5% |
| Low-resolution Criteria | CaBLAM outliers | 2 | 0.6% | Goal: <1.0% |
| | CA Geometry outliers | 1 | 0.29% | Goal: <0.5% |
| Additional validations | Chiral volume outliers | 0/439 | | |
| | Waters with clashes | 0/0 | 0.00% | See UnDowser table for details |

In the two column results, the left column gives the raw count, right column gives the percentage.

* $100^{th}$ percentile is the best among structures of comparable resolution; $0^{th}$ percentile is the worst. For clashscore the comparative set of structures was selected in 2004, for MolProbity score in 2006.

^ MolProbity score combines the clashscore, rotamer, and Ramachandran evaluations into a single score, normalized to be on the same scale as X-ray resolution.

Key to table colors and cutoffs here:

**Supplementary Figure 3;** Molprobity Ramachandran analysis of GSK-3α homology model.

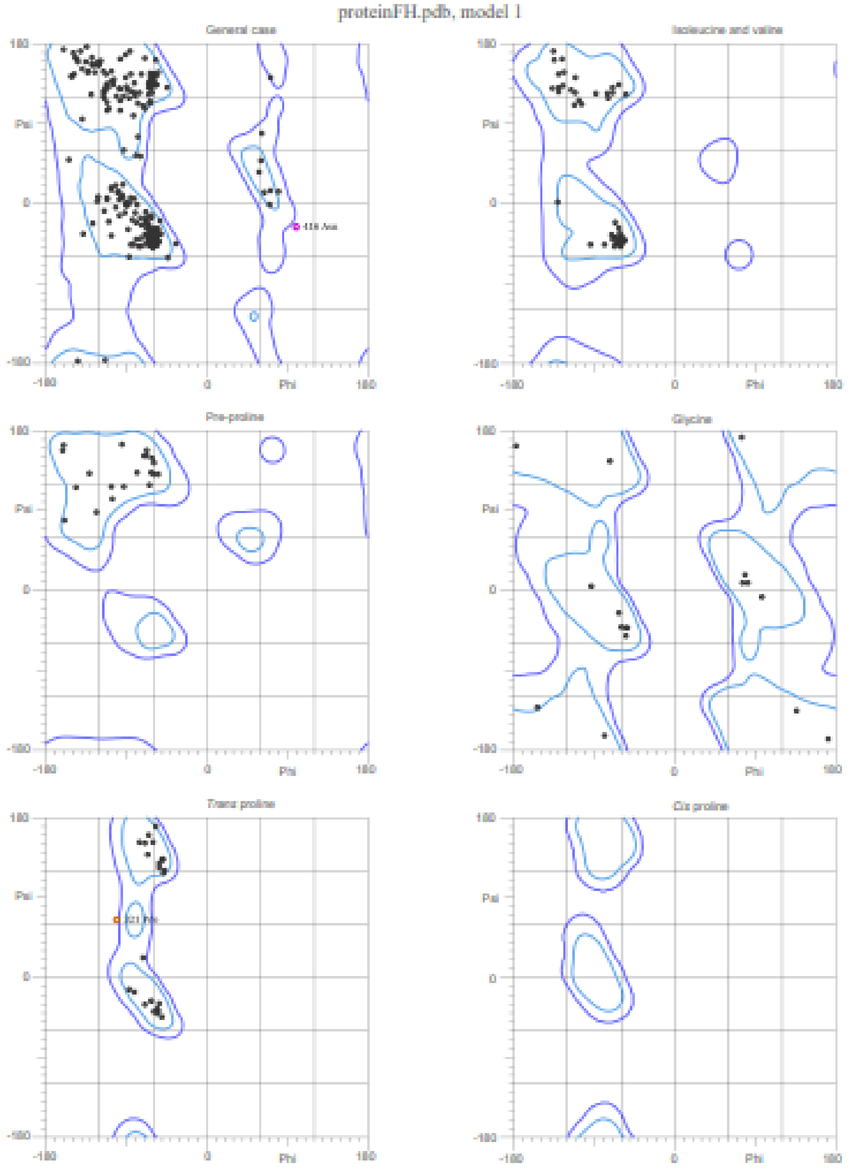


95.7% (333/348) of all residues were in favored (98%) regions.
99.4% (346/348) of all residues were in allowed (>99.8%) regions.

There were 2 outliers (phi, psi):
321 Pro (-102.8, 66.9)
416 Asn (101.6, -26.5)

**Supplementary Figure 4;** Chemical structures of included natural compounds.

1

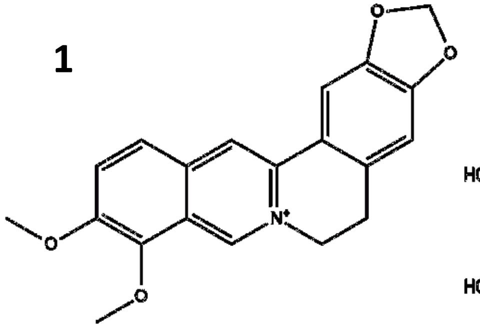

Berberine

2

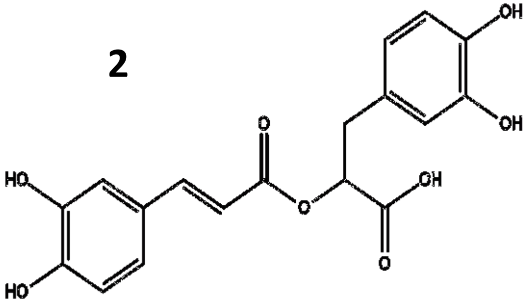

Rosmarinic acid

3

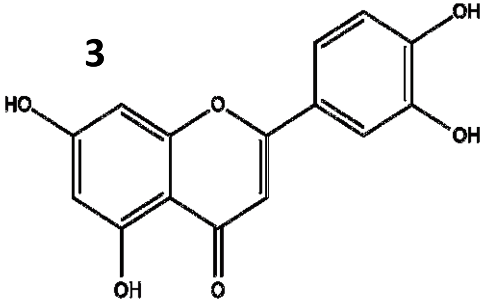

Luteolin

4

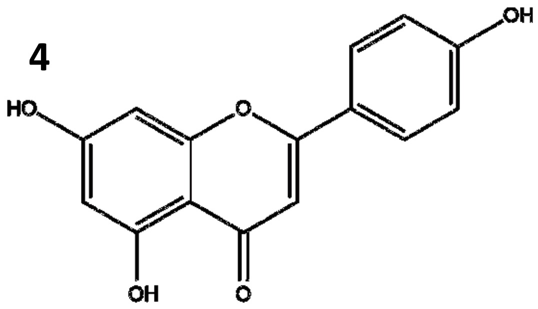

Apigenin

5

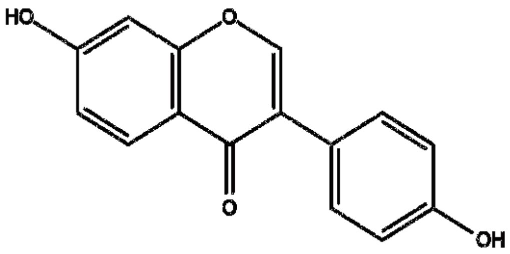

Daidzein

6

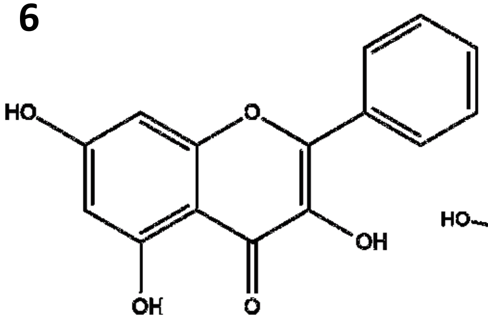

Galangin

7

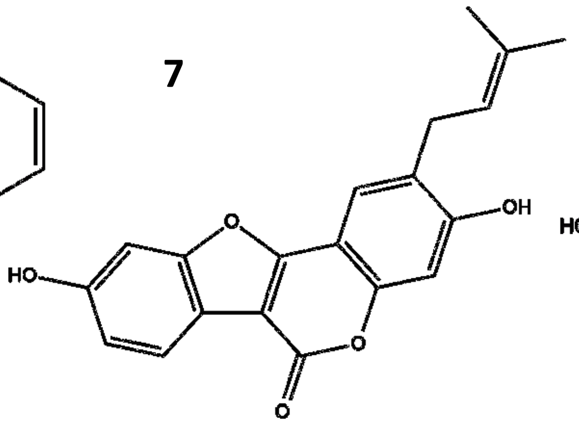

Psoralidin

8

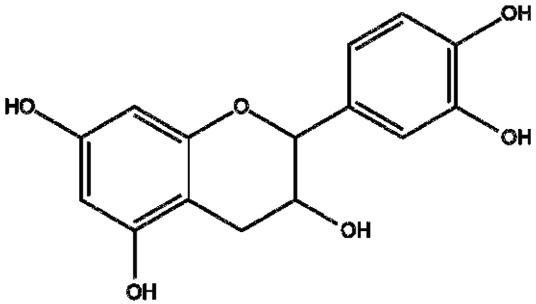

Catechin

9

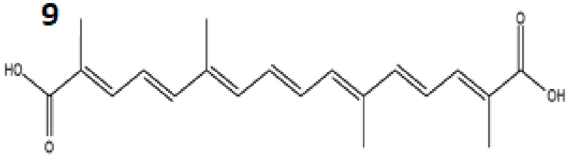

Crocetin

10

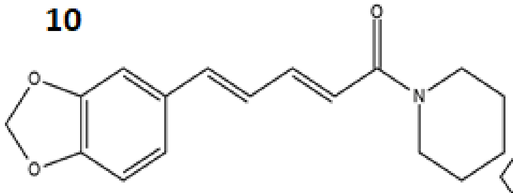

Piperine

11

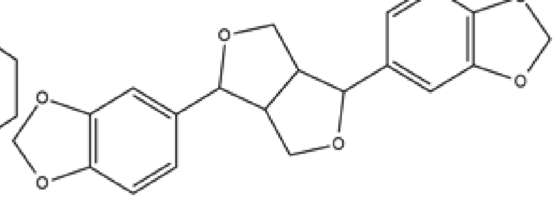

Sesamin

12

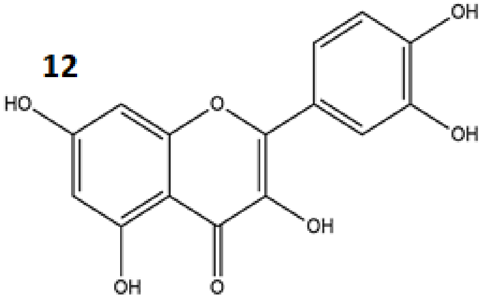

Quercetin

**Supplementary Figure 4 cont..;** Chemical structures of included natural compounds.

**13**

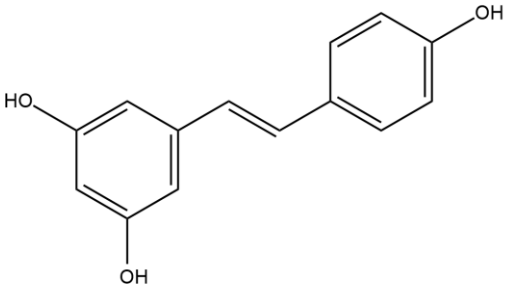

**Resveratrol**

**14**

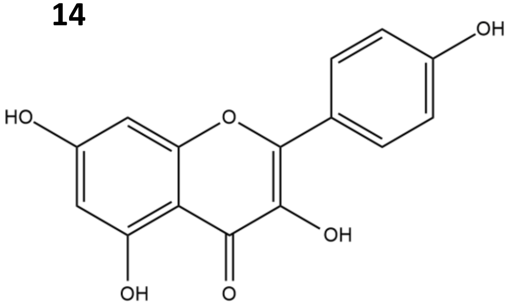

**Kaempferol**

**15**

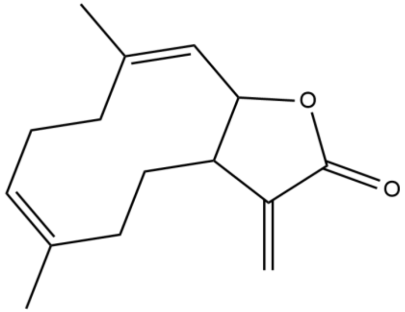

**Costunolide**

**16**

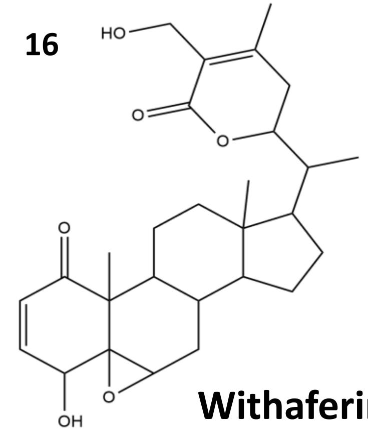

**Withaferin A**

**17**

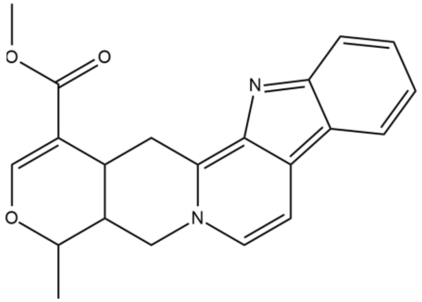

**Serpentine**

18

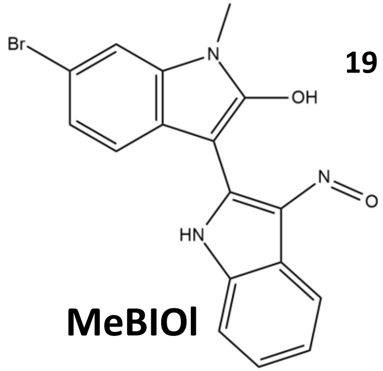

**MeBIOl**

**19**

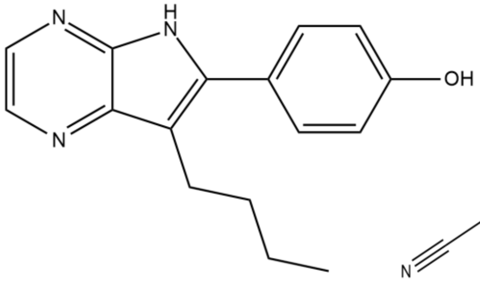

**Aloisine A**

**20**

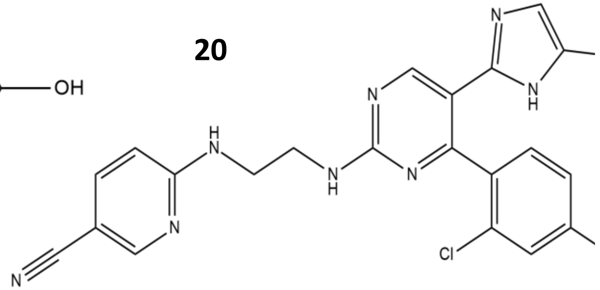

**Chir-99021**

**21**

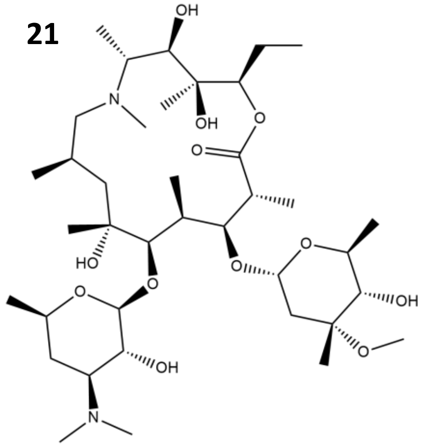

**Azithromycin**

### Supplementary Figure 5

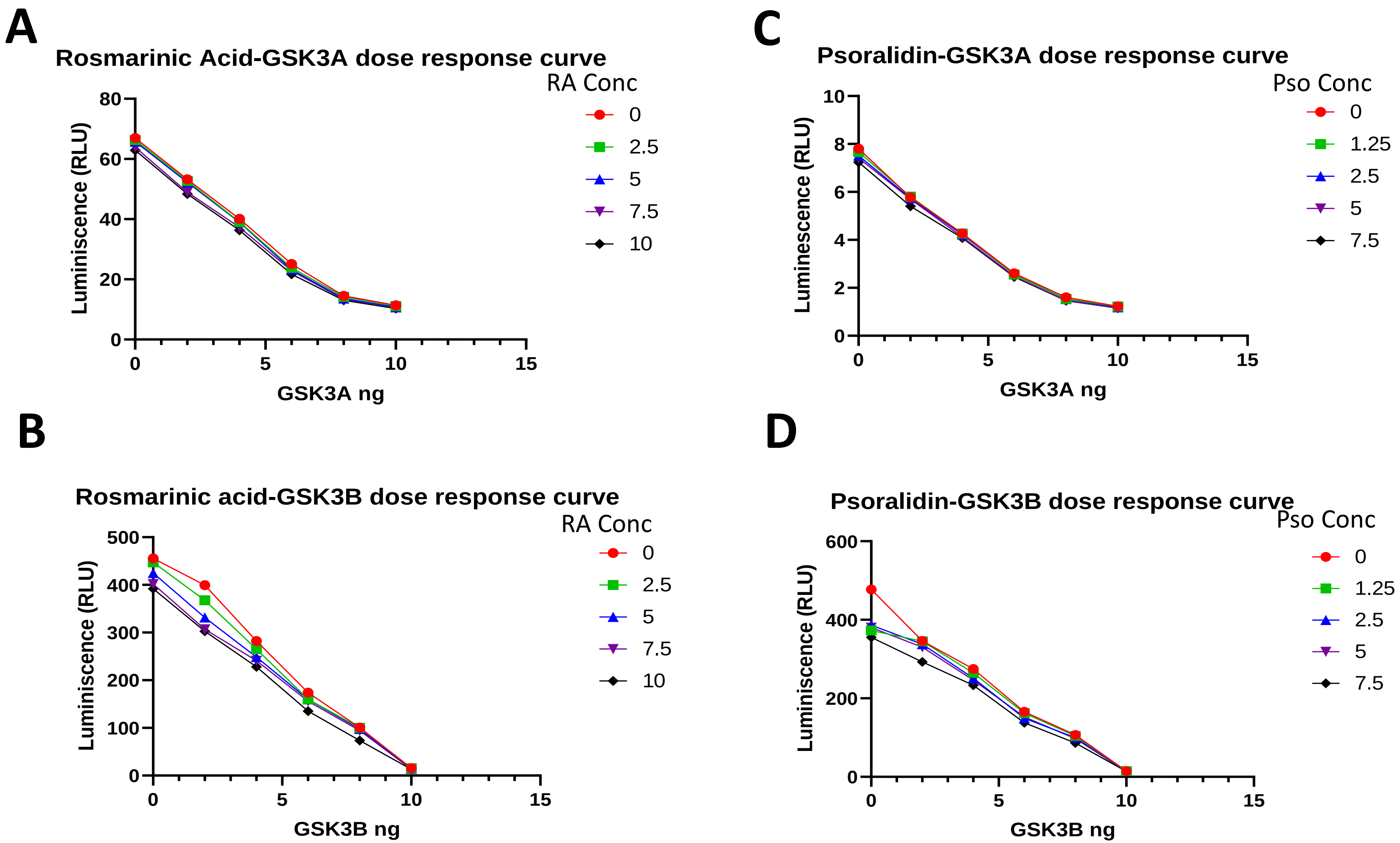
